# Single-cell DNA Methylome and 3D Multi-omic Atlas of the Adult Mouse Brain

**DOI:** 10.1101/2023.04.16.536509

**Authors:** Hanqing Liu, Qiurui Zeng, Jingtian Zhou, Anna Bartlett, Bang-An Wang, Peter Berube, Wei Tian, Mia Kenworthy, Jordan Altshul, Joseph R. Nery, Huaming Chen, Rosa G. Castanon, Songpeng Zu, Yang Eric Li, Jacinta Lucero, Julia K. Osteen, Antonio Pinto-Duarte, Jasper Lee, Jon Rink, Silvia Cho, Nora Emerson, Michael Nunn, Carolyn O’Connor, Zizhen Yao, Kimberly A. Smith, Bosiljka Tasic, Hongkui Zeng, Chongyuan Luo, Jesse R. Dixon, Bing Ren, M. Margarita Behrens, Joseph R Ecker

**Affiliations:** Genomic Analysis Laboratory, The Salk Institute for Biological Studies, La Jolla, CA, USA.; Division of Biological Sciences, University of California, San Diego, La Jolla, CA, USA.; Bioinformatics and Systems Biology Program, University of California, San Diego, La Jolla, CA, USA.; Department of Cellular and Molecular Medicine, University of California, San Diego School of Medicine, La Jolla, CA, USA.; Computational Neurobiology Laboratory, The Salk Institute for Biological Studies, La Jolla, CA, USA.; Flow Cytometry Core Facility, The Salk Institute for Biological Studies, La Jolla, CA, USA.; Allen Institute for Brain Science, Seattle, WA, USA.; Department of Human Genetics, University of California Los Angeles, Los Angeles, CA, USA.; Peptide Biology Laboratory, The Salk Institute for Biological Studies, La Jolla, CA, USA.; Center for Epigenomics, University of California, San Diego School of Medicine, La Jolla, CA, USA.; Institute of Genomic Medicine, University of California, San Diego School of Medicine, La Jolla, CA, USA.; Howard Hughes Medical Institute, The Salk Institute for Biological Studies, La Jolla, CA, USA.

## Abstract

Cytosine DNA methylation is essential in brain development and has been implicated in various neurological disorders. A comprehensive understanding of DNA methylation diversity across the entire brain in the context of the brain’s 3D spatial organization is essential for building a complete molecular atlas of brain cell types and understanding their gene regulatory landscapes. To this end, we employed optimized single-nucleus methylome (snmC-seq3) and multi-omic (snm3C-seq^1^) sequencing technologies to generate 301,626 methylomes and 176,003 chromatin conformation/methylome joint profiles from 117 dissected regions throughout the adult mouse brain. Using iterative clustering and integrating with companion whole-brain transcriptome and chromatin accessibility datasets, we constructed a methylation-based cell type taxonomy that contains 4,673 cell groups and 261 cross-modality-annotated subclasses. We identified millions of differentially methylated regions (DMRs) across the genome, representing potential gene regulation elements. Notably, we observed spatial cytosine methylation patterns on both genes and regulatory elements in cell types within and across brain regions. Brain-wide multiplexed error-robust fluorescence in situ hybridization (MERFISH^2^) data validated the association of this spatial epigenetic diversity with transcription and allowed the mapping of the DNA methylation and topology information into anatomical structures more precisely than our dissections. Furthermore, multi-scale chromatin conformation diversities occur in important neuronal genes, highly associated with DNA methylation and transcription changes. Brain-wide cell type comparison allowed us to build a regulatory model for each gene, linking transcription factors, DMRs, chromatin contacts, and downstream genes to establish regulatory networks. Finally, intragenic DNA methylation and chromatin conformation patterns predicted alternative gene isoform expression observed in a companion whole-brain SMART-seq^3^ dataset. Our study establishes the first brain-wide, single-cell resolution DNA methylome and 3D multi-omic atlas, providing an unparalleled resource for comprehending the mouse brain’s cellular-spatial and regulatory genome diversity.

## Introduction

The mouse brain is a complex organ composed of millions of cells forming hundreds of anatomical structures that exhibit diverse cell types^4–9^. This cellular and spatial diversity is stringently controlled by epigenetic mechanisms^10–12^. Cytosine DNA methylation (5mC) is a stable covalent modification that endures in post-mitotic cells for their entire lifespan^13^. This modification is associated with neuronal function, behavior, and various diseases^14^. In mammalian genomes, 5mC occurs predominantly at CpG sites (mCG). However, in neurons, non-CpG cytosine methylation (mCH, H denotes A, C, or T) is also abundant^13, 15^. Both methylation forms directly influence the DNA-binding of methyl CpG binding protein 2 (MeCP2)^16–, 19^, a critical 5mC reader and the cause of Rett syndrome^20^. Methylation of CpG and CpH dynamically occurs at regulatory elements and gene bodies with cellular and spatial diversity, modulating transcription factor binding affinity and controlling gene transcription^21^. Genome-wide differential methylation analysis can predict millions of regulatory elements^10, 22^, yielding a cellular taxonomy and a base-resolution genome atlas. Furthermore, chromatin conformation connects these regulatory elements to their target genes, providing a comprehensive view of the gene’s regulatory environment^23^. Intragenic DNA methylation displays hundred-kilobase-level patterns that coincide with genome topological features, indicating that both epigenetic modalities collaboratively regulate precise gene expression^24^.

This study uses an optimized version of single-nucleus methylation sequencing (snmC-seq3) to analyze the DNA methylome at single-cell resolution^25, 26^, and single-nucleus methylation and chromatin conformation capture sequencing (snm3C-seq)^1^ to jointly investigate the two modalities. Alongside the previous study^10^, we collect 301,626 methylomes and 176,003 m3C joint profiles from the entire mouse brain. This ultra-deep base-resolution dataset of the mouse regulatory genome comprises 786 billion final methylation reads (snmC-seq3 + snm3C-seq) and 33 billion cis-long-range chromatin contacts (snm3C-seq), with each sequencing fragment assigned to individual cells. We define the cell type taxonomy of the whole mouse brain based on DNA methylome, providing 4,673 cell clusters/spatial groups. Subsequently, we demonstrate that methylome taxonomy accurately aligns with other molecular modalities from the BRAIN Initiative Cell Census Network (BICCN) at the cluster level^7, 12^. Through integration analysis, we annotate the methylome clusters using nomenclature from the transcriptomic studies, offering a comprehensive multi-omic resource for the field.

Additionally, we observe the prevalence of spatial information in the epigenome, which was validated with a MERFISH dataset generated with spatial methyl-diverse genes. Integration with spatial transcriptomics allows us to map the DNA methylome and chromatin conformation data onto fine anatomical structures, confirming that the epigenetic spatial pattern corresponds with transcription differences in vivo.

We also investigate the regulatory landscapes of individual genes by comparing thousands of cluster-aggregated epigenetic profiles. We observe chromatin compartment diversity across brain cell types in brain development-related gene bodies, and the compartment score is negatively correlated with methylation fraction. Furthermore, in the long genes with critical neuronal and synaptic functions, we observe elevated domain boundary formation that correlates with gene body methylation, indicating a gene-activity-related chromatin conformation change potentially cooperating with DNA methylation and transcription machinery. Finally, we provide a chromatin conformation landscape for individual genes through unbiased variance and correlation analysis of the chromatin interaction. Intersecting this landscape with correlation-based analysis, we build gene regulatory networks linking transcription factors, DMRs, and target genes. We also identify numerous validated and novel regulatory relationships of critical neuronal TFs, including many immediate early genes (IEGs).

Finally, base-resolution DNA methylation profiles in many long genes reveal multiple cell-type-specific patterns corresponding to the heterogeneity of alternative gene transcription start sites (TSS) and exon usage within the same gene. We systematically investigate this phenomenon by integrating single nucleus methylation data with a companion whole-brain full-length single-cell RNA-seq (SMART-seq v4^3^) dataset generated by Allen Institute for Brain Science (AIBS)^7^. By quantifying the gene expression at the transcript level^27^, we observe cell-type-specific alternative TSS and exon usage in many long neuronal and synaptic genes, whose gene body methylation and chromatin conformation patterns can predict isoform diversity.

## Results

### Single-cell DNA methylome and chromatin conformation atlas of the entire mouse brain

We developed snmC-seq3, an optimized single-nucleus methylome sequencing method (Supplementary Information 1), to profile genome-wide 5mC at base resolution (Fig. 1a) across 117 dissection regions in the whole brain from adult male C57BL/6 mice (Fig. 1b, Extended Data Fig. 1a, Supplementary Table 1). Additionally, we employed snm3C-seq, a multi-omic technology^1^, to profile the DNA methylome and chromatin conformation jointly from 33 dissection regions (Extended Data Fig. 1b), thus adding the 3D genome context across all brain cell types (Fig. 1a). Each dissected region is represented by 2-3 replicates, obtained from pooling the same region from at least six animals. Single nuclei were captured through fluorescence-activated nuclei sorting (FANS), enriching NeuN-antibody positively labeled neurons (NeuN+ comprised 92% of snmC and 78% of snm3C, with remaining data being NeuN- or non-neurons, Methods). Collectively, including previous research^10^, we obtained 324,687 (301,626 passed QC) DNA methylome profiles. On average, the snmC-seq dataset had 1.44±0.50 million (mean±s.d.) final reads, covering 72±24 million (6.5%±2.2%) of cytosine bases in the mouse genome. We also obtained 196,172 (176,003 passed QC) joint methylome and chromatin conformation capture (3C) profiles, with each cell having 1.99±0.57 million final reads, covering 72±20 million (6.5%±1.8%) of cytosine bases. The 3C modality of each cell had 188±81 thousand (18.3%±5.7% of the total fragments) cis-long-range contacts and 108±41 (10.4%±2.3%) thousand trans-contacts (Extended Data Fig. 2, Methods, Supplementary Table 2, 3).

**Figure 1.**
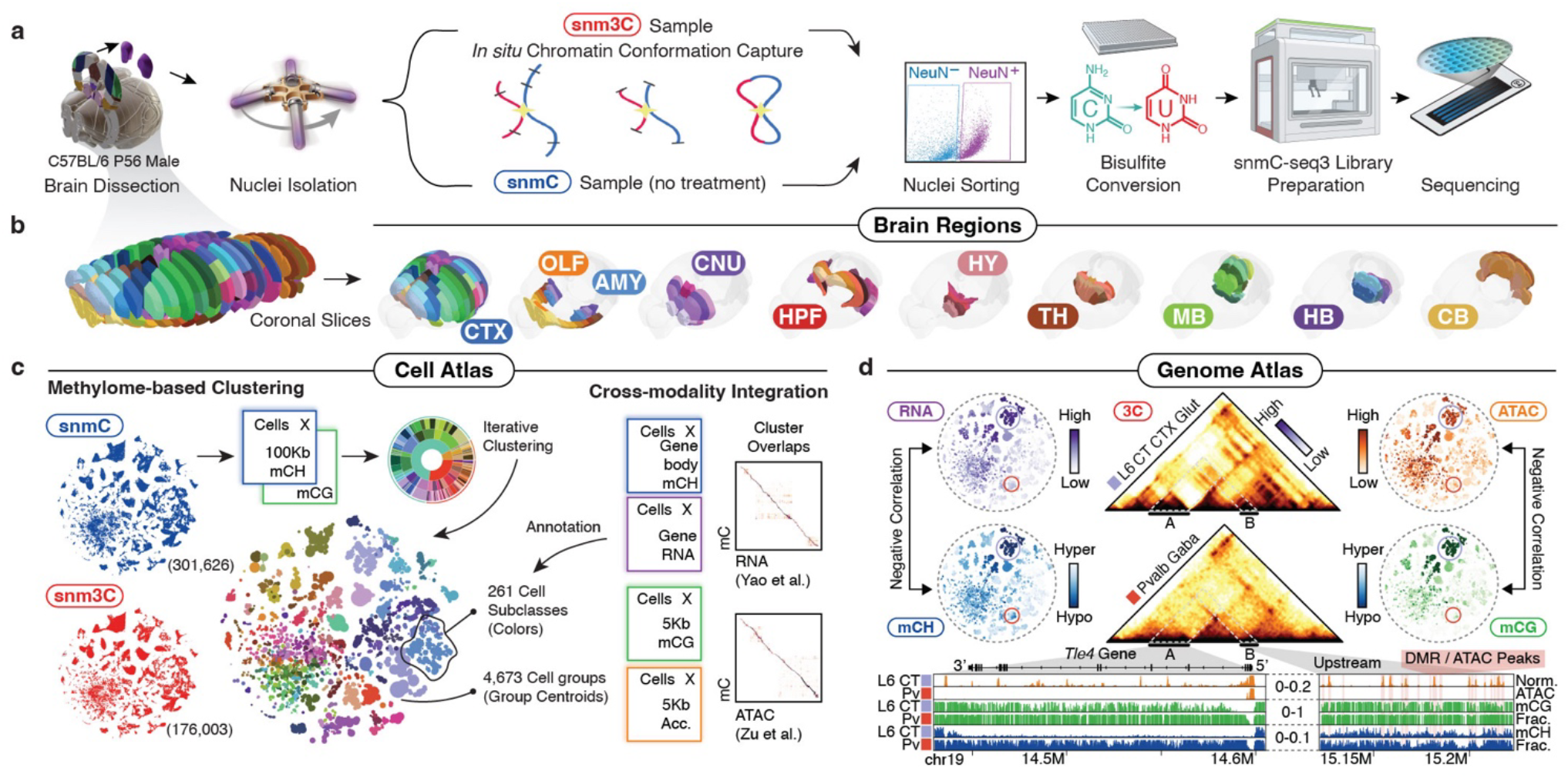
Single-cell DNA methylome and multi-omic atlas chart the cellular and genomic diversity of the whole mouse brain. **a,** The workflow of dissection, nuclei, and library preparation for snmC-seq3 and snm3C-seq. **b,** The 117 dissection regions from eighteen 600-μm coronal slices are grouped into ten major brain regions (see Supplementary Table 9 for abbreviations). Each dissection region is registered to the 3D common coordinate framework (CCF)^4^. **c,** The cell atlas: methylome-based iterative clustering on snmC and snm3C datasets. The left t-SNE plot is colored by modality; the middle plot is aggregated into 4,673 cell group centroids and colored by 261 cell subclasses; The right part demonstrates cross-modality integration of brain-wide datasets from BICCN, details in Figure 2. **d,** The genome atlas: the *Tle4* gene exemplifies pseudo-bulk profiles of five modalities across the whole brain, with genome browser view of the “L6 CT CTX Glut” and “Pvalb Gaba” subclasses in the bottom.

**Figure 2.**
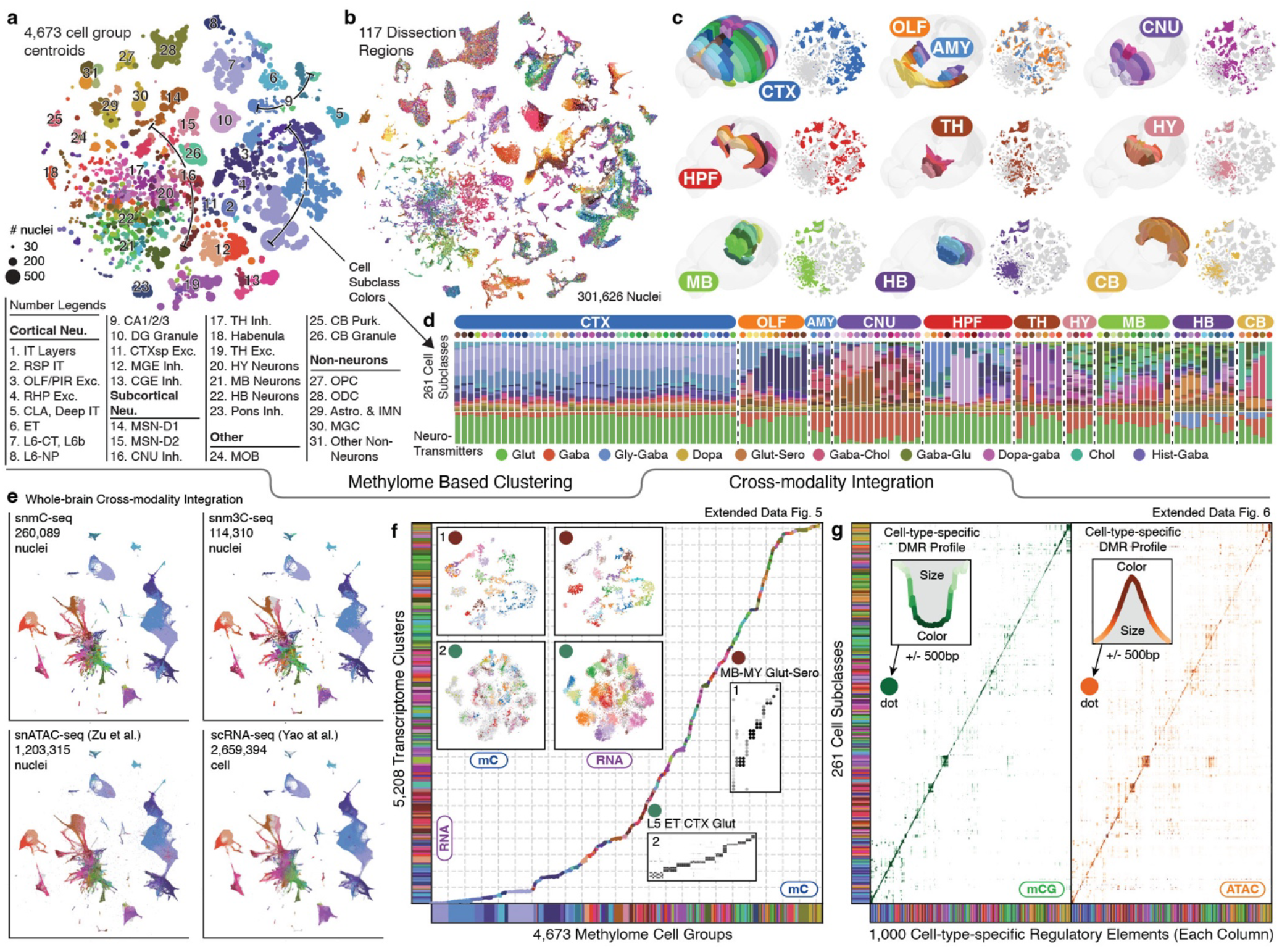
Consensus cell type taxonomy across molecular modalities. **a,** Cell-group-centroids t-SNE color by cell subclass (see Extended Data Fig. 3 for number legends’ abbreviations). **b,** Cell-level t-SNE color by 117 dissection regions. **c,** 3D CCF registration and cell t-SNE of each major region. **d,** Cell subclass (upper row) and neurotransmitter composition (bottom row) of each brain dissection region (each upper dot), grouped by major region. **e,** Integration t-SNE of all neurons from the snmC-seq, snm3C-seq, snATAC-seq, and scRNA-seq datasets, colored by matched cell subclasses. **f,** Brain-wide cluster map between the snmC-seq and scRNA-seq datasets (Supplementary Table 4) based on iterative integration. Each dot, colored by subclasses, on the diagonal represents a link between the mC clusters (x-axis) and RNA clusters (y-axis). Two examples in floating panels demonstrate highly granular correspondence of cell clusters in the final integration round: Box 1 presents the integration t-SNE colored by intra-modality clusters and confusion matrix of overlap score between “MB-MY Glut-Sero” clusters; Box 2 displays the same information for “L5 ET CTX Glut” clusters. See Extended Data Fig. 5 for more gene details. **g,** Dot plots of mCG fraction (left) and chromatin accessibility (right) of cell-type-specific CG-DMRs (columns) in each cell subclass (row). The size and color of each dot represent an aggregated epigenetic profile of 1,000 DMRs in a cell subclass; larger dot size and deeper color indicate these DMRs are more hypo-methylated or accessible in a subclass. See Extended Data Fig. 6 for more mC-ATAC integration details.

After quality control and preprocessing, we analyzed the data in cellular and genomic contexts (Fig. 1c, d). During the cellular analysis, we conducted iterative clustering of the mCH and mCG profiles in 100-Kb bins throughout the genome to establish a methylome-based whole-brain cell type taxonomy. At the highest level of granularity, we obtained a total of 4,376 cell clusters-by-spatial groups (Methods). To validate and annotate the dataset, we integrated the methylome data with other brain-wide chromatin accessibility^12^ and transcriptome datasets^7^, resulting in cluster-level mapping across modalities and annotations of these clusters into 261 subclass labels shared with companion transcriptome studies (Supplementary Table 4 and later sections).

Based on the clustering and integrative annotations, we produced pseudo-bulk profiles of five modalities (gene mCH, DMR mCG, chromatin conformation, accessibility, and gene expression) for each cell group, providing a cell-type-specific, multi-omic atlas for the mouse genome (Fig. 1d). With more details covered in later sections, we use the TLE family member 4 (*Tle4*) gene, a marker for the “L6 CT CTX Glut” subclass, as an example to illustrate the power of this comprehensive dataset. In this instance, cells with a low mCH fraction in the *Tle4* gene body exhibit high RNA expression levels (Fig. 1d, left), achieving a strong negative correlation across 4,673 cell groups (Pearson correlation coefficient, PCC. -0.86, p-value < 10^-15^). Similarly, the mCG fraction of an example DMR located in the *Tle4*-upstream region is also negatively correlated with chromatin accessibility signals (PCC. -0.73, p-value < 10^-15^, Fig. 1d, right). Furthermore, the m3C dataset provides chromatin contact information, indicating physical proximity between DMRs and *Tle4* gene in the “L6 CT CTX Glut” compared to the “Pvalb Gaba” subclass, where *Tle4* expression is low (Fig. 1d, middle). The base-resolution methylation profiles further reveal intricate cell-type specific epigenomic patterns, which offer rich information about the precise control of gene expression and transcript isoforms (Fig. 1d, bottom, and later section).

Overall, our study utilizes brain-wide single-cell mC and m3C datasets to (1) define cellular taxonomy based on the DNA methylome; (2) integrate with other atlas-level datasets from the BICCN, and (3) generate a multi-omic cell-type-specific genome atlas for the mouse brain. This unique resource allowed us to conduct several unprecedented analyses and make various discoveries, as we will describe in the upcoming sections.

### A methylome-based cell-type taxonomy

Following stringent quality control during preprocessing steps (Methods), we employed iterative clustering to classify methylome-based cell populations in the snmC-seq and snm3C-seq datasets, utilizing mCH and mCG profiles in 100-kb bins across the genome^10, 25^. In the final iteration, we identified 2,573 clusters and further separated the clusters based on brain dissection region into 4,673 cluster-by-spatial groups, which we used as the finest level of granularity for subsequent analyses (Fig. 2a, Extended Data Fig. 3). To establish a hierarchical structure for whole-brain cell types and support multi-omic data analysis, we iteratively integrated the methylome datasets with a companion brain-wide single-cell transcriptome dataset (see next section). After the integration, we annotated the mC-based cell groups in agreement with 261 RNA-based subclasses^7^ (Supplementary Table 4). The subsequent analyses rely on both the cluster-by-spatial group and subclass levels of cell classifications (Fig. 2a, Extended Data Fig. 3a,d).

**Figure 3.**
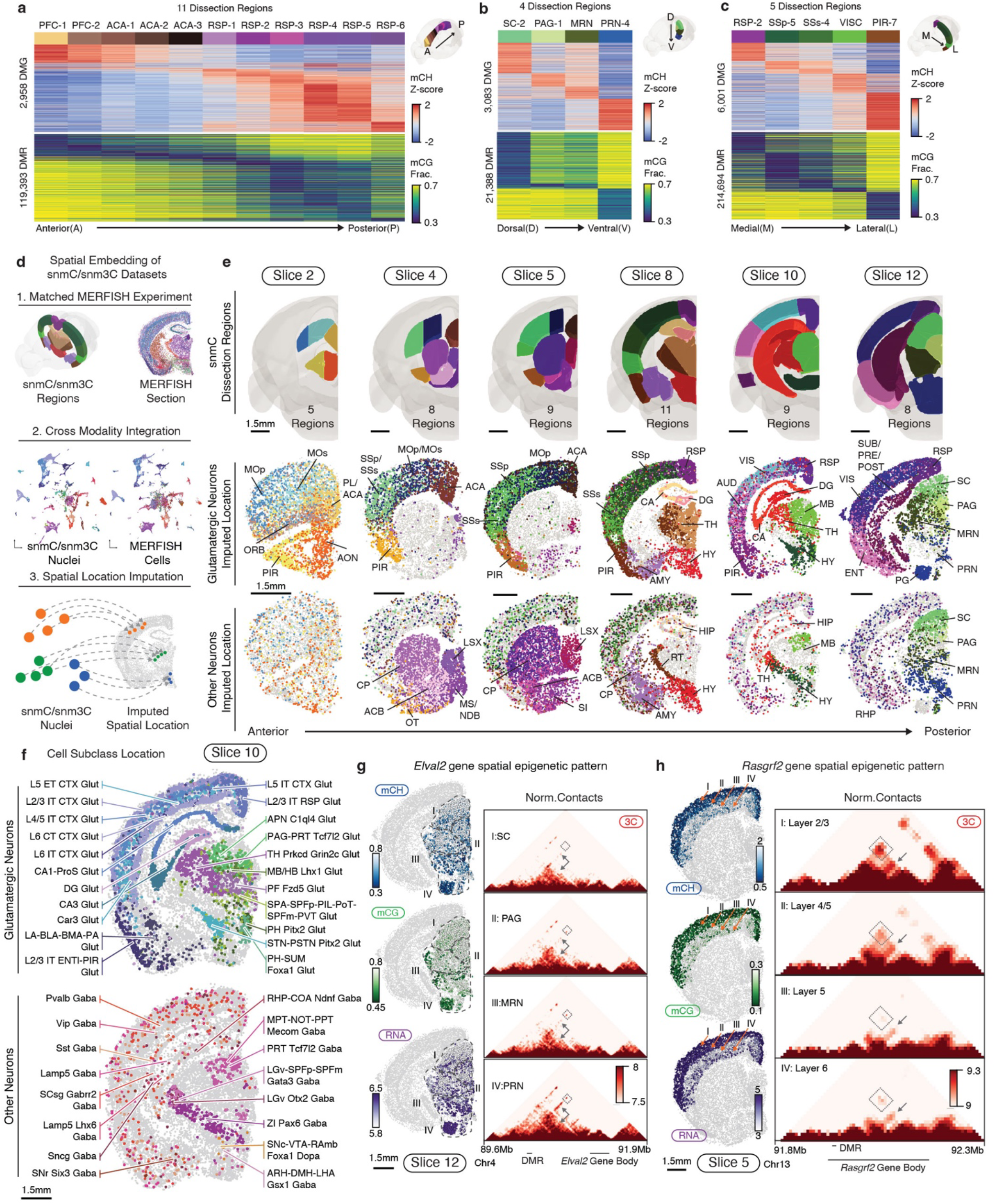
Coherent spatial epigenomic and transcriptomic diversity in the brain. **a-c,** Spatial methylation patterns of DMGs and DMRs across three brain axes (anterior to posterior **(a)**, dorsal to ventral **(b),** medial to lateral **(c)**. **d,** Workflow of mC-MERFISH integration and spatial embedding of methylome cells. **e,** Spatially mapped methylation cell atlas. The first row displays CCF-registered brain dissection regions. The second and third rows show imputed spatial locations for glutamatergic and other neurons colored by dissection regions. **f,** Spatial distribution of cell subclasses for glutamatergic neurons and other neurons on slice 10. **g,** Spatial epigenetic pattern of neuronal genes and their associated DMRs. The *Elval2* gene represents spatial pattern among subcortical regions; the left column shows gene body mCH fraction, DMR (chr13:91,164,342-91,165,792) mCG fraction, and RNA expression. The right column displays the normalized contacts heatmap between the DMR and gene. **h,** The *Rasgrf2* gene and associated DMR (chr13:92,027,775-92,028,983) exhibit cortical layer differences in the same layout as **(g)**.

We organized our dissections into ten major brain regions (Fig. 2b, c, Extended Data Fig. 4) according to their unique cell type composition and neuronal functionality (Fig. 2d): isocortex (CTX), olfactory areas (OLF, including the olfactory bulb and piriform cortex), amygdala areas (AMY, including cortical subplate, CTXsp, and striatum-like amygdalar nuclei, sAMY), cerebral nuclei (CNU, including the striatum and pallidum, but excluding sAMY), hippocampal formation (HPF, including the hippocampus and parahippocampal cortex), thalamus (TH), hypothalamus (HY), midbrain (MB), hindbrain (HB, including pons and medulla), and cerebellum (CB). Most neuronal subclasses (218) are each derived from a single major region. Eighteen neuronal subclasses are situated across two adjacent regions, which could be due to imprecise dissections but may also represent neuronal types shared between neighboring brain regions (Supplementary Table 5). In addition, remarkable cellular diversity is observed in non-telencephalic regions (TH, HY, MB, HB, Fig 2d, e, Extended Data Fig. 5), a common feature observed in other single-cell brain atlases that investigate various molecular modalities^7–9, 12^.

**Figure 4.**
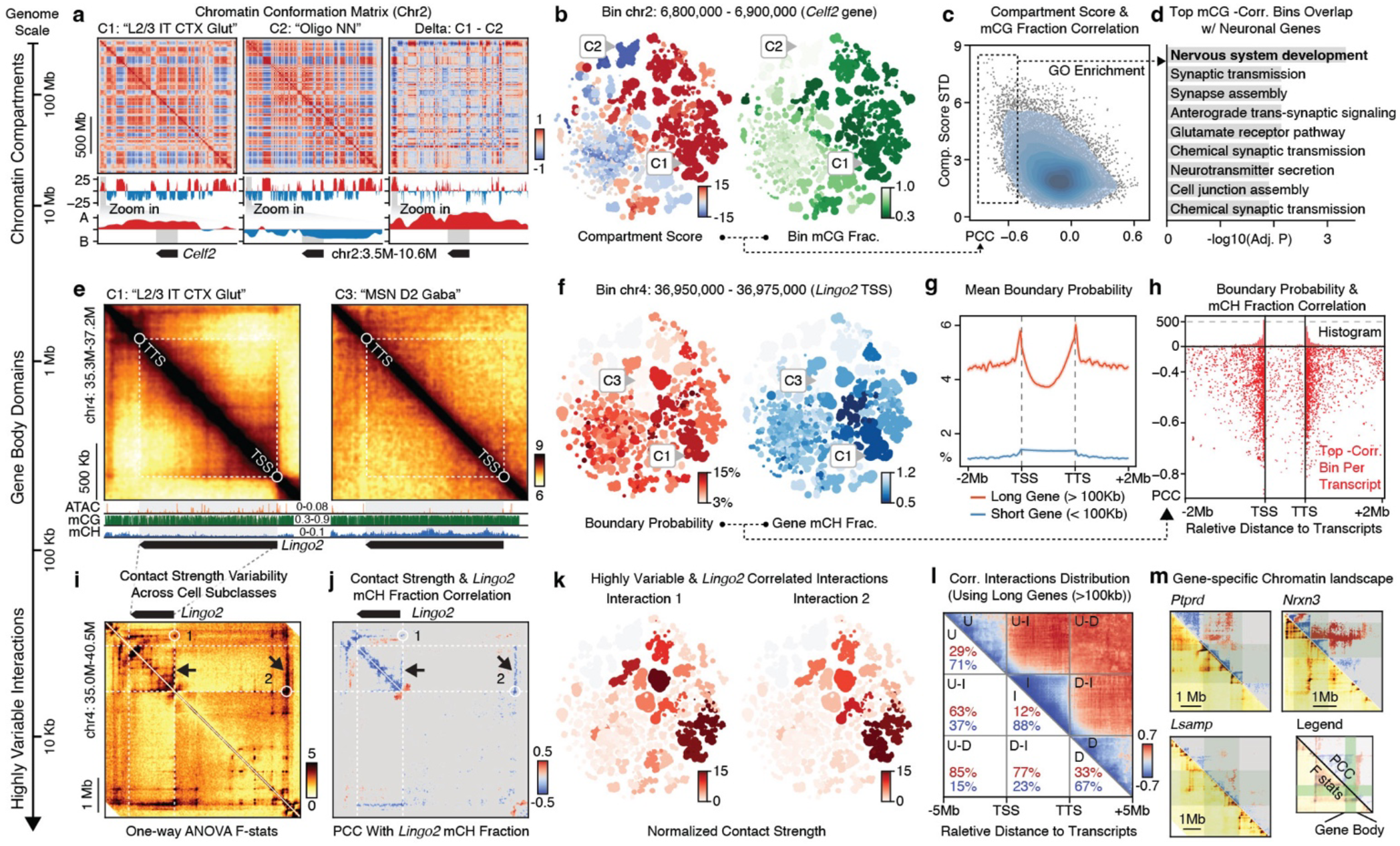
Highly dynamic chromatin conformation features correlate with DNA methylation around neuronal genes. This figure displays chromatin conformation diversity at three levels: chromatin compartments **(a-d)**, gene body domains **(e-h),** and highly variable contacts **(i-m)**. **a,** Top heatmaps are the Pearson-correlation matrices of chr2. Middle plots show the compartment score across chr2 (red and blue indicate A and B compartments, respectively); the bottom row shows the zoom-in view of the *Celf2* gene locus. Three columns from left to right are “L2/3 IT CTX Glut” (C1), “Oligo NN” (C2), and (C1 - C2) delta values. **b,** Cell-group-centroids t-SNE colored by compartment score and mCG fraction. **c,** Scatterplot of chrom100k bins, showing PCC between compartment score and chrom100k mCG fraction (x-axis) and compartment score standard deviation (STD) across cell subclasses (y-axis). The blue contours indicate the dots’ kernel density. **d,** Functional enrichment for genes intersected with negatively correlated chrom100kb bins (boxed in **c**). **e,** Top heatmaps are normalized chromatin contact matrices around the *Lingo2* gene from “L2/3 IT CTX Glut” (C1) and “MSN D2 Gaba” (C3). The bottom genome tracks are the corresponding pseudo-bulk ATAC and methylome profiles. **f,** t-SNE colored by the *Lingo2* TSS boundary probability and *Lingo2* mCH fraction. **g,** Average boundary probabilities of 25kb bins around long and short genes. **h,** The scatterplot shows the location of each long gene transcript’s most negatively correlated boundary. The y-axis is the PCC between the 25Kb bin boundary probabilities and transcript body mCH fractions; the x-axis is the relative genome location to the transcripts. **i,j,** Heatmap of F statistics from one-way ANOVA analysis measuring the variance of contact strength across cell subclasses **(i)** and PCC between the *Lingo2* mCH fraction and highly variable interactions’ contact strengths around the *Lingo2* gene **(j)**. The white circles are two loop-like highly variable interactions. Arrows point to strips between interactions and gene bodies. **k,** t-SNE colored by normalized contact strengths for interactions 1 and 2 in **(j)**. **l,** Pileup view of the relative genome location of correlated interactions from all genes. The colors in the upper triangle are average PCCs. Location categories include intragenic (I), upstream (U), downstream (D), upstream-intragenic (U-I), downstream-intragenic (D-I), and upstream-downstream (U-D). **m,** Heatmap showing chromatin landscape of megabase-long genes, green rectangles indicate the location of gene body, the lower triangle is F statistics similar to **(i)**, and the upper triangle is PCC values similar to **(j)**.

**Figure 5.**
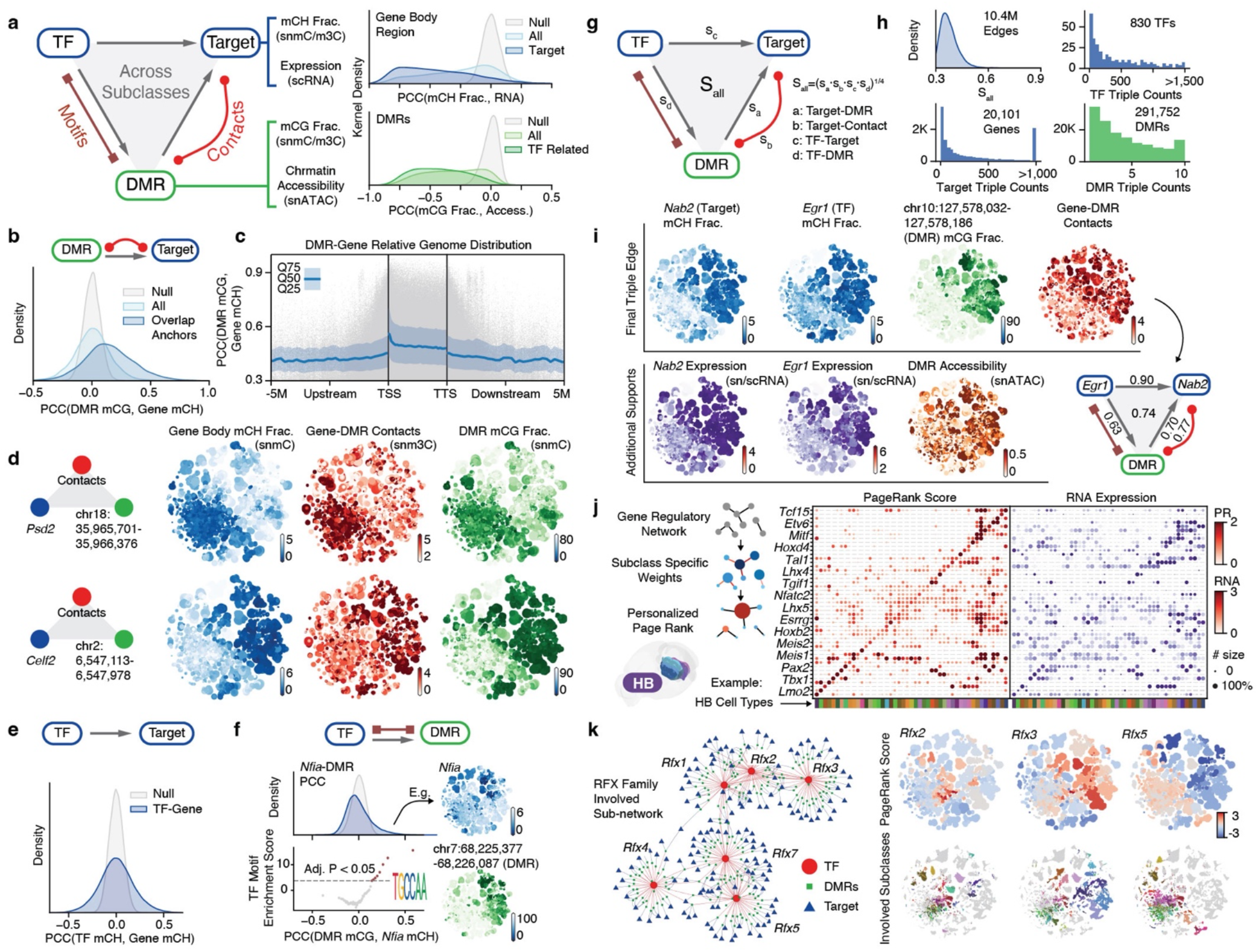
Gene regulatory networks predict binding elements, downstream targets, and cell-type importance of transcription factors. **a,** Schematic depicting the three components of the GRN with two density plots display the PCC between the gene’s mCH fractions and RNA expressions (right top) and the PCC between DMR’s mCG fractions and chromatin accessibilities (right bottom). **b,** the density plot shows the PCC between each DMR’s mCG fractions and the target gene’s mCH fractions. Gray represents the null distribution; shallow blue represents all correlations; blue represents correlations between DMRs overlapping with the target gene’s correlated interaction anchors. **c,** The scatter plot displays the DMR location and PCC between DMR mCG and gene mCH. Each gray dot represents a DMR-target edge. The blue line represents the moving quantile of PCC. **d,** Schematic of the DMR-Target edge for *Psd2* (top row) and *Celf2* (bottom row). From left to right, the t-SNE plot is colored by gene mCH fraction, gene-DMR contacts, and DMR mCG fraction. **e,** the density plot shows PCC between the mCH fraction of TF and the target gene. **f**, Top, PCC between *Nfia* mCH fraction and DMRs mCG fraction. Bottom, cisTarget^65^ motif enrichment score in 50 DMR groups ordered and grouped by the Nfia-DMR PCC value above. The example t-SNE plots are colored by the *Nfia* mCH fraction and mCG fraction of a positively correlated DMR. **g,** Schematic shows the TF-DMR-Target triple and the final score. **h,** Distribution of all triples’ final scores (from **g**) in the final network. Histograms show the number of triples that each TF, gene, and DMR is involved in. **i,** An example triple of *Egr1* (TF), *Nab2* (target), and DMR. t-SNE plot color by the gene’s mCH fraction or RNA level; DMR’s mCG fraction, chromatin accessibility; and gene-DMR contact score. **j,** Left, schematic shows the calculation for PageRange score (methods). Right, dot plots represent TF’s normalized PageRank Score and RNA expression for cell subclasses in the hindbrain (HB). Red dots are colored and sized by PageRank Score. Purple dots are colored by RNA CPM, sized by the percentage of cells in that subclass expressing this gene. **k,** Left, schematic of RFX family sub-networks. Right, t-SNE plot color by normalized PageRank Score (top) and cell subclasses where normalized PageRank score > 0.

Notably, the global methylation level changes dramatically across cells and dissection regions (Extended Data Fig. 3g-h). The global mCG fractions for all cell groups span from 66.3%-85.3% (29.0-37.3 million CpG sites), while mCH fractions range between 0.6%-5.6% (6.9-59.2 million CpH sites). Many subcortical neuronal subclasses exhibit substantially elevated mCH levels compared to excitatory neurons in cortical regions (Extended Data Fig. 3i-j). Examples include “AD Serpinb7 Glut” (from TH, mCH level 5.6%), “PG-TRN-LRN Fat2 Glut” (HB, 5.4%), “CBN Glut” (CB, 5.4%), “SNr-VTA-PPN Pax5 Npas1 Gaba” (MB, 5.2%), “PM-TM-PVp Tbx3 Hist-Gaba” (HY, 4.5%). Since CpH sites (1.1×10^9^) are more abundant than CpG sites (4.3×10^7^) in the mouse genome, the mCH sites in these cells surpass the total number of CpG sites, highlighting the functional significance of this unique neuronal epigenetic modality^15, 28^.

Lastly, our dataset extensively profiles non-neuronal cells and adult immature neurons (IMN) throughout the brain (Extended Data Fig. 4, Supplementary Table 4, 5). Consistent with other modalities^7, 12^, we detected spatial differences in astrocyte methylomes, particularly between telencephalic and non-telencephalic regions. IMNs clustered with astrocytes in the first round, with later iteration resolving one population in the dentate gyrus’ subgranular zone and another found in areas overlapping the rostral migratory stream^29^. Furthermore, the oligodendrocyte lineage demonstrates spatial distinctions between telencephalic and non-telencephalic regions at the cluster level (Extended Data Fig. 4). Our dataset also encompasses other immune and vascular cell types, such as microglia, pericytes, endothelial cells, arachnoid barrier cells, and vascular leptomeningeal cells.

### Consensus cell type taxonomy across modalities

Developing a brain cell type taxonomy requires integrating various molecular modalities, verifying cell clusters based on multiple molecular information, and applying a uniform nomenclature^30^. We began this endeavor by performing an integrative analysis with a brain-wide transcriptome dataset from the BICCN consortium created by Yao et al.^7^. After strict quality control, this single-cell RNA-seq (scRNA-seq) dataset established a cell type taxonomy that categorized 4.3 million cells into 5,200 cell clusters, 1,045 supertypes, 306 subclasses, 32 classes, and 7 divisions. Brain-wide MERFISH datasets^7, 8^ were utilized to incorporate various aspects into the cluster annotation, including spatial distribution, neurotransmitter identity, marker genes, and existing cell-type knowledge^30^.

We employed an efficient framework (adapted from the Seurat package^31^, Methods) for iterative cross-modality integration to leverage this substantial effort. The initial integration effectively matched neuronal spatial distribution and high-level annotations (Fig. 2e), while subsequent iterations refined cluster matching within subclasses to greater detail (Fig. 2f). We utilized integration overlap scores to map methylome cell groups to transcriptome clusters and annotate methylome datasets into subclasses using consistent nomenclatures (Supplementary Table 4). In summary, we matched all methylome cell groups with 4,669 (90%) transcriptomic clusters, encompassing 4.19 million (97.4%) cells corresponding to 261 subclasses (Fig. 2f). The 531 unassigned transcriptomic clusters represent only 2.6% of cells, which are primarily rare populations (<0.03% of total RNA dataset) that are insufficiently represented in the methylome dataset. Based on these integration results, we calculated the transcriptome profile for each cell group (Methods).

The final iteration’s overlap score within each subclass reveals a high-granularity correspondence between methylome and transcriptome clusters (Fig. 2f, boxes). We further examined vital neural functional genes to demonstrate this accurate match between mC and RNA. For instance, neurotransmitter-related genes provide crucial information about cell type identities and display a highly similar specificity between gene body mCH fractions and mRNA expression. Examples include *Slc17a7* and *Slc17a6* for glutamatergic, *Gad1* for GABAergic, *Slc6a5* for glycinergic, *Slc6a2* for noradrenergic, *Th* for dopaminergic, *Chat* for cholinergic, *Slc6a4* for serotonergic, and *Hdc* for histaminergic (Extended Data Fig. 5a-b). In addition, numerous immediate early genes (e.g., *Fos*, *Egr1*, *Arc*, *Bdnf*, *Nr4a2*) are also expressed in many adult brain cell types^7, 9^. Their expression levels are anti-correlated with mCH fractions (Extended Data Fig. 5c). Another gene category includes neuropeptides (e.g., *Npy*, *Vip*, *Sst*, *Penk*, *Pdyn*, *Grp*, *Tac2*, *Cck*, *Crh*), many of which are canonical cell type markers with vital signaling functions^32^. Their specificity is detectable in the gene body mCH that aligns with transcription (Extended Data Fig. 5d). Overall, this high-resolution cross-modality integration offers multi-omic evidence for identifying thousands of cell clusters in the adult mouse brain, laying the groundwork for subsequent genomic and epigenomic analyses.

## Multi-omic evidence for cell-type-specific cis-regulatory elements

Having established a consensus cell taxonomy across the entire mouse brain, we further identified 2.56 million non-overlapping CpG DMRs between the subclasses of the whole brain or the clusters of each major brain region (Methods). These DMRs involve 44% of the total CpG sites in the genome, with an average length of 189±356 (mean±s.d.) and containing 3.9±6.0 CpG pairs (each containing two bases). The CpG DMRs provide predictions about cell-type-specific cis-regulatory elements, and hypo-methylation in the DMR region usually indicates the active regulatory status in adult brain tissue^10, 22^ (Fig. 2g). To annotate the accessibility status of the DMRs systematically, we performed iterative integration between the methylome and chromatin accessibility dataset from Zu et al.^12^, using non-overlapping chromosome 5kb bins (Methods). This dataset, generated by snATAC-seq without NeuN enrichment by FANS, contains 1,372,646 neurons and 939,760 non-neuronal cells. As this dataset shares the same dissection samples with the snmC-seq dataset, we used this ground truth information to assess the integration alignment score^31^ between mC and ATAC neurons. Remarkably, the dissected regions are precisely aligned (score 0.89±0.11), indicating extensive concordance in the cellular diversity of both epigenomic modalities (Extended Data. Fig. 6a, b). After integration, we also calculated the chromatin accessibility profile for each cell group using their matched ATAC clusters. The resulting mCG fractions and chromatin accessibility levels at DMR regions show similar cell-type-specificity across brain cell subclasses, confirming the correct match of cell-type identities. (Fig. 2g, Extended Data Fig. 6c, d). By integrating the methylome and chromatin accessibility datasets, we achieved remarkable concordance in cellular diversity across both epigenomic modalities, further validating the accuracy of our approach in determining cell-type-specific regulatory elements and their activities.

**Figure 6.**
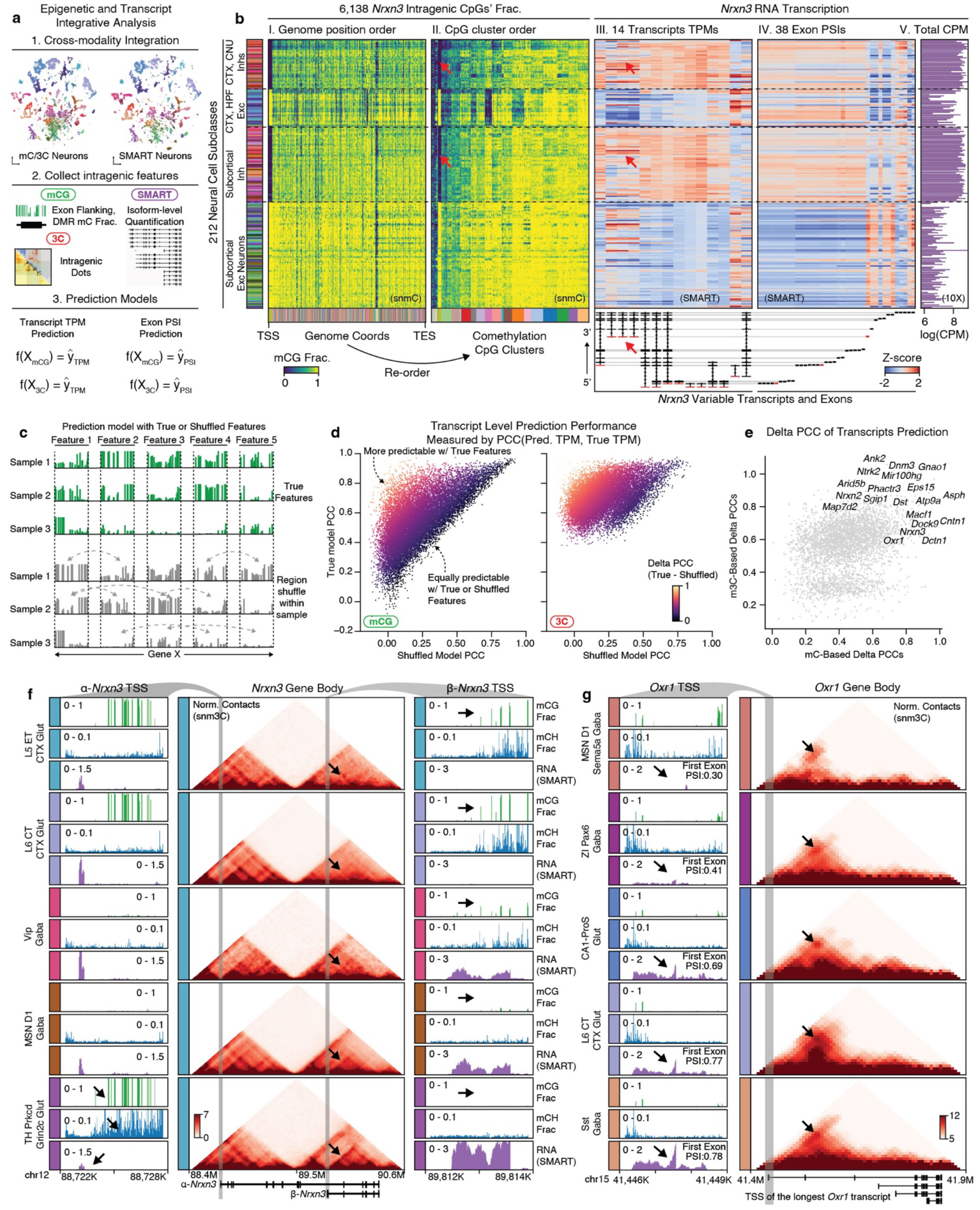
Epigenetic heterogeneity predicts gene isoform diversity. **a,** Workflow for the integrative analysis between epigenome and transcriptome datasets. **b,** Compound heatmaps illustrate the similarity between the *Nrxn* intragenic methylation heterogeneity and alternative isoform expression patterns. Rows are neuron cell subclasses. I, mCG fraction of all 6,138 CpG sites of *Nrxn* gene with columns ordered by original genome coordinates (bottom colors are CpG clusters from heatmap ll). ll, mCG fraction of CpG sites re-ordered by their CpG clusters (bottom colors) based on subclasses methylation pattern. Heatmap lll and Heatmap lV show the TPM of 14 highly variable transcripts and PSI of 38 highly variable exons of *Nrxn*, quantified with the SMART-seq dataset. All values are z-score normalized across cell subclasses. The *Nrxn* transcript structures and exon locations are indicated at the bottom plots. Red arrows point to beta-Nrxn3 transcripts and one associated CpG cluster. Heatmap V shows the *Nrxn* gene log(CPM) in scRNA-seq (10X) data. **c,** Schematic illustrates the process for constructing the prediction model. **d,** Scatterplot shows the PCC between predicted TPM and true TPM for each highly-variable transcript (dot), using methylation features (left) and chromatin contact interactions (right) to predict. **e,** Scatterplot shows the delta PCC in mC models (x-axis) and m3C models (y-axis) for highly-variable transcripts (dot). Top transcripts with large delta PCC are listed by their corresponding gene names. **f.** Genome browser view of intragenic epigenetic and isoform diversity of the *Nrxn* gene in five cell subclasses (rows). The middle heatmaps are normalized contact strengths of the *Nrxn* gene locus, with arrows pointing to strips over the beta-*Nrxn* transcript body. The zoom-in panels show alpha-*Nrxn3*’s (left) and beta-*Nrxn3*’s (right) TSS region, with mCG fraction (green), mCH fraction (blue), and SMART RNA (bottom) expression tracks. **g,** Similar to **f,** showing the corresponding intragenic epigenetic and isoform diversity in the *Oxr1* gene.

### Coherent spatial epigenomic and transcriptomic diversity in the brain

Tens of millions of cells in the mouse brain accurately form complex anatomical structures controlled by their diverse gene expression and epigenetic regulation. Our clustering analysis has demonstrated cell type composition differences across brain regions (Fig. 2). To explore the spatial information further in the DNA methylome, we performed differentially methylated gene (DMG) and DMR analyses across anterior-to-posterior, dorsal-to-ventral, and medial-to-lateral axes in the brain using representative dissection regions (Fig. 3a-c). In all three axes, we identified hundreds or thousands of DMGs related to various neuronal functions and DMRs associated with these genes, highlighting the remarkable spatial diversity encoded in the methylome.

To increase spatial resolution and investigate whether the observed methylation spatial pattern corresponds to actual transcriptomic diversity, we employed the MERFISH technology, which enables *in situ* profiling of hundreds of genes’ expression in brain sections^2, 8, 33^. We designed a 500-gene panel (Supplementary Table 6) selected based on cell type and spatial diversity in gene body hypomethylation across the brain (Methods) and profiled six coronal sections corresponding to our mC and m3C brain slices (Extended Data Fig. 7a). After quality control, we obtained 266,903 MERFISH cells and annotated their cell types by integrating with the scRNA-seq dataset (Extended Data Fig. 7b-d, Supplementary Table 7)^7^. We then performed cross-modality integration between the neurons in the methylome and MERFISH datasets, imputing the spatial location of each methylation nucleus (Fig. 3d, Supplementary Table 8). Interestingly, the predicted spatial coordinates of the methylation nuclei closely matched the dissected regions (Fig. 3e). For example, glutamatergic cells show arealization among cortical areas within each slice; dorsal-ventral separation is observed among medium spiny neurons dissected from the Caudoputamen (CP) and Nucleus Accumbens (ACB) regions; many subcortical dissection borders are faithfully preserved in the imputed spatial embedding. Furthermore, the spatial location imputation also assigned many cell types to fine anatomical structures, which were considerably smaller than our dissection regions (Fig. 3f, Extended Data Fig. 8). For instance, laminar layer information was mapped among cortical excitatory cells. In addition, many subcortical neurons were allocated to specific brain nuclei (e.g., “STN-PSTN Pitx2 Glut”, “LGv Otx2 Gaba”, “ZI Pax6 Gaba”), highlighting the correspondence between the cell-type identity and anatomical structure in the subcortical areas.

The high spatial resolution in the imputation was attributed to the strong association between cell location and DNA methylation of critical genes and regulatory elements. For example, the *Elavl2* gene, an RNA-binding protein involved in post-transcriptional regulation functions in neurons^34^, exhibited a dorsal-ventral increased expression pattern in subcortical neurons in Slice 10, which was also observed as the decrease of gene body mCH methylation of *Elavl2* and a nearby DMR’s mCG methylation (Fig. 3g). Notably, the chromatin interactions between the DMR and *Elavl2* gene showed stronger contacts in regions where *Elavl2* was highly expressed. Likewise, *Rasgrf2*, a guanosine nucleotide exchange factor for Ras GTPases, displayed differential expression and methylation across cortical layers. DMRs near *Rasgrf2* were highly correlated, with chromatin conformation data supporting physical proximity when both the DMR and *Rasgrf2* were active (Fig. 3h). *Negr1* also shows similar correspondence among modalities in cortical dissected regions (Extended Data Fig. 7e). These findings demonstrate a clear spatial pattern in DNA methylation that aligns with the spatial transcriptome, implying that epigenetic regulation exerts precise control over the cellular spatial location across the entire brain.

### Chromosomal conformation dynamics across brain cells

The annotated multi-omic datasets enabled us to leverage the cell-type diversity across the entire brain to understand the chromatin conformation landscape of individual genes at multiple genomic scales. Here, we systematically evaluated the variability of different 3D genome features (chromatin compartment, topologically associated domain (TAD), and highly variable interactions) across cell subclasses and associated them with gene activity by correlating chromatin contact strengths with methylation fractions.

We initiated this effort by examining the chromatin compartment, a genome topology feature bringing together the genomic regions tens to hundreds of megabases away^35^. The genomes are organized in two major compartments, A and B, corresponding to the active and silent chromatin^35, 36^. After calculating the compartment score of cell subclasses at the 100-Kb resolution, we observed numerous A/B compartment switches in megabase-long regions (Fig. 4a). For instance, the chromosome 2 region spanning 3.5M to 10.6M exhibited a strong negative compartment score (B compartment) in mature oligodendrocytes (“Oligo NN”), but positive scores (A compartment) in cortical excitatory neurons, such as “L2/3 IT CTX Glut” (Fig. 4b). Notably, this compartment-switching region overlaps with the *Celf2* gene, a vital RNA-binding protein that modulates alternative splicing in neurons^37^.

Given these observations, we sought to determine if compartment switching correlated with DNA methylation changes within the same regions. Upon calculating the PCC across cell subclasses, we found a negative correlation between the compartment score and mCG or mCH fraction of 100Kb chromatin bins, with mCG exhibiting a stronger correlation than mCH (Extended Data Fig. 9a). Additionally, we observed that the compartment score of negatively correlated bins demonstrated greater variability across cell subclasses (Fig. 4c, Extended Data Fig. 9b, c), suggesting that these negatively correlated bins exhibit dramatic activity change across a wide range of cell types.

We then discovered that genes overlapping with the negatively correlated bins were enriched^38^ in numerous critical neuronal-related functions, including nervous system development (Fig. 4d). To explore this further, we examined another mouse developing brain scRNA-seq atlas^39^ and found that the negatively correlated bins overlapped with genes displaying a dramatic increase in expression during prenatal brain development. In contrast, uncorrelated or positively correlated bins demonstrated no such trend (Extended Data Fig. 9d). These results suggest that dramatic chromosomal conformation changes might be established during early development and subsequently maintain cellular specificity in adult brain nuclei.

### TAD boundaries associated with long gene body regions

We also investigated the TADs^40^ and their boundaries at a 25-Kb resolution (Methods). By first identifying boundaries in individual cells and subsequently using the domain boundary probability at the cell subclass level, we were able to represent the strength of domain boundaries at each 25Kb bin (Extended Data Fig. 9e). To evaluate the variability of boundary probabilities across the genome, we performed a Chi-square test on each bin and identified 83,518 bins with significant variability across subclasses (false discovery rate, FDR < 1e-3, Methods). For example, we observed that at the *Lingo2* locus, an “L2/3 IT CTX Glut” hypo-methylated gene linked to essential tremor and Parkinson’s disease^41^, the TAD boundaries align with the gene’s TSS and transcription termination site (TTS) (Fig. 4e). Across all the neuronal subclasses, the boundary probability of the 25Kb bin at the *Lingo2* TSS exhibits a negative correlation with the transcript body mCH fraction (Fig. 4f, PCC=-0.65, FDR < 0.001, permutation-based test, Methods).

To generalize this observation, we calculated the average boundary probability at all gene TSSs and TTSs in the genome, separating them by transcript length (< 100Kb as short, > 100Kb as long^17, 42^). Long genes displayed elevated levels of boundary probability at the TSSs and TTSs (Fig. 4g), suggesting that TADs are more likely to form around the gene body (i.e., gene body domains). Our analysis then focused on the relationship between variable domain boundaries and gene bodies, particularly long genes (> 100Kb) implicated in neuronal pathogenicity and potentially regulated by mCH and MeCP2^17^. We next calculated the PCC of gene transcript body mCH or mCG fractions with the boundary probabilities of all 25Kb bins within transcript ± 2Mb distances (Extended Data Fig. 9f). The top negatively correlated boundaries were predominantly located at the TSSs and TTSs of the corresponding gene transcripts (Fig. 4h, Extended Data Fig. 9f, g). Additionally, we observed a few significantly positively correlated boundaries to the transcript body mCH or mCG, although they lacked clear TSS/TTS colocalization (Extended Data Fig. 9f, g). Functional enrichment analysis^38^ revealed that genes with strongly negatively correlated gene body domains were significantly enriched for critical neuronal and synaptic functions. In contrast, positively correlated TAD boundaries were not associated with genes enriched for specific functions (Extended Data Fig. 9h). Together, these results indicate that TAD boundaries are closely associated with the transcription start and termination sites of long genes implicated in neuronal pathogenicity and critical functions.

### Diverse neuronal gene chromatin conformation landscapes

In order to thoroughly profile the chromatin conformation diversity at high resolution and link genes to their potential regulatory elements, we analyzed chromatin interactions at the 10-Kb resolution (Extended Data Fig. 10a). We first performed a one-way analysis of variance (ANOVA) across cell subclasses and used the F statistics to summarize the variability of all interactions. Highly variable interactions correspond to dot or strip-like patterns around genes (Fig. 4i).

Subsequently, we calculated the PCC between transcript body mCH fraction and the contact strength of highly variable interactions within ± 5 Mb of the transcript body (Fig. 4j). Highly variable and gene-correlated interactions were assigned to a gene if any anchors of the interaction overlapped with the gene body. Through this assignment, the majority (95%) of gene-associated interactions are located within 1.2 Mb of the gene’s TSS (Extended Data Fig. 10b). Genes with numerous correlated interactions exhibit crucial neuronal and synaptic functions, overlapping with those genes that displayed a negatively correlated gene body domain boundary in the previous section (Extended Data Fig. 10c, d). For instance, in the *Lingo2* locus, highly variable interactions were identified within the gene body, at gene body domain boundaries, or corresponding to distal loop structures^36^ (Fig. 4j, circles). The correlation analysis further stratified interactions positively or negatively correlated with the gene’s methylation change. Notably, the correlated interaction anchors can be up to 1.6 Mb downstream (interaction 1) or 3.2 Mb upstream of the *Lingo2* TSS (interaction 2) while associating with strips along the entire gene body (Fig. 4j, k).

We then summarized the distribution of significantly correlated interactions surrounding all long genes by categorizing them into six groups based on their relative location to the gene: intragenic (I), upstream (U), downstream (D), upstream-intragenic (U-I), downstream-intragenic (D-I), and upstream-downstream (U-D) (Fig. 4l). Our results revealed that the contact strength of intragenic, upstream, and downstream interactions are mostly negatively correlated with gene body methylation (% negative PCC, I: 88%, U: 71%, D: 67%), consistent with the observation that the gene body domain forms between the TSS and TTS, insulating the interactions between I, U, and D while increasing their interaction within each group. Moreover, the U-I and D-I interactions are primarily positively correlated with gene body methylation (% positive PCC, U-I: 63%, D-I: 77%). However, the negatively correlated interactions likely remain critical as they potentially link distal regulatory elements to intragenic regions (Fig. 4j). U-D interactions exhibit the least negative correlations (% negative PCC, U-D: 15%) and do not directly interact with the gene body, potentially relating to higher-level chromatin conformation regulation.

Despite these general trends, the specific chromatin conformation landscapes of individual genes are remarkably diverse (Fig. 4m). In addition to the intriguing U-I and D-I interactions observed in the *Lingo2* gene, many megabase-long genes display complex intragenic subdomain patterns (e.g., *Ptprd*, *Nrxn3*, *Lsamp*, *lg2*, *Celf2*, *Sox5* in Fig. 4m and Extended Data 10e-j). These patterns may correspond to more subtle gene activity regulation, including alternative TSS and exon usage, which will be explored later.

### The multi-omic analysis unveils gene regulatory networks

Numerous critical transcription factors orchestrate the intricate spatial and cell-type-specific gene expression patterns within Gene Regulatory Networks (GRNs), which can be elucidated using multi-omic information^43, 44^. Here, we present a framework that connects transcription factors (TFs) with DMRs and their potential downstream target genes, leveraging DNA methylome and chromatin conformation signals to construct GRNs for whole-brain neurons (Fig. 5a, left part, Methods). Our approach employs mCH fractions as proxies for gene status and mCG fractions as indicators of regulatory element activity. To further support our findings, we incorporate integrated transcriptome and accessibility profiles as complementary evidence due to their strong negative correlation with gene mCH and DMR mCG fractions, respectively (Fig. 5a, right part).

We built connections between (1) DMRs and their potential target genes (DMR-Target edge); (2) TFs and their potential target genes (TF-Target edge); (3) TFs and their potential binding DMRs (TF-DMR edge). We established DMR-Target edges by accounting for the correlation of methylation fractions between the DMR and surrounding genes, as well as the gene’s chromatin conformation landscape discussed earlier (Fig. 5b, Methods). This approach intersected the diversity of both modalities measured in our snm3C-seq assay by limiting correlation-based edges to genome regions displaying pronounced chromatin conformation changes. This step generated 1.2×10^6^ edges between 5.7×10^5^ DMRs and 2.1×10^4^ genes (Fig. 5c), with 27% of edges connecting intragenic DMRs to genes and 73% linking distal DMRs. For instance, the edges of the *Psd2* and *Celf2* genes demonstrate highly concordant cell-type-specificity of DNA methylation and chromatin interaction between gene bodies and their associated DMRs (Fig. 5d). We proceeded to connect TF-Target edges based on their correlated methylation fractions (Fig. 5e). We identified a total of 4.6×10^6^ edges between 1,705 TFs and 2.6×10^4^ genes. Since the TF-Target edge alone is insufficient to discern gene regulation relationships^44, 45^, we also quantified the TF-DMR edges, which indicate potential regulatory elements that the TF used to control target gene expression.

We established the TF-DMR edge by considering the correlation of methylation fractions between the DMR and TF gene body and the enrichment of TF binding motifs in the correlated DMR sets (Extended Data 11a, Methods). In the motif enrichment analysis, we discovered that many TFs have their motifs solely enriched in the DMRs that positively correlated with TF gene body methylation, such as the *Nfia*, *Onecut2*, and *Rfx1* (Fig. 5f, Extended Data Fig. 11b). This finding implies that the binding of these TFs potentially activates the underlying regulatory elements. Intriguingly, we also observed some TFs with motifs enriched in negatively correlated DMRs, such as the *Foxp2* and *Foxa1* genes (Extended Data Fig. 11c). Both TF genes were reported to have transcription repression functions^46, 47^, potentially achieved by repressing active enhancers. We identified 1.2×10^7^ edges between 843 TFs and 4.6×10^5^ DMRs (Extended Data Fig. 11d).

We combined all three types of edges (DMR-Target, TF-Target, TF-DMR) to construct the final GRN with TF-DMR-Target triples. Each triple is assigned a final score representing the overall correlation of cell-type specificity between the three components (Fig. 5g, Methods). The resulting network comprises 1.04×10^7^ triples, involving 830 TFs, 20,101 genes, and 291,752 non-overlapping DMRs (Fig. 5h). The different combination of correlations in a triple provides insights into regulatory relationships between the TF, DMR, and target gene (Extended Data Fig. 11e). We summarized eight possible combinations into four models and one unknown category (Extended Data Fig. 11f). The most frequent model (39.8%) represents all-positive correlations, indicating that both the TF and DMR have an active effect on the gene (Model 1). The second most frequent model (30.5%) is negative for TF-Target and TF-DMR edges and positive for DMR-Target edges, suggesting that the TF plays a repressive role by repressing active DMRs. These two models account for most (70.3%) of the edges, indicating that intersected DMRs predominantly activate target genes. The third (Model 3, 11.1%) and fourth (Model 4, 11.1%) most popular models are negative or repressive for DMR-Target edges, with Model 3 suggesting that active TFs turn off repressive DMRs to activate genes and Model 4 indicating that repressive TFs turn on repressive DMRs to deactivate genes. The remaining edges are assigned to the “Unknown” group (Extended Data Fig. 11e), likely intersected by chance or representing indirect relationships involving additional regulatory factors. Models 1 to 4 cover 92.5% of the edges, demonstrating a remarkable correspondence between these three genomic elements among the brain-wide cell types.

In addition, the individual TF-DMR-Gene triples predict numerous TF and gene relationships, pinpointing their intermediate regulatory elements. These relationships are supported by the DNA methylome and chromatin conformation, as well as the integrated transcriptome and chromatin accessibility. For example, one high-scoring triple (0.74) links the critical neuronal TF *Egr1* to its downstream target gene *Nab2* through a distal DMR (chr10:127,578,032-127,578,186, Fig. 5i). The *Nab2* gene expression is known to be induced by *Egr1*, and the NAB2 protein then represses *Egr1* activation function, forming a negative feedback loop^48^. Moreover, the *Egr1* cofactor *Erf*^49^ is also connected to *Nab2* through another DMR (Extended Data Fig. 12a, b). In addition to these known examples, another interesting edge connects *Egr1* with the *Synpo* gene, which encodes an actin-associated postsynaptic protein, with a DMR located in its upstream correlated regions (Extended Data Fig. 12c, d). A second example is the link between the subcortical expressing TF *Stat5b* and the *Cacna2d2* gene, which encodes a calcium voltage-gated channel auxiliary subunit, connected by an intragenic DMR located in the highly correlated gene body domain (Extended Data Fig. 12e, f). These intriguing examples demonstrate the power of our approach in identifying novel and biologically relevant gene regulatory relationships by leveraging multi-omic data.

### Weighted GRNs identify key transcription factors

Transcription factors play a crucial role in regulating cell identity^50^. In order to demonstrate the importance and specificity of transcription factors within each cell subclass, we utilized the comprehensive GRN combined with the Taiji framework^51, 52^. Using the PageRank algorithm, this framework identifies key transcription factors by propagating gene and regulatory element information on the GRN with node and edge weights specific to each cell subclass.

Focusing on the hindbrain (Fig. 5j) and midbrain (Extended Data Fig. 12g) as examples, we discovered key transcription factors that exhibit highly specific PageRank scores among cell subclasses within these complex brain regions. The combination of transcription factor PageRank scores uniquely identifies each cell subclass in these regions, aligning with their respective transcription specificities. Notably, the PageRank score can capture the specificity of even extremely lowly expressed transcription factors (Fig. 5j), likely due to gene body methylation measurements. Additionally, we observed numerous transcription factors within the same family exhibit distinct cell-type-specificity (Fig. 5k). For example, the *Rfx* gene family^53^ has six members variably expressed in adult mouse brains. Their connectivity on the GRN and subclass-specific PageRank scores reveal that these members could play distinct regulatory roles that partially overlap. For example, *Rfx2* is predicted to be critical in HY and MB cell types. *Rfx3* has high PageRank scores in cell types overlapping with *Rfx2* in subcortical areas but is also inferred as an important regulator broadly in cortical excitatory neurons. *Rfx5* is predicted to be important in a wider range of cell types, including the majority of subcortical neurons, cortical inhibitory neurons, comprehensive GRN and the PageRank algorithm astrocytes, and oligodendrocyte progenitors. The effectively identify key transcription factors with high cell-type specificity in diverse brain regions. This approach generates numerous predictions about transcription factor functions in determining cell identity, paving the way for future perturbation experiments^54^.

### Intragenic epigenetic heterogeneity predicts isoform diversity

Alternative splicing leads to the production of different isoforms from the same gene, and its dysfunction in the brain has been associated with various neurodevelopmental disorders^55^. It is regulated by various RNA-binding proteins and has recently been associated with DNA methylation^56, 57^. The diversity of isoform expression has been reported in several cortical cell types^27, 58^. However, their diversity in a considerably wider range of cell types across the whole mouse brain and their relationship with the epigenome remains to be elucidated. To investigate these questions, we integrated the snmC and snm3C-seq datasets with a companion full-length single-cell RNA-seq (SMART-seq v4) dataset from AIBS, which contains 195,680 cells covering the entire adult mouse brain^7^ (Methods). This integration allowed us to explore the intragenic diversity of DNA modification and topology in conjunction with RNA transcript/exon level measurement at cell-group resolution (Fig. 6a, Methods).

To exemplify this framework, we first examined the methylation pattern of the neurexin 3 (*Nrxn3*) gene, a critical presynaptic gene known to express thousands of alternative isoforms^59^. Within the *Nrxn3* gene body, we observed multiple comethylated CpG clusters grouped by their methylation patterns across cell subclasses (Fig. 6b, box I, II). Note that these CpG clusters are different from traditionally described CpG islands, by grouping together the distal CpGs showing similar methylation specificities across cell types. Intriguingly, many of these CpG clusters showed similar cell-subclass-specificity to the alternative usage of certain *Nrxn3* transcripts and exons (Fig. 6b, box III-V). For instance, the functional isoform of the truncated beta-neurexin^59^ was predominantly active in inhibitory and a few excitatory cell subclasses, corresponding with a CpG cluster hypomethylated in the same populations (Fig. 6b, arrows). Similarly, the neuron-specific antioxidant gene *Oxr1* exhibited intragenic methylation heterogeneity that matched the diversity of several transcripts and exons (Extended Data Fig. 13a).

To systematically analyze this phenomenon, we conducted a machine-learning-based analysis to quantify the predictability of alternative splicing using intragenic DNA methylome or chromatin conformation features in each cell group (Methods). Specifically, asking how much improvement we can obtain by incorporating high-resolution intragenic features to predict isoform expression levels, compared to using whole gene body measurements as a proximate averaged activity of isoforms.

To assess this, we trained two models for each gene (Fig. 6c): one with the true intragenic features, and another using within-sample shuffled features that disrupted intragenic correspondence but still preserved the sample-level information for each gene. We calculated PCC between the predicted and true values across cell groups for both models. The delta PCC value between the true and shuffled models represented the gain in predictability through adding intragenic features (Fig. 6d, Extended Data Fig. 13b). Many crucial neuronal and synaptic genes known for functional alternative isoform expressions exhibited a large delta PCC in their highly-variable transcripts and exons (e.g., *Nrxn1-3*^59^, *Ntrk2*^60^, *nm3*^61^, *Oxr1*^62^, Fig. 6e, Extended Data Fig. 13c). Interestingly, chromatin conformation features demonstrated better overall prediction accuracy than DNA methylation in these alternatively spliced genes (Fig. 6d, Extended Data Fig. 13b), possibly because these features account for genome 3D interaction, while methylation features only consider 1D. This observation aligns with the understanding that many alternative splicing events involve nuclear compartmentalization and long-range genome interactions^63^.

Finally, the prediction models prioritize specific transcripts and exons whose alternative usage is more likely under epigenetic regulation. We evaluated several representative examples in the genome browser, such as the alpha and beta-*Nrxn3* promoters. The canonical alpha promoter has a low expression in the “TH Prkcd Grin2c Glut” subclass, evidenced by high mCG and mCH fractions downstream of the promoter. In contrast, the beta-promoter shows the highest expression level in the “TH Prkcd Grin2c Glut” subclass, with depleted surrounding methylation. Intriguingly, the transcript body domain of beta-*Nrxn3* exhibits associated interaction changes among cell types with different beta-*Nrxn3* expression (Fig. 6f). Similarly, the first exon (ENSMUSE00000683442) of the longest *Oxr1* transcript displays increased usage (Percent Spliced In, PSI) among representative cell subclasses, accompanied by corresponding methylation and chromatin conformation changes in the surrounding regions (Fig. 6g). Together, these results highlight the complex interplay between epigenetic regulation and alternative splicing, unveiling potential cell-type-specific regulatory mechanisms contributing to the brain’s post-transcriptional diversity of neuronal and synaptic genes.

## Discussion

This study presents a single-cell DNA methylation and 3D multi-omic atlas of the entire mouse brain. By employing methylome-based clustering and cross-modality integration with additional BICCN companion datasets^7, 12^, we established a cell type taxonomy consisting of 4,673 cell groups and 261 subclasses. Our integrative approach combines five molecular modalities—gene mCH, DMR mCG, chromatin conformation, accessibility, and gene expression—to create a multi-omic genome atlas featuring thousands of cell-type-specific profiles. Furthermore, we identified 2.6 million DMRs at two clustering granularities, offering a vast pool of candidate regulatory elements for various analyses. Impressively, the intricate cellular diversity within the mouse brain exhibits extensive concordance across all molecular modalities, as evidenced by the aligned cell-type-specific patterns observed in numerous essential neuronal genes (Extended Data Fig. 5) and groups of regulatory elements (Extended Data Fig. 6). These findings underscore the fundamental interplay between epigenetics and transcriptomics in shaping the brain’s cellular diversity and serve as a foundation for incorporating additional complementary molecular modalities (such as histone modification, 5hmC, translatome, and proteome) in future efforts to construct a holistic molecular representation of the brain.

Notably, we also observed extensive spatial diversity encoded within the DNA methylome across the entire mouse brain. This epigenetic spatial pattern demonstrates a high concordance with spatial transcriptional diversity, as evidenced through integration with a MERFISH dataset generated from spatially diverse methylated genes. By leveraging these datasets, we achieved a detailed spatial map of DNA methylation and chromatin conformation profiles within delicate brain structures. The results offer a valuable anatomical context for methylation and 3D multi-omic cell data and emphasize the considerable influence of epigenetic regulation on spatial cell organization within the brain.

Building on the foundation of our high-resolution, spatially annotated multi-omic brain cell atlas, we expanded our investigation to the mouse genome to explore the underlying gene regulatory diversity across multiple scales. At the whole chromosome level, the chromatin compartment identity of megabase-long regions can undergo significant alterations among different brain cell types. These changes negatively correlate with DNA methylation, particularly at mCG sites. Genes within these regions play critical roles in neuronal functions, especially in neurodevelopment. Additionally, we observed that TAD boundaries tend to form around neuronal long genes, with a negative correlation identified between boundary probability and the transcript body mCH fraction. A recent discovery of a similar gene boundary feature termed the transcription elongation loop offers a potential explanation for the higher gene domain boundary probability observed^64^. However, the mechanism by which the diversity of this feature arises across various cell types within the brain remains to be elucidated. Moreover, we conducted an unbiased investigation of the chromatin conformation context surrounding individual genes by performing ANOVA and correlation analyses using whole-brain populations. This approach yields unprecedented gene chromatin conformation landscapes that reveal general rules governing the relationship between chromatin interaction and gene body methylation and offer gene-specific predictions on the importance of individual chromatin interaction pixels at a 10-kb resolution.

Integrating the extensive gene, DMR, and chromatin conformation data enables us to construct a comprehensive GRN for gene regulation in the mouse brain. This network predicts regulatory relationships between TFs and their target genes through the precise DMRs containing TF binding motifs. Furthermore, numerous TF motifs are strongly enriched in DMRs where mCG fractions correlate positively or negatively with the TF mCH fraction, suggesting dominant activation or repression roles for the corresponding TFs. Personalized PageRank analysis of the GRN identifies the most influential TFs for each cell subclass in subcortical regions characterized by vast cellular diversity. The GRN also reveals diverse cell-type-specific patterns among members of the same TF family. Finally, the high-resolution methylome and chromatin conformation data enable us to examine the relationship between epigenetic modalities and alternative isoforms. Our findings suggest that extensive intragenic epigenetic heterogeneity may contribute to regulating alternative promoter and exon splicing in these genes. The predictive model identifies top candidates for further investigation into their causal relationships.

In summary, our analysis underscores the potential of this whole-brain dataset to characterize cellular, spatial, and epigenomic diversity at unprecedented resolution. Furthermore, this resource offers valuable insights into the fundamental gene regulation principles that shape the remarkable complexity of the mammalian brain, laying the groundwork for a deeper understanding of the molecular underpinnings of the human brain.

## Acknowledgments

We thank members of the BICCN community for discussions; This work utilized the Stampede2 supercomputing resources at Texas Advanced Computing Center through allocation MCB130189 from the Extreme Science and Engineering Discovery Environment. We also thank the SkyPilot developers from UC Berkeley Sky computing lab for helping us establish cloud computing frameworks. This work was supported by grants from NIMH, U19MH114831, and U19MH114831-04S grant to J.R.E., B.R. and Edward Callaway, and U19MH114830 to H.Z., all under the BRAIN Initiative of National Institutes of Health (NIH). The Flow Cytometry Core Facility of the Salk Institute is supported by grant NIH-NCI CCSG: P30 014195 and Shared Instrumentation Grant S10-OD023689. H.L. is supported by the Pioneer Fund Postdoctoral Scholar Award. J.R.E is an investigator of the Howard Hughes Medical Institute.

## Author Contributions

J.R.E., H.L., B.R., M.M.B. conceived the study. H.L., Q.Z., J.Z. analyzed the data and drafted the manuscript. J.R.E., C.L., B.R., J.R.D., M.M.B, H.Z., B. T. edited the manuscript. J.R.E., H.L., M.M.B., A.B., B.R., J.L., M.N. coordinated the research. J.R.E., M.M.B., C.L., C.O., M.N., A.P., J.K.O., J.L., R.G.C., J.R.N., J.A., M.K., W.T., A.B., H.L. generated the snmC-seq3 data. M.M.B., J.R.D., C.O., R.G.C., J.R.N., J.A., M.K., P.B., B.W., A.B., H.L. generated the sn-m3C-seq data. M.M.B., N.E., S.O., J.R., J.L., H.L., Q.Z., generated the MERFISH data. B.R., Y.E.L., S.Z. generated the snATAC-seq data. H.Z., K.A.S., B.T., Z.Y. generated the 10x scRNA-seq and SMART-seq data and the whole-mouse-brain transcriptomic cell type taxonomy and atlas. H.L., H.C. contributed to data archive/infrastructure. J.R.E., M.M.B. supervised the study.

## Methods

### Mouse brain tissues

All experimental procedures using live animals were approved by the Salk Institute Animal Care and Use Committee under protocol number 18-00006. Adult (P56) C57BL/6J male mice were purchased from Jackson Laboratories at seven weeks of age and maintained in the Salk animal barrier facility on 12-hour dark-light cycles with food ad-libitum for up to 10 days. Brains were extracted (between P56-P63), sliced, and dissected in an ice-cold dissection buffer as previously described^10^. For snmC-seq3 samples, brains were sliced coronally at 600 μm intervals from the frontal pole across the whole brain, yielding 18 slices, and dissected brain regions were obtained according to the Allen Brain Reference Atlas Common Coordinate Framework version 3^66^ (CCFv3, Extended Data Fig. 1a) For all the snm3C-seq samples, brains were sliced coronally at 1,200 μm, resulting in a total of 9 slices, and dissected 2-6 combined brain regions according to the CCFv3 (Extended Data Fig. 1b). For nuclei isolation, each dissected region was pooled from 3-30 animals, and 2-3 biological replicas were processed per region. Comprehensive brain dissection metadata can be found in Supplementary Table 1. Additionally, all dissected regions were digitally registered into CCFv3 using ITK-SNAP^67^ (v4.0.0) at a 25 μm resolution (Annotated voxel file available in the “Data Availability” section).

### Nuclei isolation and Fluorescence Activated Nuclei Sorting (FANS)

For snmC-seq3 samples, the nuclei were isolated and sorted into 384-well plates using previous methods^10^ with modifications described in Supplementary Information 1 (Section I, III). Briefly, single-nuclei were stained with AlexaFluor488-conjugated anti-NeuN antibody (MAB377X, Millipore) and Hoechst 33342 (62249, ThermoFisher) followed by FANS using a BD Influx sorter in single-cell (1 drop single) mode. For each 384-well plate, NeuN+ (488+) nuclei were sorted into columns 1-22, while NeuN-(488-) nuclei were sorted into columns 23-24, achieving an 11:1 ratio of NeuN+ to NeuN-nuclei (Supplementary Information 2). The snm3C-seq included additional in-situ 3C treatment steps during the nuclei preparation, allowing the chromatin conformation modality to be captured. These steps were performed using the Arima-3C BETA Kit (Arima Genomics), with a detailed protocol provided in Supplementary Information 1 (Section II).

### Library preparation and Illumina sequencing

Both snmC-seq3 and snm3C-seq samples followed the same library preparation protocol detailed in Supplementary Information 1. This protocol was automated using the Beckman Biomek i7 instrument to facilitate large-scale applications. The snmC-seq3 and snm3C-seq libraries were sequenced on an Illumina NovaSeq 6000 instrument, using one S4 flow cell per 16 384-well plates and employing a 150 bp paired-end mode.

### Mapping and primary quality control

The snmC-seq3 and snm3C-seq mapping was conducted using the YAP pipeline (cemba-data package, v1.6.8), as previously described^10^. Specifically, the main mapping steps included (1) demultiplexing FASTQ files into single cells (cutadapt^68^, v2.10); (2) read level quality control (QC); (3) mapping (one-pass mapping for snmC, two-pass mapping for snm3C) (bismark^69^, v0.20, bowtie2^70^, v2.3); (4) BAM file processing and QC (samtools^71^, v1.9, Picard, v3.0.0); and (5) methylome profile generation (allcools, v1.0.8); (6) chromatin contact calling (snm3C-seq only). Snakemake^72^ pipeline files with detailed mapping steps are provided in the “Code availability” section. All reads were mapped to the mouse mm10 genome. The gene/transcript annotation used in this study was based on a modified version of the GENCODE vm23 GTF file generated by the BICCN consortium, in accordance with Yao et al.^7^

Primary quality control for DNA methylome cells was (1) overall mCCC level < 0.05; (2) overall mCH level < 0.2; (3) overall mCG level < 0.5; (4) total final reads > 500,000 and < 10,000,000; and (5) Bismarck mapping rate > 0.5. Note that the Mccc level estimates the upper bound of the cell-level bisulfite non-conversion rate. Additionally, we calculated lambda DNA spike-in methylation levels to estimate each sample’s non-conversion rate. All samples demonstrated a low non-conversion rate (< 0.01, Extended Data Fig 2i). We chose loose cutoffs for the primary filtering to prevent potential cell or cluster loss. The clustering-based quality control described below accessed potential doublets and low-quality cells. For the 3C modality in snm3C-seq cells, we also required cis-long-range contacts (two anchors > 2500 bp apart) > 50,000.

### Analysis infrastructures

The whole-brain dataset comprised nearly 0.5 million single-cell or 5,000 pseudo-bulk mC profiles and 0.2 million single-cell or 2,500 pseudo-bulk 3C profiles. The dataset size was much larger than previous bulk and single-cell studies on mC or 3C^1, 10^. To enable efficient whole-brain data analysis, we formatted the entire multidimensional epigenomic data into three primary tensor datasets and used them as inputs for analysis at two different stages.

The first stage was cellular analysis. We employed a cell-by-feature tensor called “Methylome Cell DataSet” (MCDS) to carry out methylome-based clustering and cross-modality integration, as illustrated in Figures 2-3. Here, we focused on individual cells with aggregated genomic features, such as kilobase chromosome bins and gene bodies. This analysis allowed us to aggregate single-cell profiles into pseudo-bulk levels by clustering and annotation. The pseudo-bulk merge increased genome coverage while eliminating the need to frequently access hundreds of terabytes of single-cell files in the subsequent analysis stage.

The second stage was genomic analysis, where we used a pseudo-bulk-by-base tensor for mC, called “Base-resolution DataSet” (BaseDS), and a pseudo-bulk-by-2D-genome tensor for 3C, termed “Cooler dataset” (CoolDS), to perform methylome and chromatin conformation analysis at flexible genomic resolutions, as depicted in Figures 4-6. These pseudo-bulk tensors were generated at cell-group (1000s profiles) and subclass (100s profiles) levels to support multiple cellular granularities required by different analyses.

The large tensor datasets were stored using the chunked and compressed Zarr format^73^, hosted within the object storage of the Google Cloud Platform. Data analysis was conducted using ALLCools^10^, Xarray^74^, and dask^75^ packages. To facilitate large-scale computation, the Snakemake package^72^ was employed to construct pipelines, while the SkyPilot package^76^ was utilized to set up cloud environments. Additionally, the ALLCools package (v1.0.8) was updated to perform methylation-based cellular and genomic analyses, and the scHiCluster^77^ package (v1.3.2) was updated for chromatin conformation analyses. In the data and code availability section, we provided these tensor storages and reproducibility-related details (package version, analysis notebook, and pipeline files). For simplicity, the description below focused mainly on key analysis steps and parameters.

### Methylome clustering analysis

After mapping, single-cell DNA methylome profiles of the snmC-seq and snm3C-seq datasets were stored in the “All Cytosine” (ALLC) format, a tab-separated table compressed and indexed by bgzip/tabix^78^. The “generate-dataset” command in the ALLCools package helped generate a methylome cell-by-feature tensor dataset (MCDS). We used non-overlapping chromosome 100Kb (chrom100k) bins of the mm10 genome to perform clustering analysis; gene body regions ±2 kb for clustering annotation and integration with the companion transcriptome dataset; non-overlapping chromosome 5Kb (chrom5k) bins for integration with the chromatin accessibility dataset. Details about the integration analysis are described in the following section.

#### Pre-clustering

We performed two iterative clustering analyses for both the snmC and snm3C datasets. The first was a four-round pre-clustering for quality control purposes. The pre-clusters defined in this round contained potential doublets or low-quality cells (corresponding to debris or debris clumps in sorting). We started with all cells passing the primary quality control filters and used the “plate-normalized cell coverage” (PNCC) metric to mark problematic pre-clusters. This metric was calculated using the final mC reads of each cell divided by the average final reads of cells from the same 384-well plate. We reasoned that cells at the same plate underwent all the library preparation steps inside the same PCR machine, sharing the closest batch conditions. We observed some pre-clusters aggregating cells mostly showing extreme PNCC value (<0.5 or >2 fold) compared to most other clusters, which is a hallmark of problematic cells (Extended Data Fig. 2i). For each pre-cluster, we performed a permutation-based statistical test to call this abnormality. First, we randomly sampled null population cells with the cluster size, stratified on sample composition 10,000 times. We then calculated p-values for the observed PNCC mean (two-tailed test, larger or smaller) and standard deviation (std, one-tailed test, larger) comparing to null PNCC mean and std distribution. After calculating the false-discovery rate by the Benjamini-Hochberg procedure^79^ (FDR for short), we marked pre-clusters as low-quality with (1) abs(log2(PNCC)) > 0.8 and (2) FDR < 0.01 (for mean or std). In total, 8,979 (2.77%) snmC and 737 (0.38%) snm3C cells were removed from further analysis.

#### Methylome clustering

We then performed iterative clustering using the DNA methylome to determine whole-brain cell clusters. For both the snmC and snm3C datasets, we performed four rounds of iteration with the mCH and mCG fractions of chrom100k matrices. The clustering analysis within each iteration was described in a previous study^10^. We also provided annotated Jupyter notebooks in the “Code availability section,” detailing the functions and parameters used in each step. Most functions were derived from the allcools^10^, scanpy^80^, and scikit-learn^81^ packages. In summary, a single iteration consisted of the following main steps:

(1) Basic feature filtering based on coverage and ENCODE blacklist^82^.

(2) Highly Variable Feature (HVF) selection.

(3) Generation of posterior chrom100k mCH and mCG fraction matrices, as used in the previous study and initially introduced by Smallwood et al.^83^

(4) Clustering with HVF and calculating Cluster Enriched Features (CEF) of the HVF clusters. This framework was adapted from the cytograph2^39^ package. We first performed clustering based on variable features and then used these clusters to select CEFs with stronger marker gene signatures of potential clusters. The concept of CEF was introduced by Zeisel et al.^84^. The CEF calling and permutation-based statistical tests were implemented in “ALLCools.clustering.cluster_enriched_features”, where we selected for hypo-methylated genes (corresponding to highly-expressed genes) in methylome clustering.

(5) Calculate principal components (PC) in the selected cell-by-CEF matrices and generate the t-SNE^85^ and UMAP^86^ embeddings for visualization. t-SNE was performed using the openTSNE^87^ package with procedures described in Kobak and Berens 2019^88^.

(6) Consensus clustering. We first performed Leiden clustering^89^ 200 times, using different random seeds. We then combined these result labels to establish preliminary cluster labels, typically larger than those derived from a single Leiden clustering due to its inherent randomness^89^. Following this, we trained predictive models in the principal component (PC) space to predict labels and compute the confusion matrix. Finally, we merged clusters with high similarity to minimize confusion. The cluster selection was guided by the R1 and R2 normalization applied to the confusion matrix, as outlined in the SCCAF package^90^.

The iterative process of training and merging continued until the model’s performance on withheld test data achieved a specified accuracy (0.95 for the first round, >0.9 for all subsequent rounds). The Leiden algorithm’s resolution parameter significantly influenced cluster number and randomness (i.e., variation in cluster membership as random seeds changed), so we employed relatively small resolution values during each clustering stage (0.25 for the first iteration, 0.2-0.5 for the remaining iterations; the Scanpy default is 1).

This approach substantially reduced randomness while also underestimating cluster numbers. However, during the four rounds of iteration, any under-split clusters were further delineated in subsequent rounds. This framework was incorporated in “ALLCools.clustering.ConsensusClustering”.

For each clustering round, we assessed whether a cluster required additional clustering in the next iteration based on two criteria: (1) the final prediction model accuracy exceeded 0.9, and (2) the cluster size surpassed 20. In total, we executed four iterative clustering rounds, yielding the following cluster numbers: 61 (L1), 411 (L2), 1,346 (L3), and 2,573 (L4). We further separated cells within L4 clusters in the final round by considering their brain dissection region metadata. We first divided all dissection regions with more than 20 cells in an L4 cluster while combining other regions with fewer than 20 cells with their nearest regions based on average Euclidean distance in the PC space of L4 clustering. The final 4,673 cell groups combined L4 clusters and dissection regions. Incorporating dissection region data, which offered insights into a cell’s physical location, enhanced the analysis’s flexibility, such as enabling spatial region comparisons. Furthermore, we acknowledged that generating pseudo-bulk profiles from cell-level data demanded substantial computational resources. Aggregating cells at a higher granularity initially facilitated more straightforward merging later, such as combining them at the subclass level during subsequent analyses.

### Cluster-level DNA methylome analysis

After clustering analysis, we merged the single-cell ALLC files into pseudo-bulk level using the “allcools merge-allc” command. Next, we used the “allcools generate-base-ds” to generate the BaseDS from multiple ALLC files. The BaseDS was a Zarr dataset storing sample-by-base tensors for the entire dataset and allowed querying cytosines by genome position and methylation context (CpG, CpH). Next, we performed DMR calling as previously described^10, 22, 91^ using the “ALLCools.dmr.call_dms_from_base_ds” and “ALLCools.dmr.DMSAggregate” function that was reimplemented for BaseDS. In brief, we first calculated CpG differential methylated sites (DMS) using a permutation-based root mean square test^91^. The base calls of each pair of CpG sites were combined before analysis. We then merged the DMS into DMR if they were (1) within 250 bp and (2) having PCC > 0.3 across samples. Because the genome coverage was unbalanced between samples, we proportionally downsampled the coverage at each base in each sample to base call coverage (*cov*) of 50 and a total *cov* across samples of 3,000.

We applied the DMR calling framework across subclasses of the whole mouse brain and cell clusters within each major region. The two sources of DMRs were combined to capture the CpG fraction diversity in different cell-type granularities. There were around 10 million unique yet overlapping DMRs after the combination. We then merged the DMRs to get a final non-overlapping DMR list (“bedtools merge -d 0”), which included 2.56 million DMRs. We reported all the overlapping DMRs and non-overlapping DMRs in the “Data Availability” section. In the following analysis, when DMR was used to calculate correlation or scan motif occurrence, we started with the 10M overlapping DMRs. We selected the DMR with the strongest value (i.e., most significant PCC or highest motif score) among the overlapping ones. The DMRs in the final results were nonoverlapping.

### Atlas-level data integration and cluster annotation

We established a highly efficient framework based on the Seurat R package^31^ integration algorithm to perform atlas-level data integration with millions of cells. The integration framework consisted of 3 major steps to align two datasets onto the same space: (1) Using dimension reduction to derive embedding of the two datasets in the same space; (2) using canonical correlation analysis (CCA) to capture the shared variance across cells between datasets and find anchors as five mutual nearest neighbors (MNN) between the two datasets; (3) aligning the low-dimensional representation of the two datasets together with the anchors. We used genes to integrate methylome and transcriptome; chrom5k bins to integrate methylome and chromatin accessibility profiles.

#### Integrate methylome and transcriptome

To integrate our snmC-seq dataset with scRNA-seq data^7^, the gene expression levels of RNA cells were normalized by dividing the total UMI count of the cell and multiplying the average total UMI count of all cells and then log-transformed. For mC cells, the posterior gene-body mC level was used. The cluster-enriched genes (CEGs, similar to CEF described above) were identified in each cell subclass and cluster using mC data. We checked the variance of the mC CEGs among the mC cells and RNA cells and only used the CEGs with mC variance > 0.05 and expression variation > 0.005 for the analyses. We reversed the sign of mC levels before integration due to the negative correlation between gene body DNA methylation and gene expression (Fig. 1d). We fit a PCA model with the mC cells and transformed the RNA cells with the model. The PCs were normalized by the singular value of each dimension to avoid the embedding being driven by the first few PCs.

To find anchors between mC and RNA cells, we first Z-score scaled the mC matrix and expression matrix of CEGs across cells, and the resulting matrices were represented as *X* (mC: cell-by-CEG) and *Y* (RNA: cell-by-CEG), respectively. CCA was used to find the shared low dimensional embedding of the two datasets, solved by singular value decomposition (SVD) of their dot product *USV^T^ = XY^T^*. *U* and V were normalized by dividing the L2-norm of each row, and were used to find five MNNs as anchors and scored anchors using the same method as Seurat^31^.

The original CCA framework of Seurat (v4) is hard to scale up to millions of cells due to the memory bottleneck, where the mC cell-by-RNA matrix was used as the input to CCA. To handle this limitation, we randomly selected 100,000 cells from each dataset (*X_ref_* and *Y_ref_*) as a reference to fit the CCA and transformed the other cells (*X_qry_* and *Y_qry_*) onto the same CC space. Specifically, the canonical correlation vectors (CCV) of *X_ref_* and *Y_ref_* (denoted as *U_ref_* and *V_ref_*) were computed by 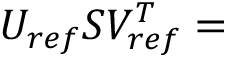 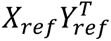, where 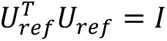 and 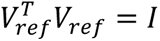. Then the CCV of *X_qry_* and *Y_qry_* (denoted as *U_qry_* and *V_qry_*) were computed by 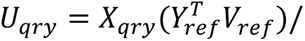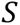 and 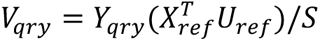. The embeddings from the reference and query cells were concatenated for anchor identification.

The PCs derived from the first step were then integrated using the same method as Seurat^31^ through these anchors. Rather than working on the raw feature space in Seurat, our integration step projected the PCs of scRNA-seq (query, denoted as *Ur)* to the PCs of the snmC-seq (reference, denoted as *Um*) while keeping the PCs of the reference dataset unchanged. This approximation considerably reduced the time and memory consumption for computing the corrected high-dimensional matrix and redoing the dimension reduction. For anchor *k* pairing mC cell *km* and RNA cell *kr*, *B_k_ = Um_km_ – Ur_kr_* was considered the bias vector between mC and RNA. Then for each RNA cell as a query, we used its 100 nearest anchors to compute a weighted average bias vector representing the distance to move an RNA cell into the mC space. The distance between the query RNA cell and an anchor was defined as the Euclidean distance on the RNA dimension reduction space between the query RNA cell and the RNA cell of the anchor. The weights for the average bias vector depended on the distances between the query RNA cell and the anchors, where close anchors received high weights.

#### Integrate methylome and chromatin accessibility profiles

PCA on gene body signals was insufficient to capture the open chromatin heterogeneity in snATAC-seq data^11, 31^. Latent Semantic Indexing (LSI) applied to binarized cell-by-5kb bin matrices had demonstrated promising results for snATAC-seq data embedding and clustering^31^. Therefore, to align snATAC-seq data with snmC-seq data at a high resolution, we developed an extended framework based on the previously described approach to utilize binary sparse cell-by-5kb bin matrices as input.

We first derived a cell-by-5kb bin matrix to represent the snmC-seq data. In a single cell *i*, we modeled its mCG base call *M_ij_* for a 5 kb bin *_j_* using a binomial distribution *M_ij_ ∼ Bi*(*cov_ij_,p_i_*), where *p* represented the global mCG level of the cell. We then computed *P(M_ij_ > mc_ij_)* as the hypomethylation score of cell *i* at bin *j*. The less likely to observe smaller or equal methylated base calls, the more hypomethylated the bin was. We next binarized the hypomethylation score matrix by setting the values greater than 0.95 as 1, otherwise 0, to generate a sparse binary matrix *A*. We selected the columns with more than five non-zero values, then computed the column sum of the matrix 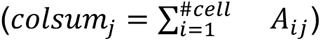 and kept only the bins with Z-scored *log*_2_*colsum* between -2 and 2. The snATAC-seq data was also represented in a binary cell-by-5kb bin matrix, where 1 represented at least one read detected in a 5 kb bin in a cell. The features were filtered in the same way as the mC matrix, and the bins remaining in both datasets were used for further analysis.

LSI with log term frequency was used to compute the embedding. Term Frequency-Inverse Document frequency (TF-IDF) transformation was applied to convert the filtered matrix B to *X*. Specifically, *B* was normalized by dividing the row sum of the matrix to generate the term frequency matrix *TF,* and further converted to *X* by multiply the inverse document frequency vector *IDF*.

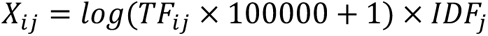

, where 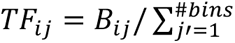 and 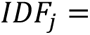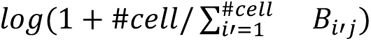. The embedding of single cells *U* was then computed by SVD of *X*, where *X* = *USV^T^.* We fit the LSI model with mCG data *Bm* to derive *Um.* The intermediate matrices *S* and *V* and vector *IDF* were used to transform the ATAC data *Ba* to *Ua,* by CCA was also performed with the downsampling framework using 100,000 cells from each dataset as reference and the others as query, but taking the TF-IDF transformed matrices as input. The query cells were projected to the same space using the IDF and CCV of the reference cells. Specifically,

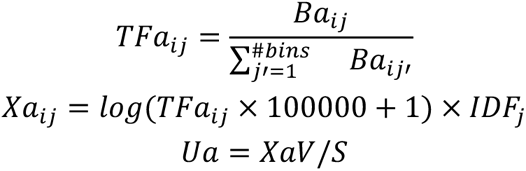

*Bm_ref_* and *Ba_ref_*were converted to *Xm_ref_*and *Xa_ref_* with TF-IDF, and the CCVs (denoted as *U_ref_*and *V_ref_*) were computed by 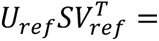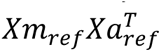. Then *Bm_qry_* and *Ba_qry_* were converted to *Xm_qry_* and *Xa_qry_* with TF-IDF using the IDF of reference cells, and the CCVs (denoted as *U_qry_* and *V_qry_*) were computed by 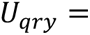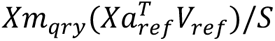 and 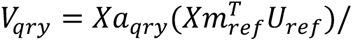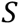. The following steps to find anchors and align *Um* and *Ua* were the same as integrating the mC and RNA data.

#### Iterative integration group design

Like clustering analysis, we integrated two datasets iteratively to match cell or cell clusters at the highest granularity. We first separated the pass-QC datasets into integration groups based on independent cell type annotation (described above or provided by data generators) and dissection information. For instance, non-neuronal cells, IMN, and granule cells (“DG Glut” and “CB Granule Glut”) were separated from neurons because they were (1) showing large global methylation differences from other neurons and (2) unbalanced in cell numbers across datasets due to different sampling and sorting strategies. Within each integration group, we performed the integration iteratively. We used the co-clustering from the integrated low-dimensional space to match cells or clusters between the two datasets (see below). We then performed the next round of integration until the matched cells or clusters fulfilled the stopping criteria. We listed details about each pair of iterative integration below. The resulting cluster map between datasets and mC/m3C cluster annotation was provided in Supplementary Table 4. A set of Jupyter Notebooks for a single integration process between each pair was provided in the “Code Availability” section.

#### Integration between snmC-seq and scRNA-seq or SMART-seq dataset

We used the gene body ± 2kb as features to integrate mC and RNA datasets^7^, mapping the RNA clusters to mC cell groups. We used the mCG fraction of the gene bodies for non-neuronal cells, IMN, and granule cells and the mCH fraction of the gene bodies for other neurons. In each iteration, we calculated a confusion matrix between 4,673 mC cell groups and 5,200 RNA clusters (provided by data generators) using the overlap score as previously described^10, 92^. We then built a weighted graph using the confusion matrix as the adjacency matrix and performed a Leiden clustering (resolution=1) to bicluster mC and RNA clusters. This step puts similar mC and RNA clusters into integration groups based on their overlap score. The RNA and mC clusters in the same integration group were further integrated to match at finer granularity in the next iteration unless any stop criteria were met: (1) there was only one integration group from this round; (2) there was only one mC or RNA cluster in the integration group; (3) the mC cell number < 30; (4) RNA cell number < 100 for the scRNA-seq dataset or < 30 for the SMART-seq dataset. After integration, we obtained an mC to RNA cluster map for each mC cell group, which we used as the reference to annotate cell subclasses and remaining hierarchies in the transcriptomic taxonomy. We also evaluated the spatial location and marker genes (neurotransmitter-related genes or other markers provided in the transcriptome annotation). We resolved conflicts manually when the RNA clusters corresponded to more than one subclass by checking the dissection metadata and marker genes. We combined all RNA cells assigned to each mC cell group to generate the matched transcriptome profile.

#### Integration between snmC-seq and snATAC-seq dataset

The snmC-seq dataset and snATAC dataset^12^ shared the same dissection tissues. We utilized this experiment design to integrate mC and ATAC cells within each major region. Besides, the snmC-seq data was enriched for NeuN+ by FANS, while the snATAC data unbiasedly profiled all cells. Therefore, we also separated neurons with non-neuronal cells and IMN to balance the integration, especially in the first round. We used the mCG hypo-methylation score of chromosome non-overlapping 5kb bins to perform the integration. The cluster assignment and stop criteria were similar to the mC-RNA integration. The alignment score (Extended Data Fig. 6a) is calculated as previously described^93^, using K=1% cells of the dissection region or k=20, whichever is larger. We combined all ATAC cells assigned to each mC cell group to generate the matched chromatin accessibility profile.

#### Integration between snmC-seq and snm3C-seq dataset

We used the non-overlapping chromosome 100kb bin as features to integrate snmC-seq and snm3C-seq datasets. The cluster assignment and stop criteria were similar to the mC-RNA integration. After integration, we also annotated the snm3C cell groups with the transcriptomic taxonomy.

### MERFISH Experiment

#### MERFISH gene panel design

The genes in the GTF file were first filtered based on length > 1kb. We then selected genes based on the Zhang et al.^33^ methods but used the snmC-seq dataset and gene body mCH fraction to perform the calculation. In brief, there are two approaches to prioritizing genes. The first approach was to use mutual information between gene body mCH fraction, and neuron subclasses labels, which aims to select genes differentially methylated between groups of cell subclasses. The second approach was to perform pairwise differentially methylated gene analysis (ALLCools.clustering.PairwiseDMG) among clusters within the same major region and select genes being identified as DMGs in most cluster pairs. For the first approach, we selected the top 100 genes. We selected the top 50 genes from each major region for the second approach. Due to the overlaps, there were 325 genes after this selection. In addition to the cell-type markers, we also selected spatial markers by calculating the mutual information between the cell’s major region label and mCH fraction across the brain; or between the dissection region label and mCH fraction within each major region. We added another 175 non-overlapping genes to a total of 550 genes. We then performed the same analysis using the scRNA-seq dataset from Yao et al.^7^ to get the RNA-based prioritization lists. We selected 500 final genes as the gene panel based on rank in the RNA list to ensure these genes are also expressed and highly diverse in the transcriptome. Encoding probes for these genes were designed and synthesized by the Vizgen company (Supplementary Table 6).

#### MERFISH tissue preparation and imaging

Fresh P56-63 whole mouse brains were sliced coronally at 1,200 μm intervals, and each slice was then embedded in OCT, rapidly frozen in isopentane/dry ice, and stored at -80 C° until ready for slicing. 12-μm-thick coronal sections were obtained from each OCT-blocked tissue using a Leica CM1950 cryostat, immediately fixed in 4% formalin (warmed to 37 °C) for 30 min, and permeabilized in 70% ethanol following manufacturer procedures. Sample preparation, including probe hybridization and gel embedding, was performed using Vizgen’s sample preparation kit (Vizgen:10400012) following the manufacturer’s protocol. Each section was imaged using MERSCOPE 500 Gene Imaging Kit (Vizgen:10400006) on a MERSCOPE (Vizgen).

#### MERFISH data preprocessing and annotation

MERFISH data analysis, including imaging, spot detection, cell segmentation, and cell-by-gene matrix generation, was conducted by the MERSCOPE Instrument Software. We removed abnormal cells (artificial segmentation, doublets) from the cell-by-gene matrix in each experiment: (1) Cell volume < 30μm^3^ or > 2000μm^3^; (2) total RNA counts < 10 or > 4000; (3) total RNA counts normalized by cell volume < 0.05 or > 5; (6) total gene detected < 3; (5) cells with > 5 blank probes detected (negative control probe included in the gene panel). We then integrated the pass-QC MERFISH cells with the scRNA-seq datasets^7^ to annotate the MERFISH cells with transcriptome nomenclatures using the ALLCools integration functions described above.

#### Integration between MERFISH and snmC and snm3C dataset

We integrated the snmC and snm3C datasets with the MERFISH dataset to evaluate whether the spatial pattern observed in the DNA methylome matched the spatial diversity observed in the gene expression data. Integration was similar to the mC-RNA integration described above. To utilize the dissection region metadata, we grouped the snmC-seq and snm3C-seq data by the slice and integrated them with a matched MERFISH slice. We also separated neurons and other cells, similar to the mC-RNA integration above. We used the 500 genes in the MERFISH gene panel to perform the integration. After integration, we imputed the spatial location of each methylation nucleus on the integrated low-dimensional space. We calculated the ten nearest MERFISH neighbors for each mC nucleus in each integration group. We assigned the coordinate of these MERFISH cells’ centroids as the mC nucleus’s spatial location.

### Cell and cluster-level chromatin conformation analysis

#### Generate chromatin contact matrix and imputation

After snm3C-seq mapping, we used the cis-long range contacts (contact anchors distance > 2,500 bp) and trans contacts to generate single-cell raw chromatin contact matrices at three genome resolutions: chromosome 100-Kb resolution for the chromatin compartment analysis; 25-Kb bin resolution for the chromatin domain boundary analysis; 10-Kb resolution for the chromatin interaction analysis. The raw cell-level contact matrices were saved in the scool format^94^. We then used the scHiCluster package (v1.3.2) to perform contact matrix imputation as described previously^77^. In brief, the scHiCluster imputed the sparse single-cell matrix by first performing a Gaussian convolution (pad=1) followed by using a random work with restart algorithm on the convoluted matrix. For 100-Kb matrices, the whole chromosome was imputed; for 25-Kb matrices, we imputed contacts within 10.05Mb; for 10-Kb matrices, we imputed contacts with 5.05Mb. The imputed matrices for each cell were stored in cool format^94^. The cell matrices were aggregated into cell groups or subclass levels identified in the previous section. These pseudo-bulk matrices were concatenated into a tensor called CoolDS and stored in Zarr format for brain-wide analysis.

#### Compartment analysis

We used the imputed subclass-level contact matrices at the 100-Kb resolution to analyze the compartment. We first filtered out the 100kb bins that overlapped with ENCODE blacklist v2^82^ or showed abnormal coverage. Specifically, the coverage of bin i on chromosome c (denoted as R_c,i_) was defined as the sum of the i-th row of the contact matrix of chromosome c. We only kept the bins with coverage between the 99th percentile of R_c_ and twice the median of R_c_ minus the 99th percentile of R_c_. Contact matrices were normalized by distance, and the PCC of the normalized matrices was used to perform the Principal Component Analysis (PCA) (Aiden 2009 Science). The IncrementalPCA class from the sklearn package^81^, which allows fitting the model incrementally, was used to fit a single PCA model incrementally for each chromosome using all the cell subclass matrices. We then transformed all the cell subclasses with the fitted model, so the PCs for each subclass were transformed from the same loading and eased the cross-sample correlation analysis. We also calculated the correlation between PC1 or PC2 and 100-Kb bin CpG or gene density. We use the component with higher absolute correlation as the compartment score and assign the compartment with higher CpG density with positive scores (A compartment).

#### Compartment score and mC fraction correlation

We first performed quantile normalization along subclasses using the Python package qnorm (v0.8.0)^95^ to normalize the mC fractions and compartment scores. We then calculated the PCC between the compartment scores of non-overlapping chromosome 100Kb bins with the corresponding bin’s mCH or mCG fractions across cell subclasses. Because the negatively correlated bins’ compartment score had a much higher standard deviation among cell types (Fig. 4c), we selected the 300 most negatively correlated chrom100k bins and used their overlapped genes to perform gene ontology (GO) enrichment analysis (Fig. 4d) using Enrichr^38^. We randomly selected gene-length matched background genes to adjust the long-gene bias in all the GO enrichment analyses^38^. To investigate the developmental relevance indicated by the GO enrichment result, we used the developmental mouse brain scRNA-seq atlas^39^ at the subtype level (approximate granularity of subclass in this study). Genes overlapping 300 most negatively correlated bins, 300 mostly positively correlated bins, and 300 low correlation bins were used to plot Extended Data Fig. 9d.

#### Domain boundary analysis

We used the imputed cell-level contact matrices at the 25-Kb resolution to identify the domain boundaries within each cell using the TopDom algorithm^96^. We first filtered out the boundaries that overlap with ENCODE blacklist v2^82^. The boundary probability of a bin was defined as the proportion of cells having the bin called a domain boundary among the total number of cells from the group/subclass. To identify differential domain boundaries between “n” cell subclasses, we derived an nx2 contingency table for each 25kb bin. The values in each row represent the number of cells from the group that has the bin called a boundary or not as a boundary. We computed each bin’s Chi-square statistic and p-value and used FDR <1e-3 as the cutoff for calling 25kb bins with differential boundary probability.

#### Domain boundary probability and transcript body mC fraction correlation

We first performed quantile normalization along subclasses using the Python package qnorm (v0.8.0)^95^ to normalize the transcript body mC fractions and chromosome 25Kb bin boundary probabilities. We then calculated the PCC between the differential boundary probabilities of 25Kb bins with the transcript body mCH and mCG fractions. We grouped transcripts with >90% overlap within a gene and used their longest range. We calculated the transcript-body mCH and mCG fraction at the subclass level for each transcript. We then calculated the PCC between the mC fractions and boundary probabilities of bins overlapping the transcript body ± 2 Mb. We used a permutation-based test to estimate the statistical significance of the correlation^97^. Specifically, we shuffled the boundary probability and mC fraction values within each sample (subclass), disrupting the genome relationship between the bins while preserving the sample-level global difference. We calculated the PCC using the shuffled matrices 100,000 times and used a normal distribution to approximate the null distribution for more precise p-value estimation in FDR correction. We then used FDR < 1e-3 as the significance cutoff for each PCC between a transcript and a 25Kb bin. In Fig. 2g, we used deeptools^98^ (v3.5.1) to profile the boundary probability at transcript ± 2 Mb 25Kb bins. In Fig. 2h and Extended Data Fig. 2f-h, we selected the top positively correlated bin and top negatively correlated bin for each long gene (transcript body length > 100Kb) and performed the GO analysis using length-matched background genes, as described above (Extended Data Fig. 2h).

#### Highly variable interaction analysis

We used the imputed cell-level contact at the 10-Kb resolution to perform the highly variable interaction analysis, where the interaction represented one 10Kb-by-10Kb pixel in the conformation matrix. We filtered out any interactions that had one of the anchors overlapping with ENCODE blacklist v2^82^. We then performed a one-way analysis of variance (ANOVA) for each interaction to test whether the single-cell contact strength of that interaction displayed significant variance across subclasses. The F statistics of ANOVA represented an overall variability of the interaction across the brain. To select highly variable contacts, we used F > 3 as the cutoff, which was decided by visually inspecting the contact maps as well as fulfilling the FDR < 0.001 criteria. The ANOVA analysis was only performed on interactions whose anchor distance was between 50 Kb and 5 Mb, given that increasing the distance only led to a limited increase in the number of significantly variable and gene-correlated interactions (Extended Data Fig. 10b).

#### Interaction strength and mC fraction correlation

To investigate the relationship between gene status and the surrounding chromatin conformation diversity, we first performed quantile normalization along subclasses using the Python package qnorm (v0.8.0)^95^ to normalize the transcript body mCH fractions and contact strengths of highly variable interactions. We then calculated PCC between the transcript body mCH fraction and the highly variable interactions if any anchor of the interactions had overlapped with the gene body. Similar to the domain boundary correlation analysis, we shuffled the contact strengths and mCH fractions within each sample and used the shuffled matrix to calculate null distribution and estimate FDR. We select FDR < 0.001 as a significant correlation.

### Gene Regulatory Network (GRN) analysis

We presented a framework for building GRN based on the DNA methylome and chromatin conformation profiles at the cell subclass level. We used 212 neuronal cell subclasses requiring them to have> 100 cells in both snmC and snm3C datasets. Notably, the same framework can be applied to other brain cell types or a subset of cells (such as certain brain regions or cell classes based on specific questions). The GRN was composed of relationships between TFs, their potential binding elements (represented by DMRs), and downstream target genes. Pairwise edges were constructed between DMRs and target genes (DMR-Target), TFs and target genes (TF-Target), and TFs and DMRs (TF-DMR). The basis of each pairwise edge was the correlation between the methylation fractions of the two genome elements across cell subclasses. We performed quantile normalization along subclasses using the Python package qnorm (v0.8.0)^95^ to normalize the two matrices involved in calculating the correlation. Gene body mCH fraction was used as a proxy for TF and target gene activity, and mCG fractions were used to represent DMR status. Variable genes and TFs were selected if they were identified as CEFs (described in the clustering steps) in any subclass.

For the DMR-Target edges, we selected the highly variable and positively correlated chromatin contact interactions of the gene based on the results in the previous section and included DMRs situated in any anchor regions of the interactions. We then calculated the PCC between DMR mCG and gene mCH fraction. For a group of overlapping DMRs, we selected the one with the highest absolute PCC value to represent that group, making the edges’ DMRs non-overlap. Similar to the domain boundary and interaction correlation analysis, we shuffled the DMRs and genes within each sample to calculate null PCC and estimate FDR. We filtered DMR-Target edges with FDR < 0.001. For the TF-Target and TF-DMR edges, we calculated the PCC between TF and all CEF genes or between TF and all DMRs, respectively, and applied the same FDR < 0.001 cutoff to filter edges. For the TF-DMR edge, we further performed motif enrichment analysis on the significantly correlated DMRs (explained in the next section). We only keep TF-DMR edges when the TF has any motif significantly enriched in the correlated DMR set, and the particular DMR has that motif occurrence.

After getting the three pairwise edges, we intersected the edges together into triples based on shared genes (including TFs and targets) and DMR ids. We calculated a final edge score 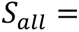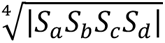 for each triple by taking the geometric mean of the absolute values of four correlations, where *S_a_* was the correlation of DMR-Target edge, *S_b_* was the correlation of TF-DMR edge, *S_c_* was the TF-Target edge, and *S_d_* was the correlation between target gene mCH fraction and gene-DMR contact strength. If multiple gene-correlated interactions have anchors overlapping with DMR and gene body, we select the one with the lowest negative correlation.

#### DMR motif scan and TF motif enrichment analysis

We used an ensemble motif database from SCENIC+^44^, which contained 49,504 motif position weight matrices (PWM) collected from 29 sources. Redundant motifs (highly similar PWMs) were combined into 8,045 motif clusters through clustering based on PWM distances calculated by TOMTOM^99^ by the SCENIC+ authors^44^. Each motif cluster was annotated with one or more mouse TF genes. To calculate motif occurrence on DMR regions, we used the Cluster-Buster^100^ implementation in SCENIC+, which scanned motifs in the same cluster together with Hidden Markov Models.

To perform motif enrichment analysis in the “TF-DMR edge” analysis, we used the recovery-curve-based cisTarget algorithm^44, 65^. In brief, the cisTarget algorithm performed motif enrichment on a set of DMRs by calculating a Normalized Enrichment Score (NES) for each motif based on all other motifs in the collection. For each TF gene, we applied the cisTarget algorithm to positively correlated or negatively correlated DMRs separately. We used the package default cutoff (NES > 3) to select enriched motifs for a DMR set. A leading-edge analysis was performed by cisTarget to assign motif occurrence in DMRs with Cluster-Buster scores passing a cutoff in enriched cases^44^.

#### PageRank analysis on weighted networks

We adopted the Taiji framework^51^ to perform TF analysis on weighted GRN for each cell subclass. This framework employed the personalized PageRank algorithm^101^ to propagate node and edge weight information across the network, calculating the importance of each TF. To add subclass information as network weights, we simplified the network by only including TF and target gene nodes, and weighing the gene node by inverted gene body mCH value in the subclass. Specifically, we first performed quantile normalization across all subclasses. We then performed a robust scale of the matrix using “sklearn.preprocessing.RobustScaler” with quantile_range=(0.1, 0.9). We then inverted the scaled mCH fraction by

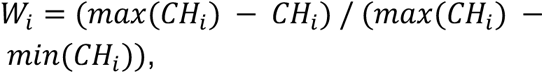

where *CH_i_* and *W_i_* denoted the scaled gene mCH fractions and inverted values for subclass *i*, respectively.

We also added DMR mCG fraction into the edge weights. Specifically, we performed the same quantile normalization and robust scale on all the DMRs’ mCG fractions involved in the network and calculated the inverted DMR mCG value by

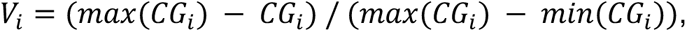

where *CG_i_* and *V_i_* denoted the scaled DMR mCG fractions and inverted values for subclass *i*, respectively. The edge weight between a TF and a target gene in subclass *i* was calculated as *e =* 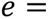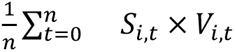, where *n* denoted the number of DMRs that connecting the TF to target gene, *S_i,t_* was the final score of one TF-DMR-Target triple, *V_i,t_* was the inverted DMR mCG value.

### Intragenic epigenetic and transcriptomic Isoform Analysis

#### Integration and isoform quantification of the SMART-seq dataset

Preprocessing and gene level quantification via STAR^102^ (v2.7.10) was performed by AIBS data generators as previously described^6^. We used gene-level counts to perform cross-modality integration iteratively as described in previous sections. We used kallisto^103^ with steps described in a previous study^27^ to quantify the SMART-seq at the isoform level with the same GTF file used in transcriptome and methylome analysis above. We calculated cell-group-level transcript per million (TPM) based on the integration result. We also calculated the exon Percent Spliced In (PSI) from the transcript counts in each gene. The SMART-seq browser tracks (Fig. 6f, g) were constructed from STAR-aligned BAM files.

#### Prediction model training

First, we quantified mC and m3C intragenic features for predicting the alternative isoform and exon usage. We used the exon, exon-flanking region, and intragenic DMRs as the mC features of each gene. The exon flanking region was defined as upstream or downstream 300 bp of each exon. We removed features with variance < 0.01, and combined features with > 90% overlap in their genome coordinates. For 3C features, we used all the intragenic highly variable interactions (f statistics > 3) from the above section as features.

After collecting all the features, we selected genes with highly variable transcripts and exons among cell groups for model training. Highly variable transcripts were selected based on: (1) mean TPM across cell groups > 0.2; (2) TPM standard deviation > 0.3; (3) transcript body (TSS to TTS) length > 30Kb. Highly variable exons were selected based on: (1) PSI standard deviation > 0.02; (2) PSI 90% quantile and 10% quantile difference > 0.05. We trained four models for each gene including predicting transcripts TPMs using mC or 3C features and predicting exon PSIs using mC or 3C features. The training contains two steps: first, we used “sklearn.feature_selection.SelectKBest” with the score function “f_regression” to select the top 100 features for each transcript or exon. We then used all features that had been selected at least once. We performed five-fold cross-validation to train random forest models using selected features and “sklearn.ensemble.RandomForestRegressor”. In each cross-validation run, we calculated the PCC between predicted values and true values as the model performance. We also shuffled the selected features within each sample (Fig. 6c) to train the model and calculate PCC again as the shuffled model performance.

### Data Availability

The snmC-seq2/3 single-cell sequencing data are accessible through Neuroscience Multi-omic Data (NeMO) Archive https://tinyurl.com/cembanemo. The snm3C-seq single-cell sequencing data will be accessible through NeMO and GEO. The MERFISH dataset will be accessible through GEO. The whole-brain snATAC-seq dataset is shared by Zu et al^12^. The whole-brain scRNA-seq and SMART-seq dataset is shared by Yao et al^7^. All the processed data related to results and method sections are shared in this GitHub repository: https://github.com/lhqing/wmb2023.

### Code Availability

Mapping pipeline for snmC-seq3 and snm3C-seq is available at https://hq-1.gitbook.io/mc/. Single-cell DNA methylome data analysis tools are available at ALLCools (v1.0.8) python package, https://lhqing.github.io/ALLCools/intro.html; Single-cell chromatin conformation data analysis tools are available at the scHiCluster (v1.3.2) python package, https://github.com/zhoujt1994/scHiCluster. Other codes and Jupyter Notebooks related to results and method sections are shared in this GitHub repository: https://github.com/lhqing/wmb2023.

**Extended Data Figure 1.**
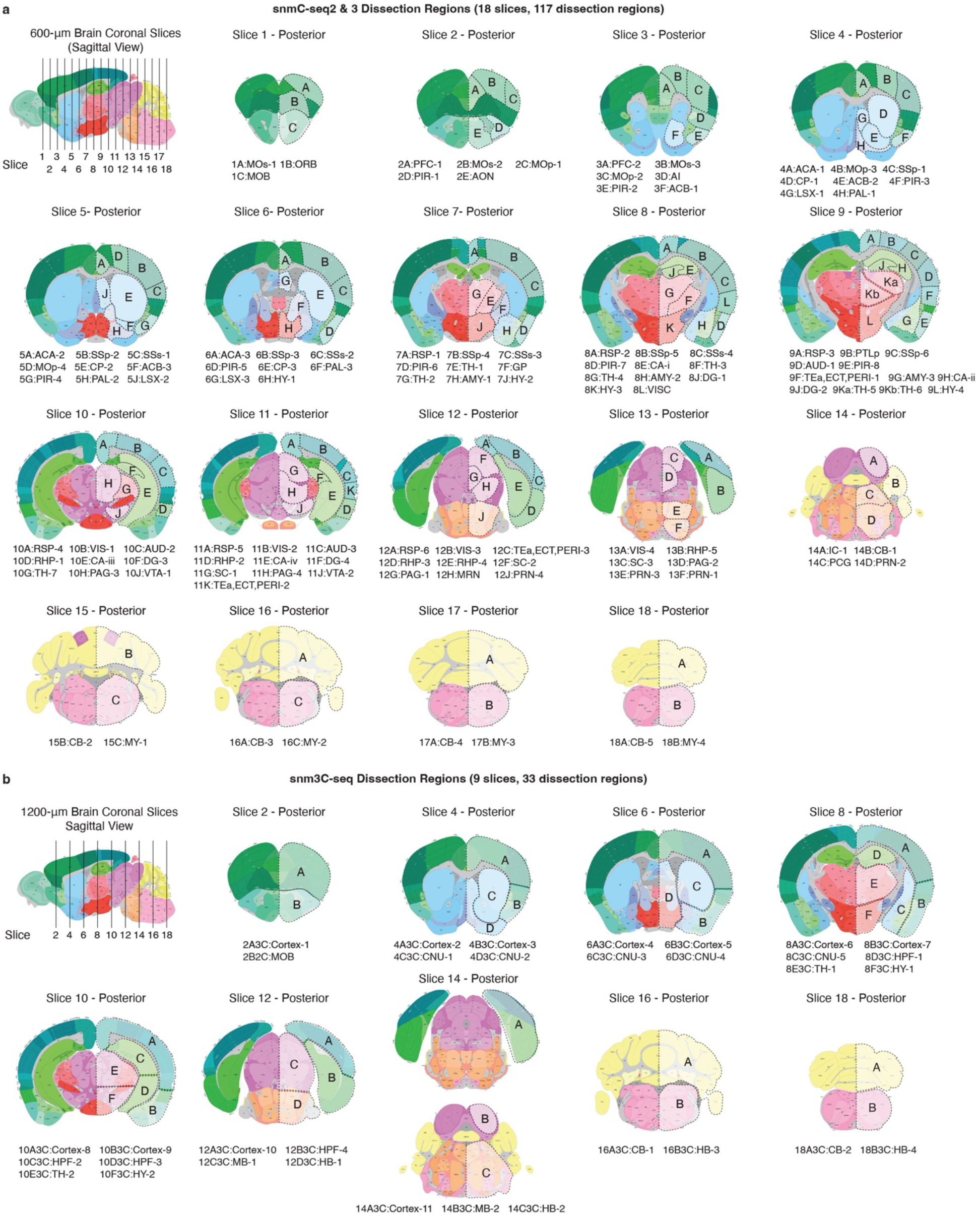
Brain dissection regions. Schematic of brain dissection steps. Each male C57BL/6 mouse brain (age P56) was dissected into 600-μm slices for snmC-seq3 **(a)** and 1,200-μm slices for snm3C-seq3 **(b)**. We then dissected brain regions from both hemispheres within a specific slice.

**Extended Data Figure 2.**
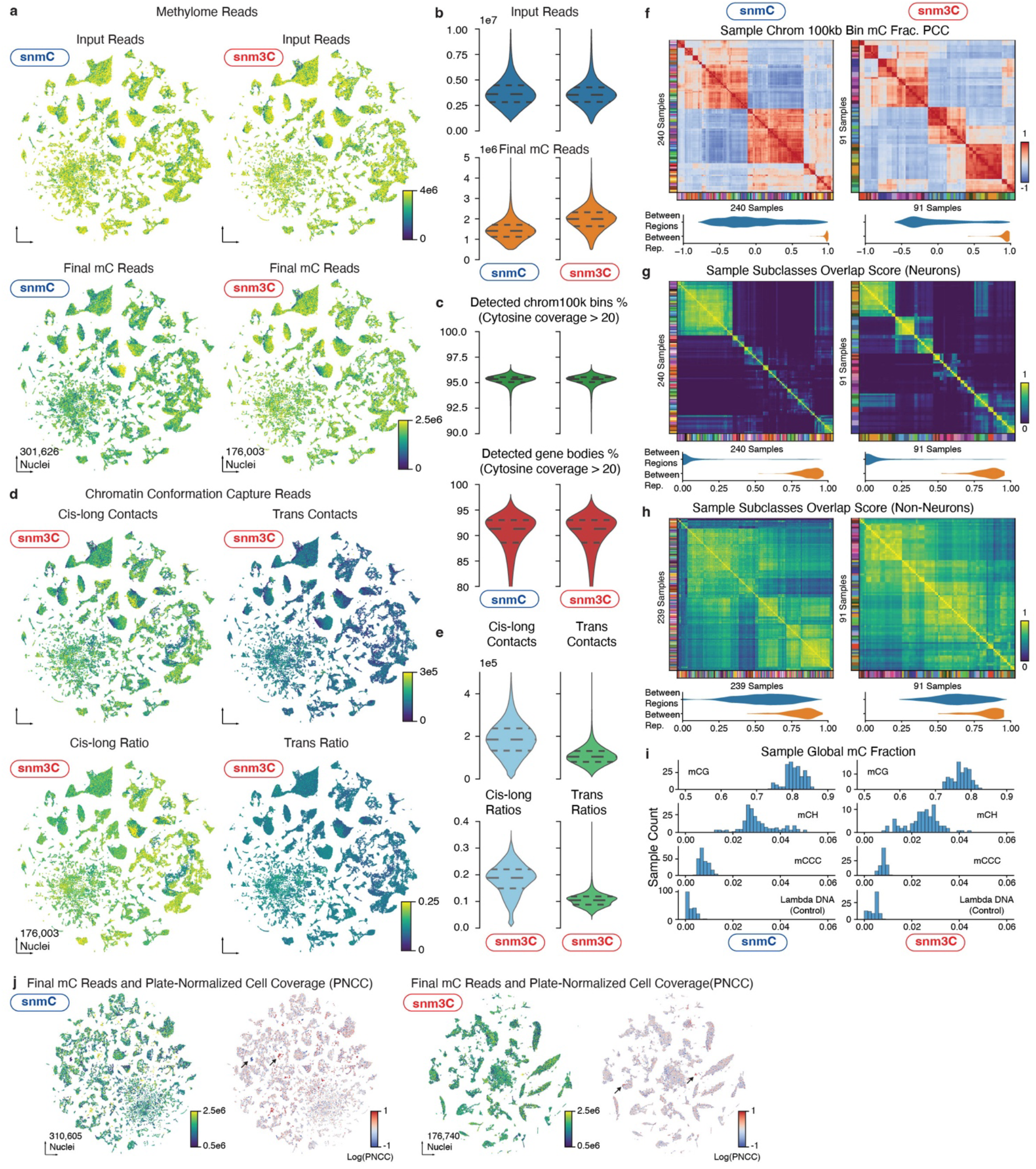
Quality Control for snmC and snm3C dataset. **a-b,** The number of input reads and final pass QC reads in snmC-seq3 and snm3C-seq shown by t-SNE **(a)** and violin plot **(b) c,** The percentage of chrom100k bins or genes detected per cell in snmC-seq3 and snm3C-seq. Gray lines from top to bottom indicate the 75%, 50%, and 25% quantiles. **d-e,** The number and ratio of cis-long and trans contacts in snm3C-seq, depicted by t-SNE **(d)** and violin plot **(e)**. **f,** Heatmap of PCC between the average methylome profiles (mean mCH and mCG fraction of all chromosome 100-kb bins across all cells belonging to a replicate sample). The violin plot below summarizes the values between replicates within the same brain region or between different brain regions. **g-h,** Pairwise overlap score (measuring co-clustering of two replicates) of neuronal subtypes and **(g)** non-neuronal subtypes **(h)**. The violin plots summarize the subtype overlap score between replicates within the same brain region or between different brain regions. **i,** Distribution of the mCG, mCH, mCCC, and Lambda DNA fraction (non-conversion rate) at sample level in snmC-seq3 and snm3C-seq. **j,** Pre-clustering t-SNE of snmC and snm3C dataset colored by final mC reads and plate-normalized cell coverage. Arrows indicate typical low-quality clusters filtered out from the further analysis.

**Extended Data Figure 3.**
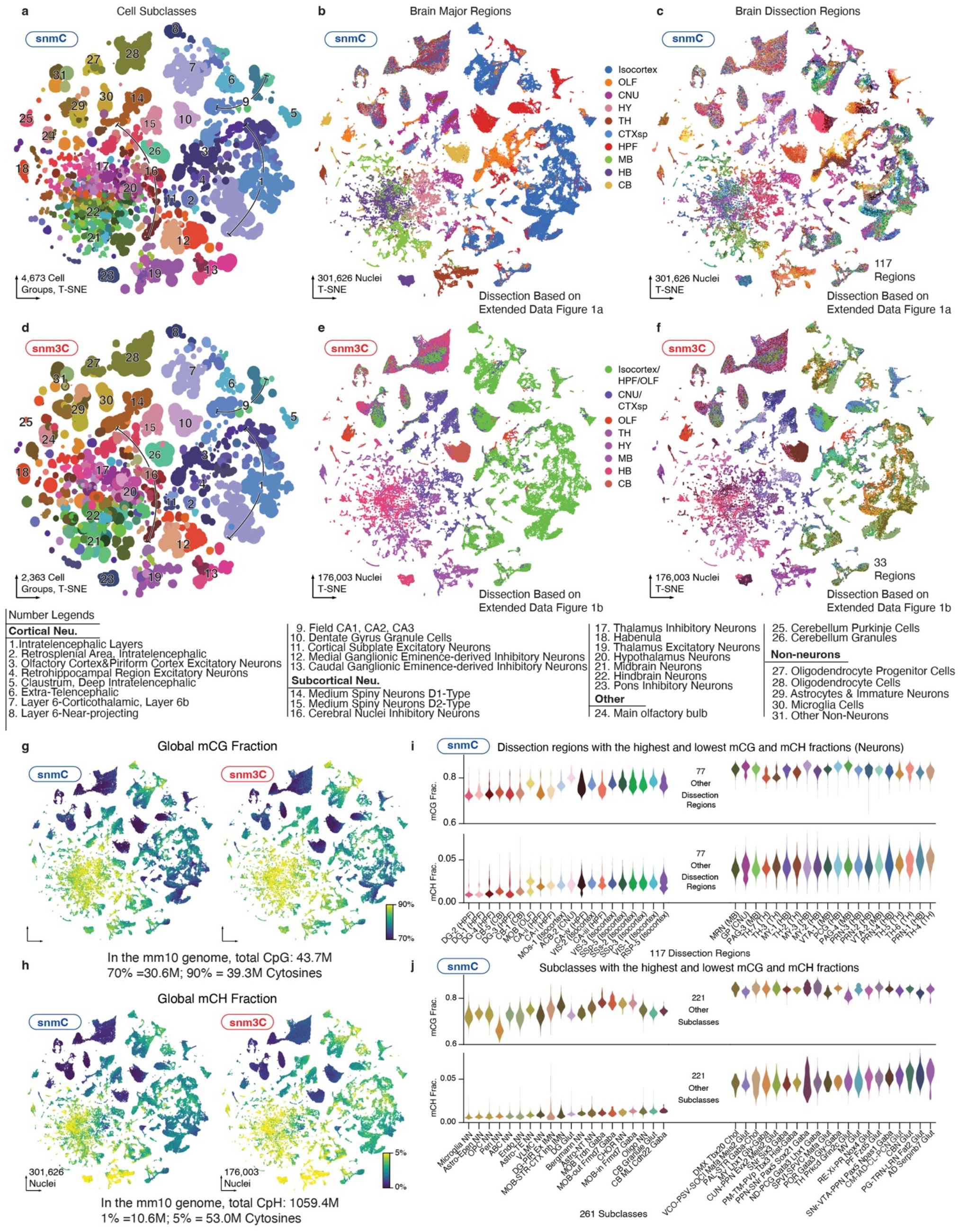
Metadata of snmC-seq and snm3C-seq dataset. **a-c**, t-SNE of snmC-seq color by cell subclass **(a)**, major regions **(b)**, and dissection regions **(c)**. **d-f**, t-SNE of snm3C-seq color by cell subclass **(d)**, major regions **(e)**, and dissection regions **(f)**. **g,h,** Cell-level t-SNE of snmC-seq and snm3C-seq color by global mCG **(g)** and global mCH **(h)** fraction. **i,** The average global mCG and mCH fractions for neurons in different dissection regions. Regions are ordered by the global mCH fractions, and only the top and bottom 20 regions are shown. **j,** The average global mCG and mCH fractions for all cell subclasses. Subclasses are ordered by the global mCH level, and only the top and bottom 20 subclasses are shown.

**Extended Data Figure 4.**
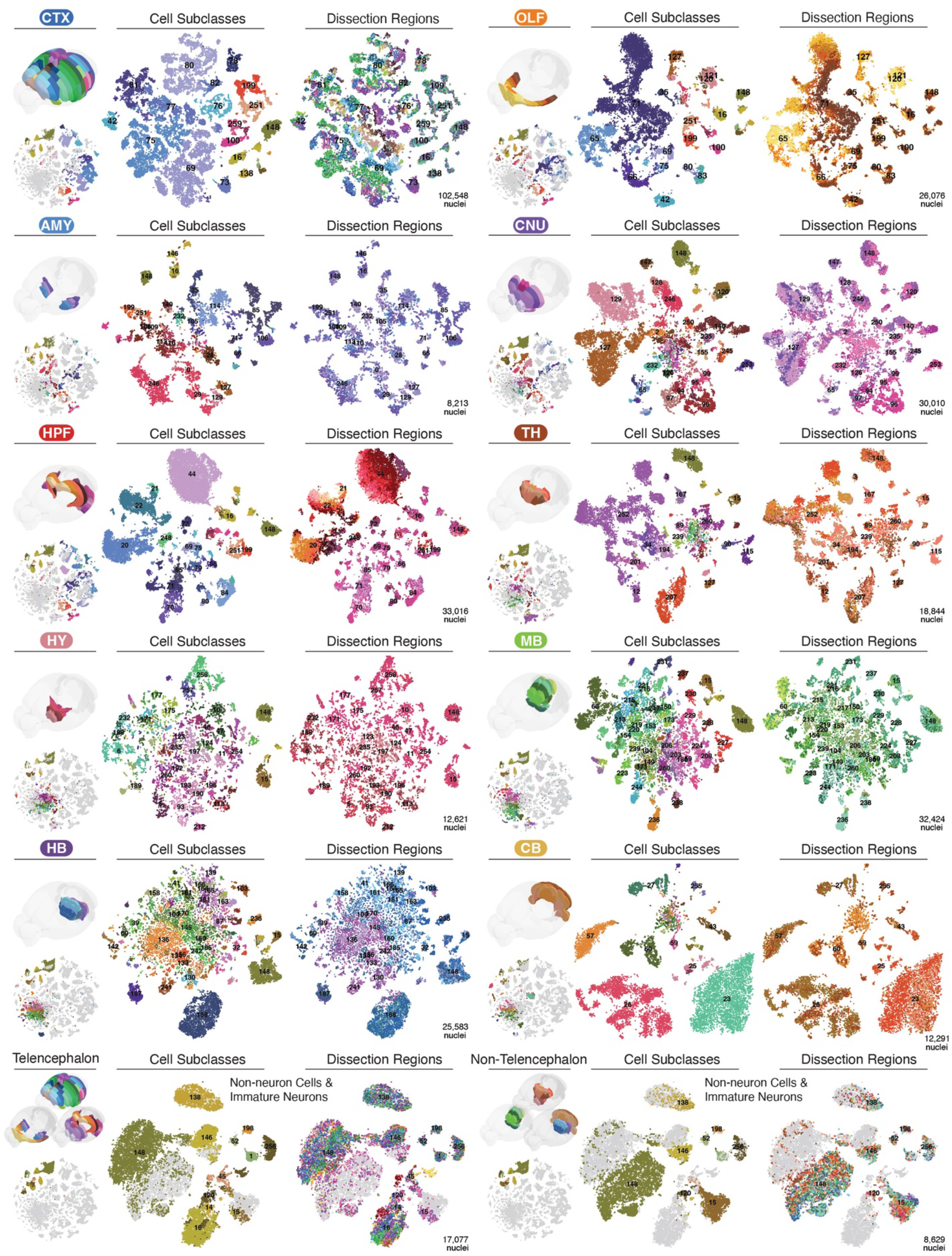
t-SNE embedding by major regions. This figure groups cells by major regions (first five rows), including isocortex (CTX), olfactory bulb (OLF), amygdala (AMY), cerebral nuclei (CNU), hippocampus (HPF), thalamus (TH), hypothalamus (HY), midbrain (MB), hindbrain (HB), and cerebellum (CB). Each section comprises three columns. The left column displays the CCF-registered 3D brain dissection regions and the corresponding cell on the whole brain t-SNE. The middle and right columns show the t-SNE embedded by cells from this major region, colored by cell subclasses and dissection regions, respectively. The numbers on the t-SNE plot indicate the cell subclass ID, which refers to in Supplementary Table 4. The final row groups non-neuron cells into two sections based on telencephalon and non-telencephalon dissection regions.

**Extended Data Figure 5.**
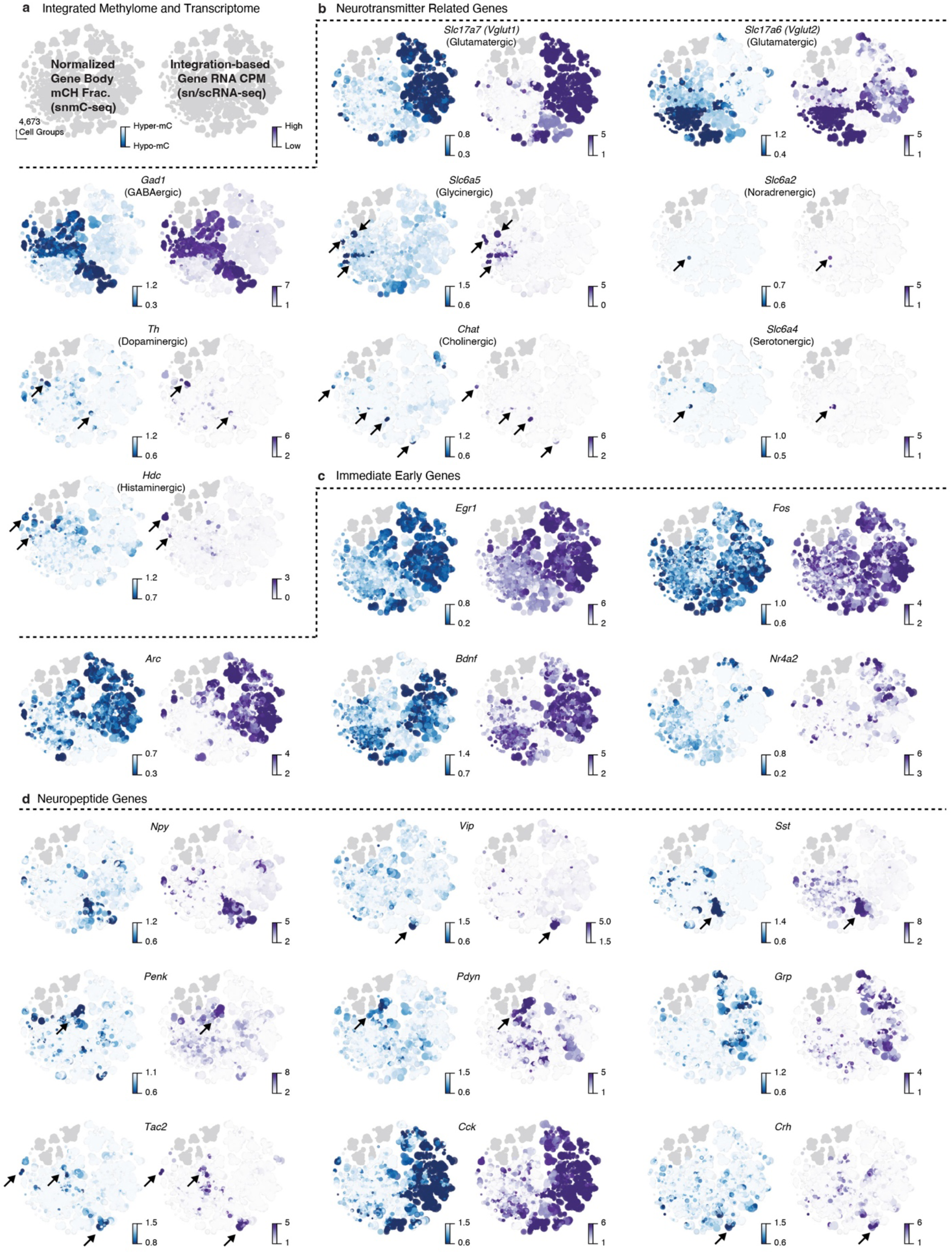
Example genes illustrating high-granularity correspondence between methylome and transcriptome. **a,** Schematic representation of the normalized gene body mCH fraction (left panel) and RNA CPM value (right panel) at the cell-group-centroids t-SNE plot for each gene. **b-d,** Example gene groups: neurotransmitter-related genes **(b)**, immediate early genes **(c),** and neuropeptide genes **(d)**.

**Extended Data Figure 6.**
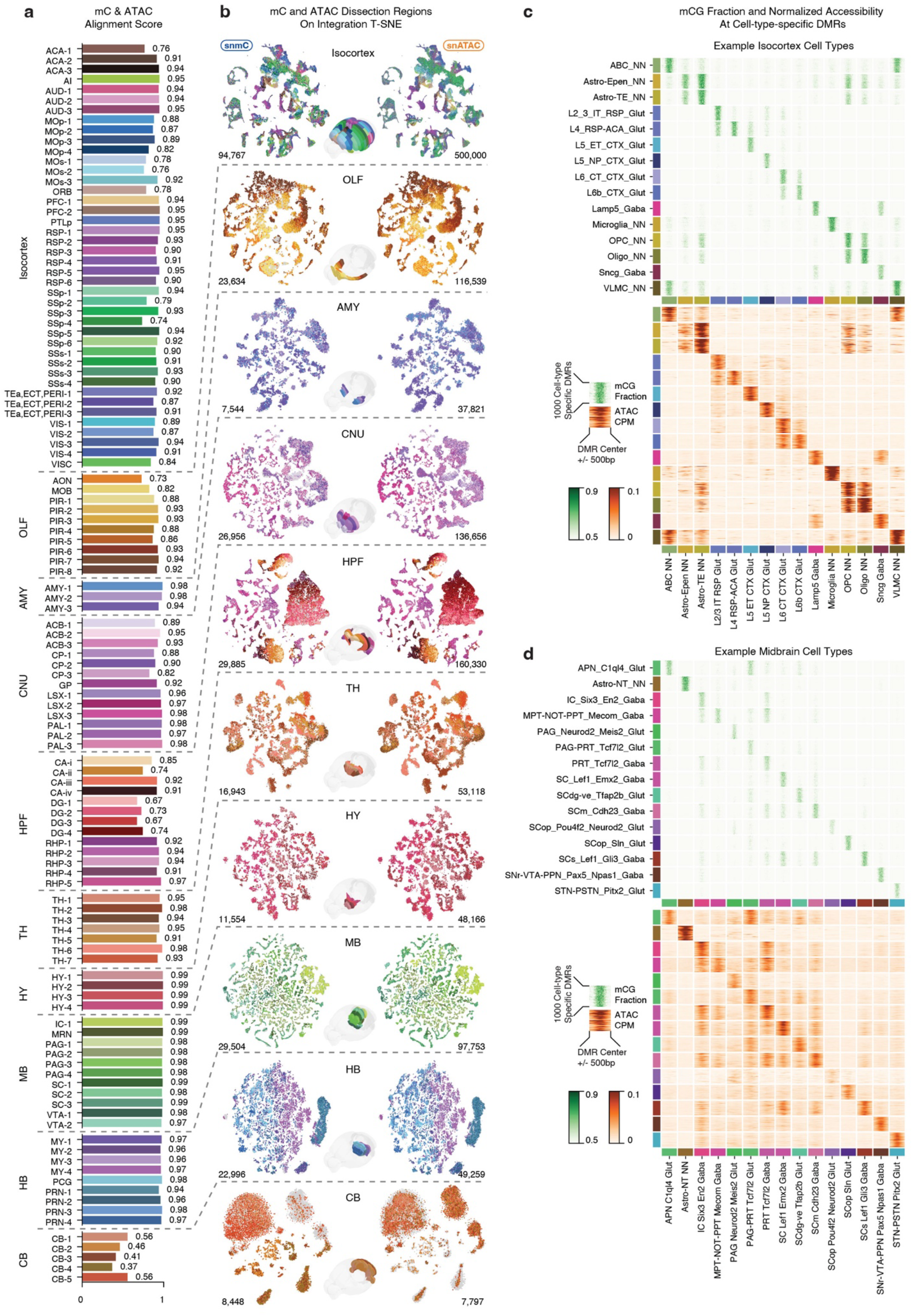
Integration of snATAC-seq and snmC-seq3 data. **a,** Barplot displays the alignment scores of each dissection region calculated the low dimensional space of snATAC-seq and snmC-seq integration. **b,** t-SNE shows the co-embedding of snmC-seq and snATAC-seq data, grouped by major regions and colored by dissection regions. **c-d,** Heatmap visualization of 15 x 15 small heatmaps. Each small heatmap represents the mCG fractions (green) and the corresponding accessibility level of 1,000 cell-type-specific CG-DMRs. Cell subclasses from isocortex **(c)** and midbrain **(d)** are shown as examples.

**Extended Data Figure 7.**
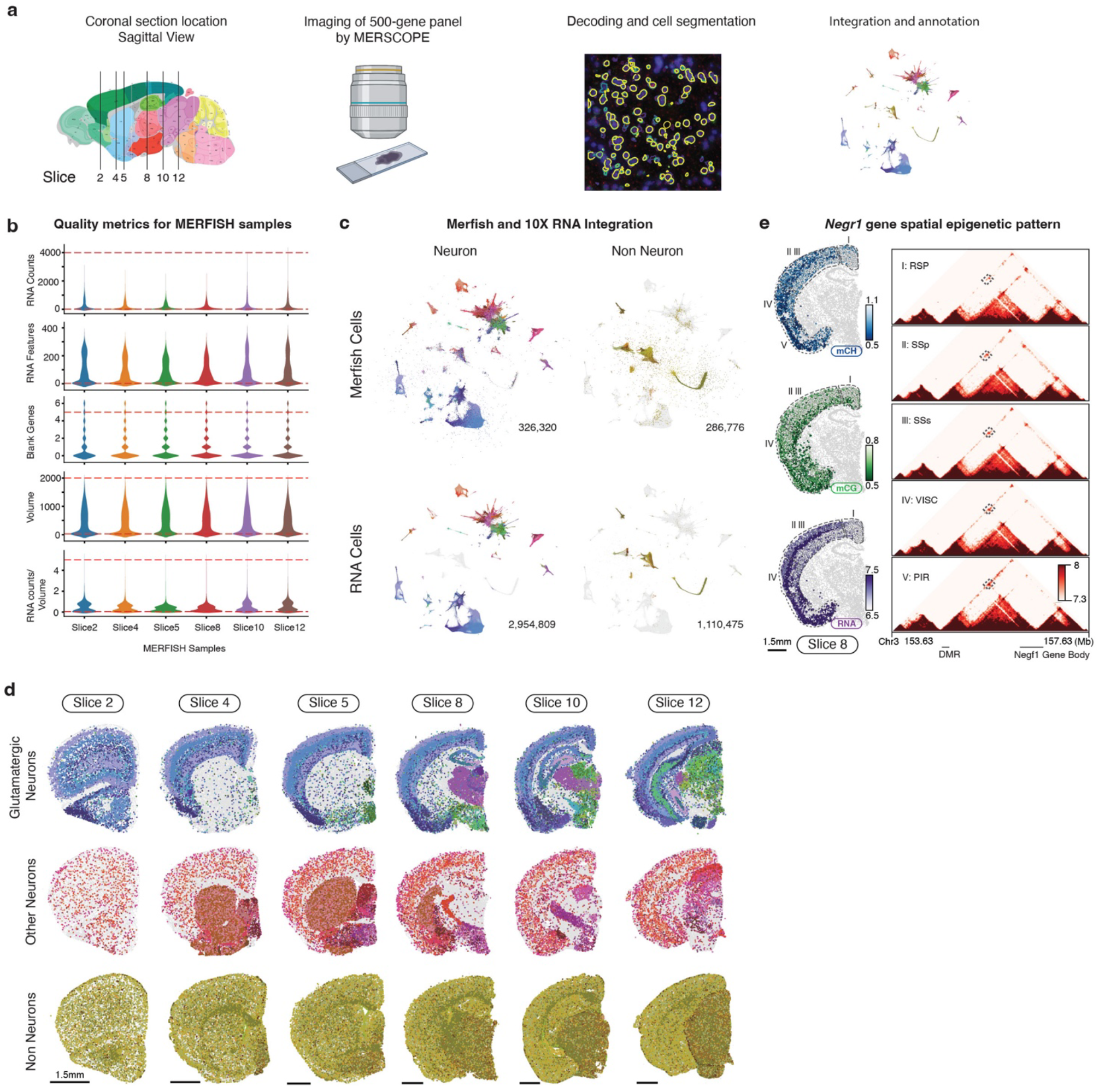
MERFISH data processing and annotation. **a,** Workflow illustrating the generation of MERFISH data, including sample preparation, imaging, and data analysis steps. **b,** Quality control assessment for each MERFISH sample, where the red lines represent the filtering cutoff for various quality metrics, including RNA total counts, RNA feature counts, blank gene number, cell volume (μm^3^), and RNA counts per volume. **c,** Integration t-SNE plot of MERFISH and scRNA dataset^7^ color by cell subclasses. **d,** MERFISH cells colored by cell subclasses, with labels obtained from the integration with the RNA dataset. From top to bottom, the cells are displayed by glutamatergic neurons, other neurons, and non-neurons. **e,** Spatial epigenetic patterns of *Negr1* and its associated DMRs. Brain slices in the left column are color-coded by normalized gene body mCH fraction, mCG fraction of the DMR (chr3:154,927,600-154,929,099), and RNA expression. The right column displays the normalized contacts heatmap between the DMR and gene.

**Extended Data Figure 8.**
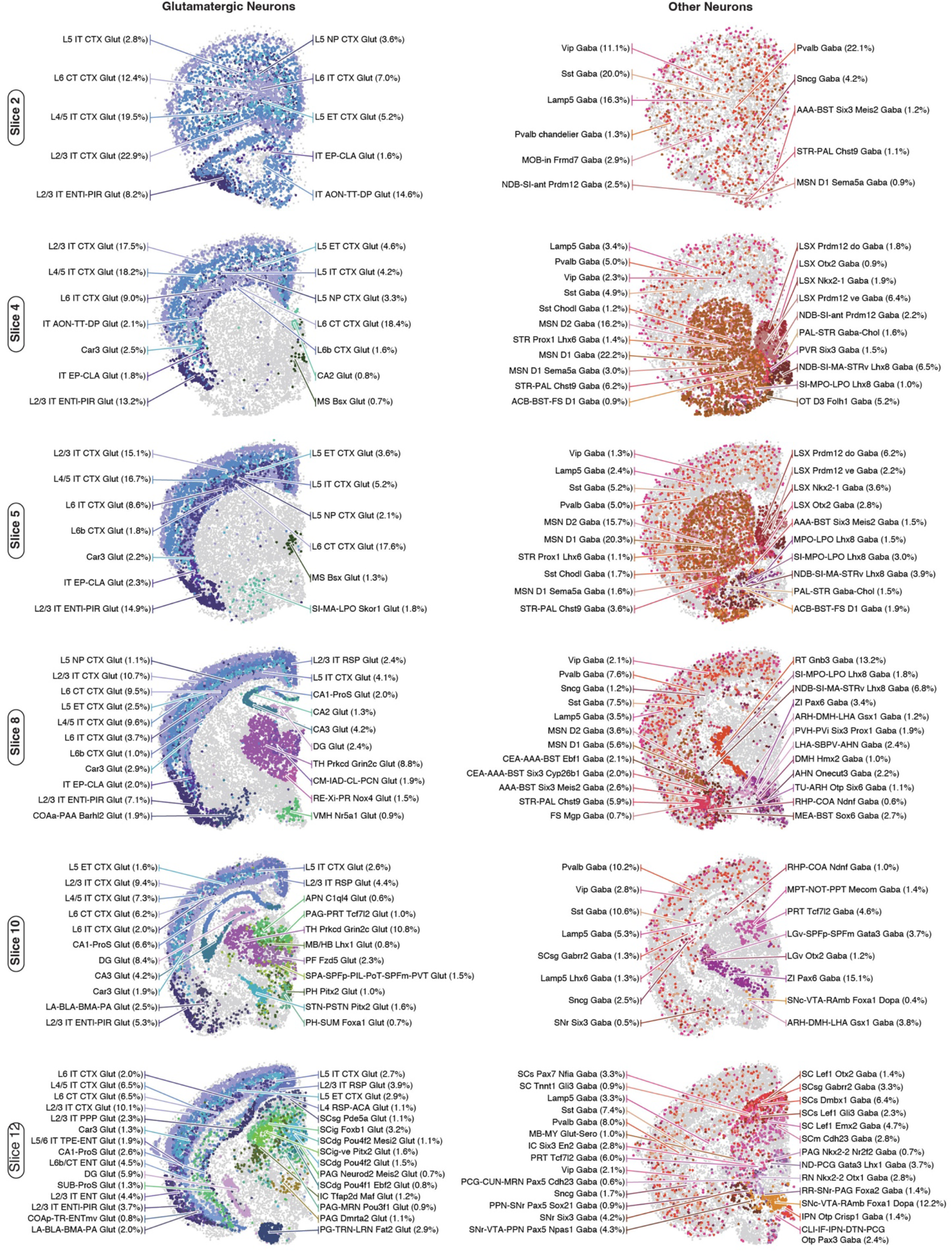
Distribution of snmC-seq cells subclasses on MERFISH slices. MERFISH plot depicting the spatial distribution of snmC-seq cells colored by cell subclass on imputed MERFISH locations (Methods). Each row represents a different MERFISH slice. The left column shows glutamatergic neurons and the right column shows other neurons. Centroids of each cell subclass are indicated by arrows, with the numbers indicating their cell proportion on that slice.

**Extended Data Figure 9.**
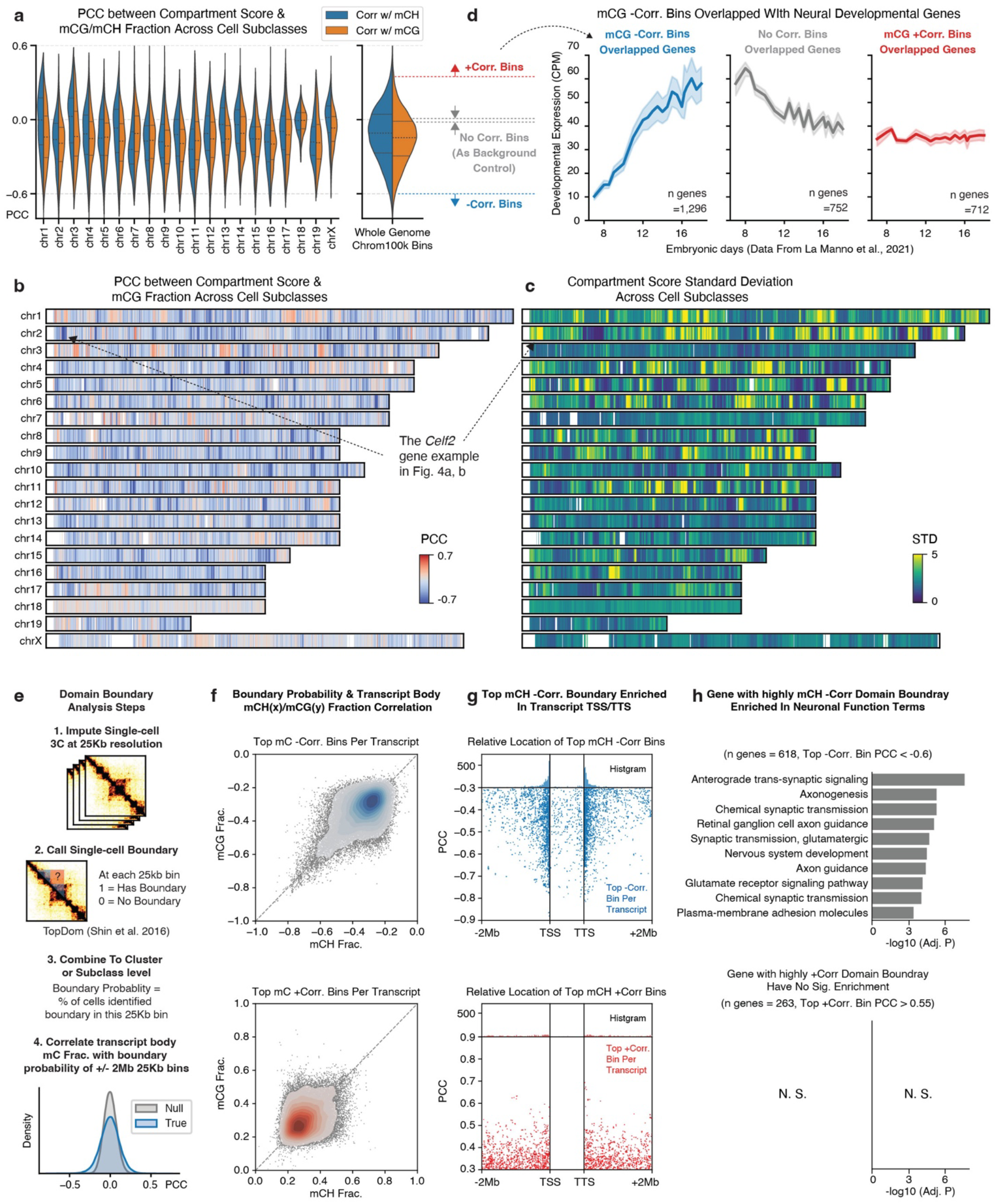
Chromatin conformation analysis at compartment and domain level. **a,** PCC between compartment score and mCG (orange)/mCH (blue) fractions of all 100kb bins on each chromosome (left panel) or whole genome (right panel). The dot lines inside each violin plot are 75%, 50%, and 25% quantiles from top to bottom. **b-c,** chromosome 1-D heatmaps show PCC between compartment score and mCG fraction **(b)** and the compartment score STD across cell subclasses **(c)** for each chromosome at a 100-Kb resolution. Arrows indicate the location of the *Celf2* gene used as an example in Fig. 4a, b. **d,** The line plot (mean±s.d.) shows the developmental gene expression level among subtypes defined in La Manno et al.^39^ across embryonic days. The genes in each subpanel are selected by overlapping with top negatively correlated (left), positively correlated (right), or uncorrelated (middle) chrom100k bins in **(a)**. **e,** Workflow for gene body domain boundary analysis. **f,** The scatter plots of the most negatively (top) or positively (bottom) correlated boundary to each long gene transcript. Both the x and y axis is the PCC between 25Kb bin boundary probability and transcript body mCH (x-axis) or mCG (y-axis) fractions. **g,** The scatterplot shows the location of each long gene transcript’s most negatively (top) or positively (bottom) correlated boundary. The y-axis is the PCC between the 25Kb bin boundary probabilities and transcript body mCH fractions; the x-axis is the relative genome location to the transcripts. **h,** Functional enrichment for genes associated with negatively correlated domain boundaries (upper) or positively correlated boundaries (lower).

**Extended Data Figure 10.**
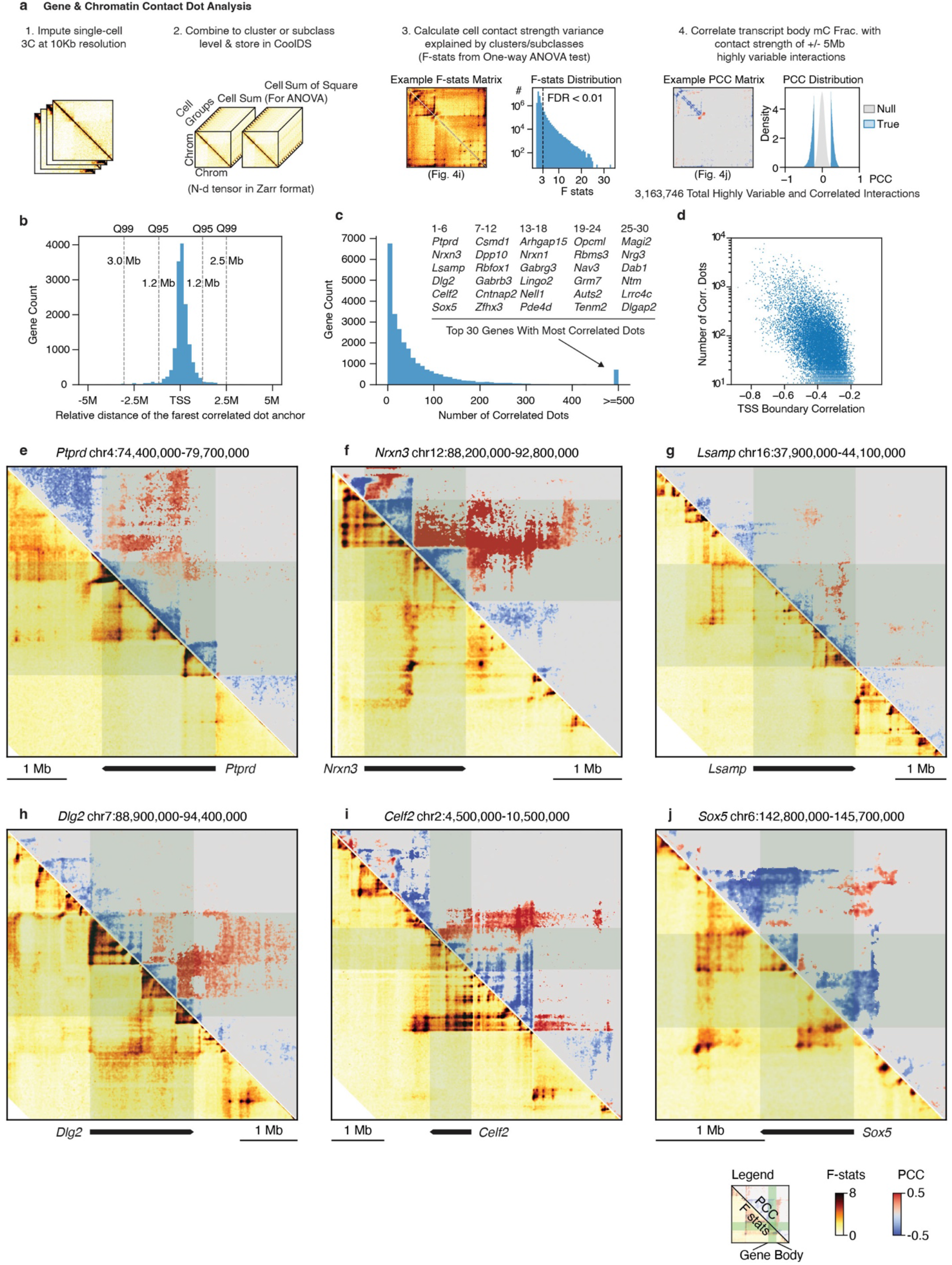
Correlation between gene expression and chromatin contacts. **a,** Workflow for highly variable and gene correlated interaction analysis. **b,** The distribution of the distance between the furthest correlated interaction and gene TSS. Q95 and Q99 stand for the quantile of all interactions ordered by the distance to TSS. **c,** Distribution of the number of highly variable and correlated interactions per gene; top 30 gene names are listed. **d,** Scatterplot shows each gene’s number of correlated interactions (y-axis) and TSS boundary probability correlation (x-axis, PCC between mCH and TSS boundary probability, from Extended Data Fig. 9e). **e-j,** Compound heatmaps display the chromatin conformation landscape of megabase-long genes, including *Ptprd* **(e)**, *Nrxn3* **(f)**, *Lsamp* **(g)**, *Dlg2* **(h)**, *Celf2* **(i),** and *Sox5* **(j)**. For each panel, green rectangles indicate the location of the gene body, the lower triangle shows the F statistics from ANOVA analysis analyzing the variance of contact strength across all cell subclasses (similar to Fig. 4i), and the upper triangle shows the PCC between contact strength and mCH fraction (similar to Fig. 4j).

**Extended Data Figure 11.**
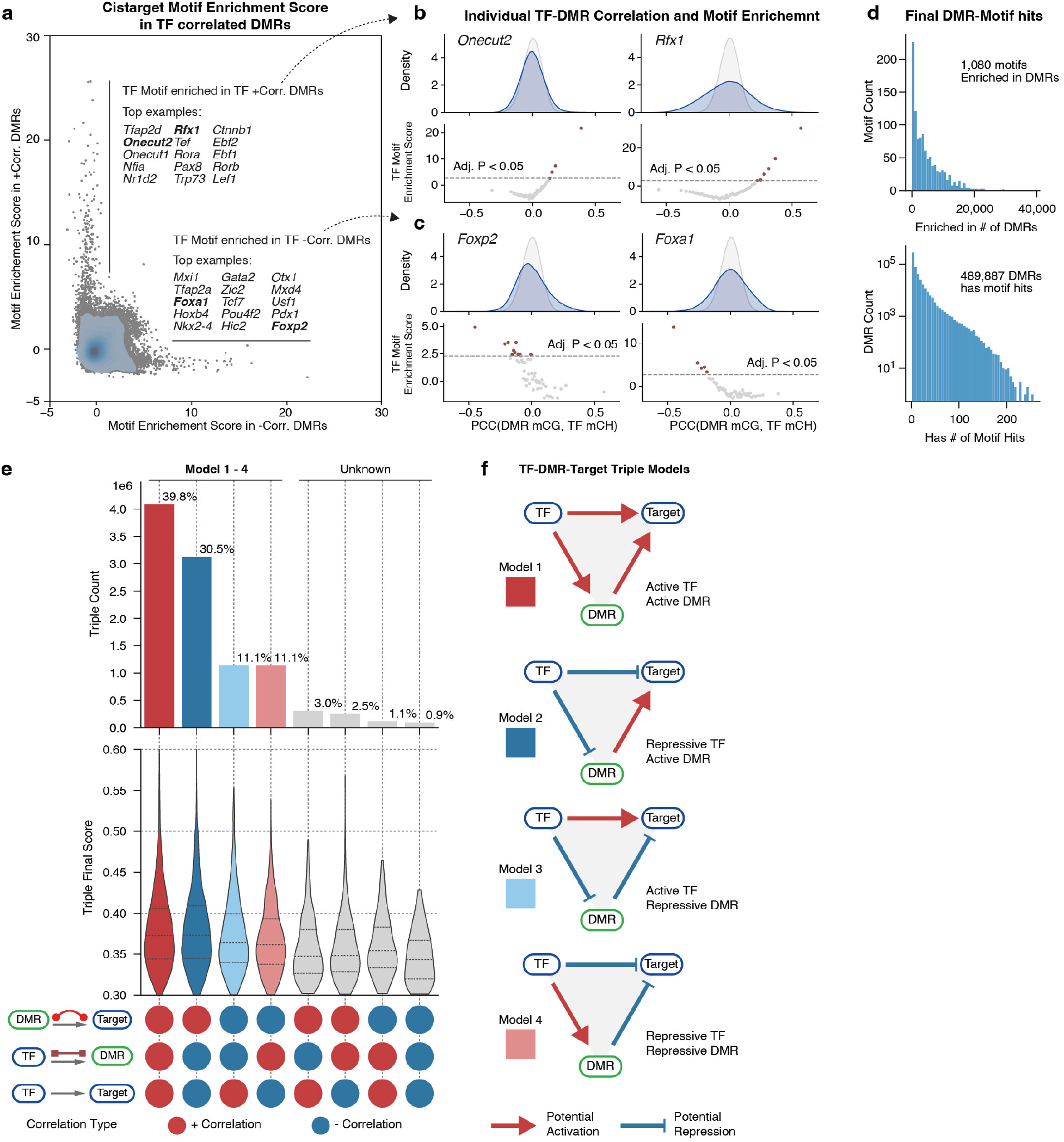
Construction of TF-DMRs-Target regulatory networks. **a,** Scatterplot shows the motif enrichment scores in negatively correlated DMRs (x-axis) and positively correlated DMRs (y-axis) for each TF. The top TFs with the highest motif enrichment scores are listed. Blue contours are the kernel density of the dots. **b-c,** Example TFs with motifs enriched in positively correlated DMRs or negatively correlated DMRs are shown in more detail (similar to Fig. 5f). The *Onecut2* and *Rfx1* gene **(b)** are examples of having motifs enriched in positively correlated DMRs, the Foxp2 and Foxa1 gene **(c)** are examples of having motif enriched in negatively correlated DMRs. **d,** The top histogram shows the distribution of the number of DMRs each motif is enriched in. The bottom histogram shows the distribution of the number of motif occurrences each DMR has. **e,** The TF-DMR-Target triples are separated into eight categories (columns) based on their PCC sign between Gene-DMR, TF-DMR, and TF-Gene. The top barplot is the triple distribution in each category. The middle violin plot is the triple final score distribution within each category. Lines inside the violin plot are 25%, 50%, and 75% quantiles, respectively. The bottom dots show the correlation sign combination of each category. Column colors match the schematic in **(f)**. **f,** The schematic displays the potential regulatory model for the four most common (based on **e**) TF-DMR-Target triple categories.

**Extended Data Figure 12.**
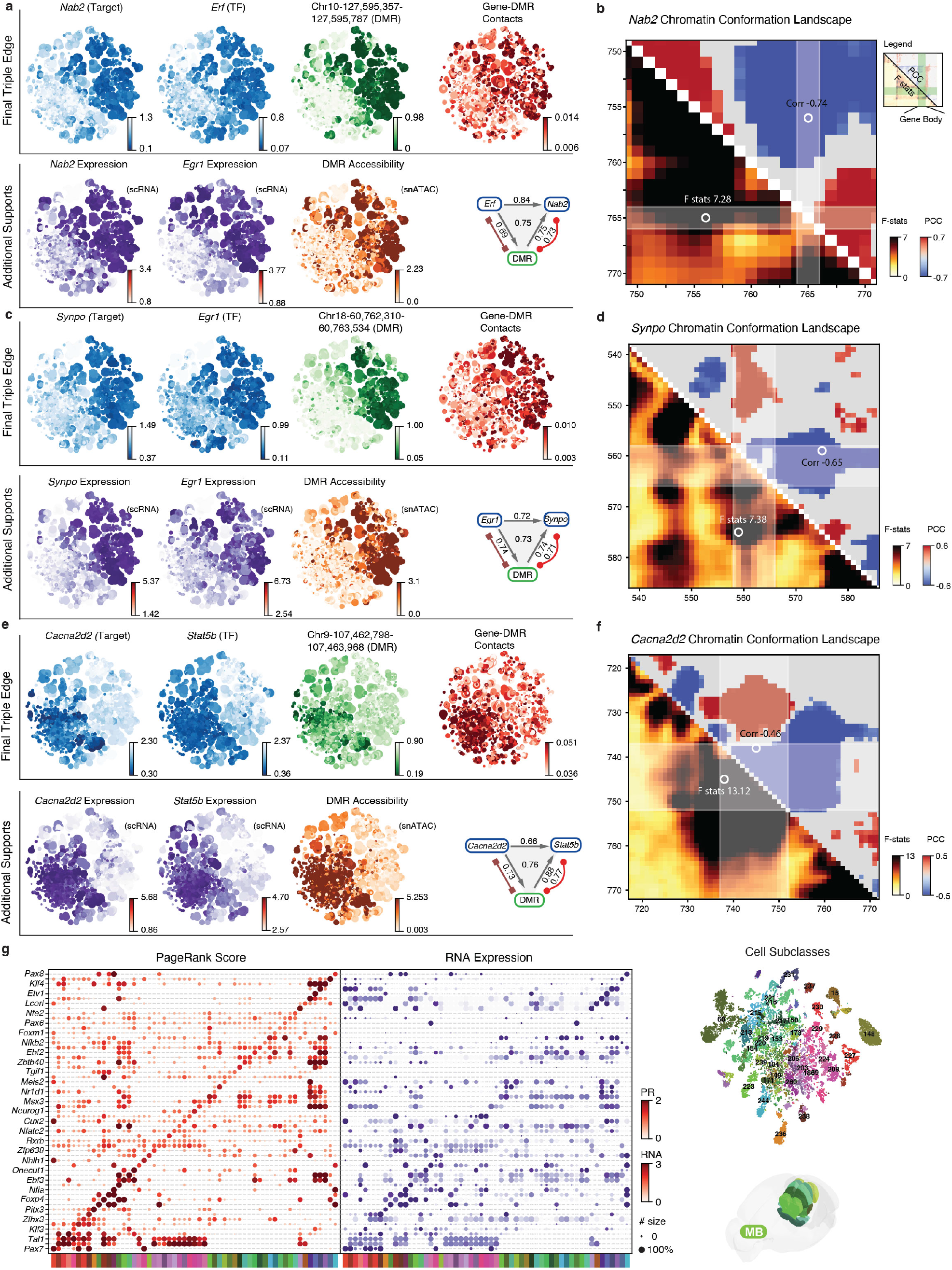
TF-DMR-Gene triple predict TF and gene relationships. **a-f**, Example TF-DMR-Target triple, including 1: Erf (TF), Nab2 (target) and DMR (Chr10:127,595,357-127,595,787) **(a-b)**; 2: Egr1 (TF), Synpo (target) and DMR (Chr18:60,762,310-60,763,534) **(c-d)**; 3: Cacna2d2 (TF), Stat5b (target) and DMR (Chr9:107,462,798-107,463,968) **(e-f)**; For each example, left are t-SNE plot colored by the mCH fraction (blue) or RNA level (purple) for target and TF; mCG fraction (green) and chromatin accessibility (orange) for DMR; and gene-DMR contact score (red) **(a,c,e)**. The compound heatmaps on the right show the chromatin landscape of target genes, including *Nab2* **(b)**, *Synpo* **(d),** and *Cacna2d2* **(f);** the layout is similar to Exnteded Data Fig. 10e-j. **g,** The dot plots represent TF’s normalized PageRank Score and RNA expression for cell subclasses in the hindbrain (MB). Red dots are colored and sized by PageRank Score. Purple dots are colored by RNA CPM, sized by the percentage of cells in that subclass expressing this gene. Right, the t-SNE plot of snmC-seq cells from MB colored by dissection region and the CCF-registered 3D brain dissection regions.

**Extended Data Figure 13.**
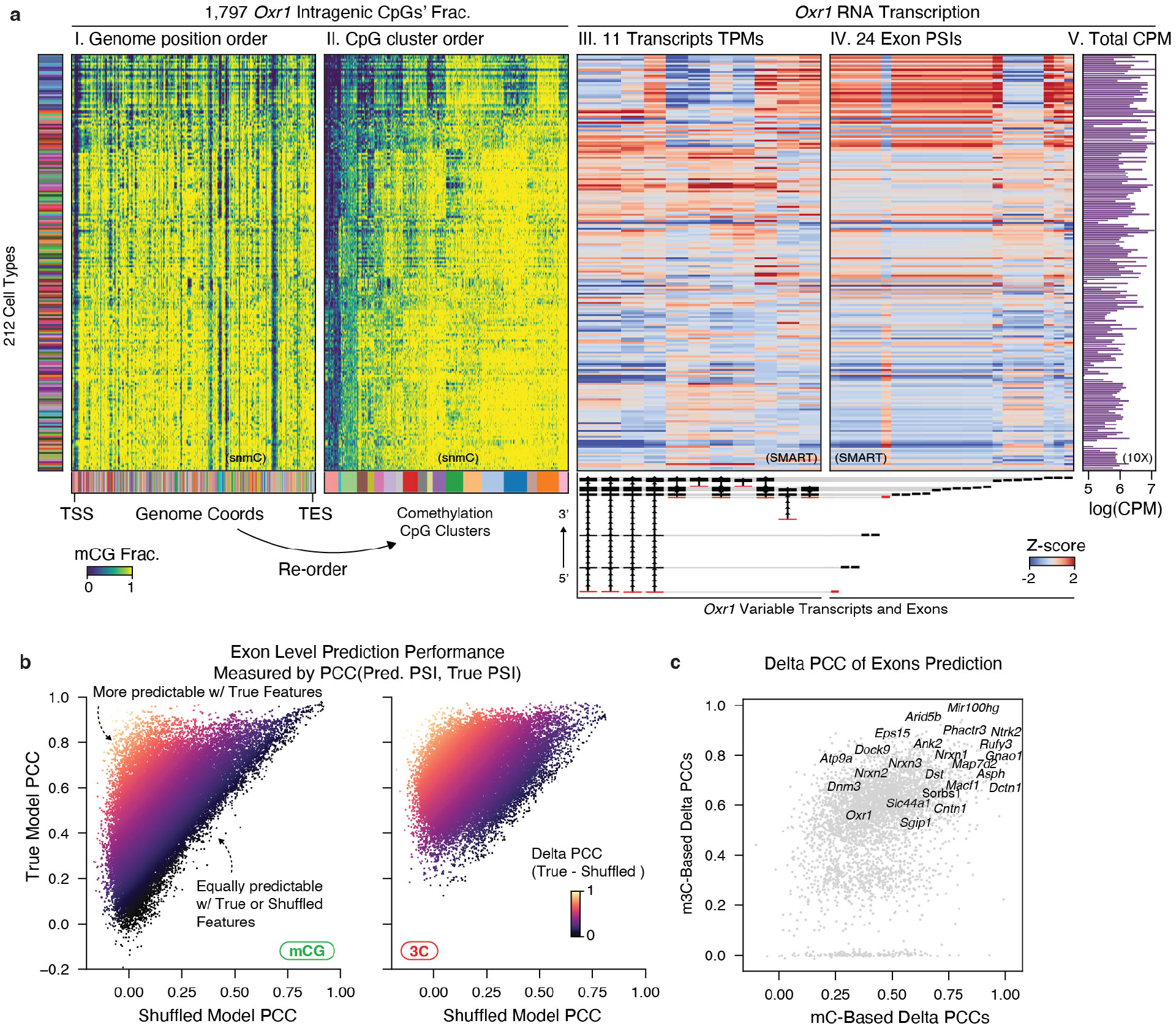
Epigenetic heterogeneity and gene exon usage. **a,** Compound heatmaps illustrate the similarity between the *Oxr1* intragenic methylation heterogeneity and alternative isoform expression patterns. Rows are neuron cell subclasses. I, mCG fraction of all 1,797 CpG sites of *Oxr1* gene with columns ordered by original genome coordinates (bottom colors are CpG clusters from heatmap ll). ll, mCG fraction of CpG sites re-ordered by their CpG clusters (bottom colors) based on subclasses methylation pattern. Heatmap lll and Heatmap lV show the TPM of 11 highly variable transcripts and PSI of 24 highly variable exons of *Oxr1*, quantified with the SMART-seq dataset. All values are z-score normalized across cell subclasses. The *Oxr1* transcript structures and exon locations are indicated at the bottom plots. Heatmap V shows the *Oxr1* gene log(CPM) in scRNA-seq (10X) data. **b,** Scatterplot shows the PCC between predicted PSI and true PSI for each highly-variable exon (dot), using methylation features (left) and chromatin contact interactions (right) to predict. **c,** Scatterplot shows the delta PCC in mC models (x-axis) and m3C models (y-axis) for highly-variable exons (dot). Top exons with large delta PCC are listed by their corresponding gene names.

## Supplementary Information Guide

### Supplementary Tables

Supplementary Table 1. snmC/snm3C sample brain dissection region metadata.

Supplementary Table 2. snmC-seq dataset cell metadata.

Supplementary Table 3. snm3C-seq dataset cell metadata.

Supplementary Table 4. snmC and snm3C cluster annotation and cross-modality cluster mapping.

Supplementary Table 5. Cell subclass brain region distribution.

Supplementary Table 6. List of 500 genes included in the MERFISH gene panel.

Supplementary Table 7. MERFISH dataset cell metadata.

Supplementary Table 8. snmC and snm3C spatial embedding based on integration with MERFISH data.

Supplementary Table 9. Glossary and abbreviations.

### Supplementary Information

Supplementary Information 1. snmC-seq3/snm3C-seq Library Preparation Protocol.

Supplementary Information 2. FANS gating examples for snmC and snm3C samples.

