## Supplementary Information 1 for "Single-cell DNA Methylome and 3D Multi-omic Atlas of the Adult Mouse Brain"

**snm3C/snmC-seq3 Beckman i7 Protocol – Ecker Lab**

**MATERIALS-**

*General reagents*

- FACS sorted single-nuclei into 384-well PCR plate
- HyClone™ HyPure™ Molecular Biology Grade (MB) Water (GE Life Sci. SH30538.08)
- 200-Proof (100%) Ethanol (Koptec V1001)

*Collection of single nuclei by Fluorescence-activated Cell Sorting (FACS)*

- M-Digestion Buffer (Zymo D5021-9)
- Proteinase K w/Storage Buffer (Zymo D3001-2-20)
- CpG Methyltransferase (M.SssI) (NEB M0226)
- Unmethylated Lambda DNA (Promega D1521)

*Bisulfite Conversion and Cleanup*

- CT Conversion Reagent (Zymo D5003-1)
- M-Solubilization Buffer (Zymo D5021-7)
- M-Dilution Buffer-Gold (Zymo D5006-2)
- M-Reaction Buffer (Zymo D5021-8)
- MagBead J (Zymo custom product D4100-3-100)
- M-Binding Buffer (Zymo D5021-7)
- M-Wash Buffer (Zymo D5040-4)
- M-Desulphonation Buffer (Zymo D5040-5)
- M-Elution Buffer (Zymo D5007-6)

*Random primers*

- Random Indexing Primer (IDT custom DNA oligo, standard desalted)

*Sera-Mag Solid Phase Reversible Immobilization (SPRI) beads*

- Sera-Mag SpeedBeads Magnetic Carboxylate Modified (GE Healthcare 5152105050250)
- Poly(ethylene glycol) PEG 8000 (Sigma cat no. 89510-250G-F)
- TE buffer pH=8.0 (Ambion AM9858)
- 5M NaCl
- 1M Tris-HCl pH=8.0
- 0.5M EDTA pH=8.0
- AMPure XP beads (Beckman Coulter cat no. A63881)
- 100 bp DNA ladder (New England Biolabs cat no. N3231L)

*snmC-seq3 Library Preparation*

- Blue Buffer (10x) (Enzymatics P7010-HC-L)
- Klenow Exo- (50U/µL) (Enzymatics P7010-HC-L)
- Deoxynucleotide Solution Mix (10mM each dNTP) (NEB N0447L)
- Exonuclease I (20U/µL) (Enzymatics X8010L)
- Shrimp Alkaline Phosphatase (rSAP) (1U/µL) (NEB M0371L)
- Sera-Mag SPRI beads

- M-Elution Buffer (Zymo 5007-6)
- EB buffer (Qiagen 19086)
- Accel-NGS® Adaptase™ Module (Swift Bio 330384)
- P5 Indexing Primer (IDT custom DNA oligo, standard desalted)
- P7 Indexing Primer (IDT custom DNA oligo, standard desalted)
- KAPA HiFi HS RM (Kapa KK2602)
- Qubit dsDNA BR Assay Kit (Thermo Fisher Q32850)

*Equipment*

- 384-Well Hardshell PCR Plate Clear, 20 Pcs/pk (Thermo Fisher 4483285)
- 96-Well Hardshell PCR Plate GPLE, 20 Pcs/pk (Thermo Fisher 4483348)
- MicroAmp™ Splash-Free 96-Well Base (Thermo Fisher 4312063)
- Plateone® Deep 384-Well 200 µL Polypropylene Plate (USA-SCI. 1884-2410)
- Reservoir Single Well 384 Bottom Low Profile, Clear (Axygen RES-SW384-LP)
- Reservoir Single Well 96 Bottom High Profile, Clear (Axygen RES-SW96-HP)
- Reservoir Single Well 96 Bottom Low Profile, Clear (Axygen RES-SW96-LP)
- 15mL Centrifuge Tubes (Olympus 28-103)
- 50mL Centrifuge Tubes (Olympus 28-106)
- 1.7mL Microtube, Clear (Olympus 24-282LR)
- Microamp Clear Adhesive Film, 100pc (Thermo Fisher 4306311)
- Speedball Deluxe Soft Rubber Brayer, 4 inches (Statesville N.C.)
- 37°C Incubator
- 384-well and 96-well Compatible Thermocycler
- DynaMag™-96 Side Magnet (Thermo Fisher 12331D)
- DynaMag™-2 Magnet (Thermo Fisher 12321D)
- 384-well Magnet Plate (Alpaqua A001222)
- Allegra 25R Centrifuge with S5700 2x96 Swinging Bucket Rotor (Beckman Coulter 368954)
- Axygen® 384-well tips, 45µL, Clear, Filtered, Sterile, Fine Point, SLAS Rack (Corning FXF-384-45FP-R-S)
- Axygen® 96-well tips, 200µL, Clear, Filtered, Sterile, SLAS Rack (Corning FXF-200-R-S)
- Axygen® 96-well tips, 30µL, Clear, Filtered, Sterile, SLAS Rack (Corning FXF-50-R-S)
- Beckman Coulter Biomek i7 dual-Multichannel with MC384 and MC96 Pipetting Heads

**REAGENT SETUP-**

*Methylated λ DNA*

Dilute supplied SAM to 1600µM with Nuclease-free Water. Set up the reaction in the following order:

| Reagent | Volume |
|---|---|
| Nuclease-free Water | up to 50 µL (36 µL) |
| Methyltransferase Reaction Buffer (10x) | 5 µL |
| Diluted SAM (1600 µM) | 5 µL |
| DNA | 1 µg (2 µL) |
| Methyltransferase | 8 units (2 µL) |

Mix by pipetting. Incubate at 37°C for 1 hr. Stop the reaction by heating at 65°C for 20 minutes. Purify product by Sera Mag bead cleanup (1.6 bead vol : sample vol ratio) or Qiagen DNA Column Cleanup.

*Digestion Buffer Plates (for FACS sorting)*

In a 50mL tube, combine 15mL M-Digestion Buffer (incubate for 10 minutes at 37°C to dissolve precipitate) with 14mL MB water. Resuspend Proteinase K with 1mL Proteinase K Resuspension Buffer and add 1mL to the digestion buffer mix. Add 1.5µL methylated λ DNA to aid determination of bisulfite conversion efficiency. Invert to mix. Aliquot 1µL per well to 50 384-well plates to use for FACS sorting of stained nuclei (This is automated using Biomek i7). Plates can be stored for 3 months at -20°C.

*Sera-Mag Solid Phase Reversible Immobilization (SPRI) Beads*

1. Mix Sera-Mag SpeedBeads and transfer 1mL to a 1.5ml tube.
2. Place SpeedBeads on a magnetic stand until clears and carefully remove the supernatant. Wash the beads twice with 1ml of TE. For each wash, remove the tube from the magnet and mix by inversions. Resuspend the washed beads in 1ml of TE.
3. Add 9g of PEG 8000 to a new 50ml sterile conical tube.
4. Add 10ml of 5M NaCl to the 50ml tube.
5. Add 500µl of 1M Tris-HCl pH=8.0 and 100ul of 0.5M EDTA pH=8.0 to the 50ml tube.
6. Mix until all dissolves into solution.
7. Add 1ml of resuspended SpeedBeads to the 50ml tube and fill the volume with MB water.
8. Test against AMPure XP beads using 100bp DNA ladder.

*Ethanol, 80%*

To 40mL of 200-proof Ethanol, add 10 µL MB water in a 50mL tube. Keep solution sealed when not in use. Prepare ~9 tubes fresh before each 16-plate library preparation.

*Random Primer solution*

Sequences of Random Primers with unique barcodes are provided in the end. Transfer 1.5 µL stock primer (100 µM) into 298.5 µL MB water. Vortex to mix. If preparing for 384-well plates, aliquot into deep-well 384-well plates.

*PCR Primer Mix*

Sequences of indexing primers with unique dual barcodes are provided in the end. Each PCR Primer Mix contains a P5L indexing primer (600 nM) and a P7L indexing primer (1 µM).

**PROTOCOL-**

**SECTION I: Prepare Single Nuclei**

A. Prepare Stock Solutions:
    1. NIM: Sucrose (250 mM), KCl (25 mM), MgCl2 (5 mM), Tris-Cl pH 8.0 (10 mM)
    2. Dilutent: Tris-Cl pH 8.0 (120 mM), KCl (150 mM), MgCl2 (30 mM)
B. Prepare the following solutions fresh on ice before each experiment:
    1. NIMT: NIM + Triton X-100 (0.1%) + DTT (1 mM) + Proteinase Inhibitor (1:100 dilution)
    2. 50% Iodixanol: 5 vol. Optiprep (60% Iodixanol) + 1 vol Dilutent
    3. 25% Iodixanol: 1 vol. 50% Iodixanol + 1 vol NIM
    4. DPBS + 1% BSA
C. Prepare Nuclei
    1. Fast cool centrifuge with swinging bucket tube rotor to 4°C.
    2. Remove frozen tissue from -80°C, place on ice.
    3. Transfer tissue with 2.4 mL NIMT from tube into the large dounce.
    4. Use loose pestle A 40 times gently without introducing bubbles.
    5. Use tight pestle B 40 times gently without introducing bubbles.
    6. Transfer lysed solution into a 5-mL or 15-mL tube and keep on ice.
    7. If doing NeuN stain: Add 6 µL of NeuN 488 (1:500 dil). Mix and incubate 15 min on ice, keeping solution covered from light.
    8. Mix the lysis suspension with 1.5 mL of 50% Iodixanol by pipetting.
    9. Slowly pipette 1 mL of cell mixture onto 500 µL 25% Iodixanol cushion, 4 x 2-mL tubes in total per sample.
    10. Centrifuge in swing rotor for 20 min at 10,000 x g at 4°C.
D. Nuclei Staining and Count
    1. Remove supernatant very carefully, avoiding debris.
    2. Using ice-cold DPBS+RNase Inhibitors, resuspend and combine the pellets from each tube into a total of 1 mL nuclei solution in a new tube.
    3. Add Hoechst 33342 (dil: 1:1000): Dilute the Hoechst 1:10 (5 µL + 45 µL DPBS), then add 5 µL to 1 mL sample.
    4. Incubate on ice for 5 min.
    5. Mix 10 µL Nuclei suspension with 10 µL Trypan Blue and mix by pipetting.
    6. Add 10 µL stained solution to a Cell Counter Slide and read using the Bio-Rad Cell Counter TC-20.
        a. Nuclei count is Total Count minus Live Count

**SECTION II: 3C Protocol**

E. Crosslinking
    1. Split sample into 2 tubes (1-2 million nuclei in each tube) and bring volume up to 1 mL with DPBS.
    2. Add 57 µL of 37% formaldehyde (for a final concentration of 2%) to each tube and mix well by inverting 10 times.
    3. Incubate 10 min (5 min if a NeuN pre-stain was done to avoid stain degradation) at room temperature, inverting occasionally.

4. Add 91.9 µL Stop Solution (or 2.5 M Glycine) and mix well by inverting 10 times.
5. Incubate for 5 min at room temperature, inverting occasionally.
6. Place sample on ice and incubate an additional 5 min.
7. Pellet cells by centrifuging 5 min at 2500 x g or 10 min at 1000 x g.
8. Discard supernatant, being sure to leave no residual liquid, and resuspend in 1 mL of 1x PBS.
9. Repeat steps 7-8 two more times to completely wash away residual formaldehyde/glycine.
10. Proceed directly to next section or freeze samples at -80°C up to 5 days.

F. 3C Nuclei Conditioning
1. Resuspend each reaction of crosslinked nuclei in 20 µL DPBS.
2. Add 24 µL Conditioning Solution and mix gently by pipetting.
3. Incubate at 62°C for 10 min. If using a thermal cycler, set the lid temperature to 85°C.
4. Add 20 µL Stop Solution 2 and mix gently by pipetting.
5. Incubate at 37°C for 15 min. If using a thermal cycler, set the lid temperature to 85°C.

G. 3C Enzymatic Reactions

Note: Some of the next steps require addition of several reagents in the same step. These reagents should be combined into master mixes using the following tables.

1. Digestion: Add 28 µL of master mix containing the following reagents:

| | Volume per reaction | 10% extra | | # Reactions | | Final |
|---|---|---|---|---|---|---|
| **1X CutSmart** | 12 µL | 13.2 µL | **x** | | **=** | |
| **10X CutSmart (or Buffer H)** | 7 µL | 7.7 µL | **x** | | **=** | |
| **NLAIII /Enzyme H1** | 4.5 µL | 4.95 µL | **x** | | **=** | |
| **MboI /Enzyme H2** | 4.5 µL | 4.95 µL | **x** | | **=** | |
| **Total** | **28 µL** | | | | **=** | |

2. Mix gently by pipetting and incubate as follows. If using a thermal cycler, set the lid temperature to 85°C.
   a. 37°C for 60 min (or up to overnight)
   b. 65°C for 20 min
   c. 25°C for 5 min
3. Mix gently by inversion.
4. Ligation: Add 82 µL of master mix containing the following reagents:

| Reagent | Volume per reaction | 10% extra | | # Reactions | | Final |
|---|---|---|---|---|---|---|
| **Buffer C** | 70µL | 77µL | x | | = | |
| **Enzyme C** | 12µL | 13.2µL | x | | = | |
| **Total** | **82µL** | | | | = | |

5. Mix gently by pipetting and incubate at room temperature for 15 min.
6. Samples can be stored at 4°C overnight or continue to FACS.

**SECTION III: FACS**

H. Prepare samples for FACS

1. Combine the reactions for each sample and add DPBS + 1% BSA up to a total volume of 1 mL.
2. Centrifuge at 1,000 x g for 10 min at 4°C in swinging bucket rotor.
3. Remove supernatant and gently resuspend in 200 µL DPBS + 1% BSA. Add another 800 µL DPBS + 1% BSA to bring total volume up to 1 mL.
4. Add 5 µL 1:10 diluted Hoecsht 33342.
5. Add 1 µL of antibody to enhance labeling.
6. Incubate on ice for 5 min.
7. Filter the cells through a 40 µm cell strainer and transfer into a polypropylene tube for sorting. Keep on ice until ready to sort.

I. Single nuclei sorting
1. Thaw and spin down (5 sec at 1000 x g) 384-well plates containing 1 µL digestion buffer.
2. Sort single nuclei into wells using BD Influx or other sorter.
    a. Use 2N gating for Hoechst stain
    b. If NeuN 488 stained: Sort NeuN+ in col 1-22 and NeuN- in col 23-24
    c. Sort using 1 drop single mode
3. After sorting spin down the sample plates for 5 sec at 1000 x g.
4. Incubate plates for 20 min at 50°C to run proteinase K reaction.
5. Plates can be stored at -20°C for up to 1 year.

**SECTION IV: Bisulfite Conversion**

J. Prepare the Zymo CT Conversion Reagent (3 bottles for 16 x 384-well plates.
1. Add 7.9 mL Solubilization Buffer and 3 mL Dilution Buffer to each Conversion Reagent bottle and mix for 10 minutes.
2. Add 1.6 mL Reaction Buffer to each Conversion Reagent bottle and mix for an additional 5 minutes.

K. Bisulfite Reaction

*This step is automated using Biomek i7
1. Add 6 µL CT Conversion Reagent to each well.
2. Seal, vortex, and quick spin (500 rcf) plates.
3. Incubate plates using [DIRECT] method on PCR machines with 384-well plate block.
    a. 98 °C for 8 minutes
    b. 64 °C for 3.5 hours
    c. Hold at 4 °C for up to 20 hours

L. Bisulfite Cleanup

*Steps in this section are automated for up to 8 x 384-well plates using Biomek i7
1. Prepare MagBead master plate:
    a. Invert Magbead J stock tube repeatedly to ensure homogeneity.
    b. Using an 8-channel pipette, aliquot 58 µL Magbead J stock to each well of a 96-well plate (avoid bubbles and do not spin). Tap plate to remove bubbles.
    c. Distribute 14 µL beads to each well of a clean 384-well plate.
    d. Tap down 384-well plate and pipette out any bubbles from the bottom of the wells (do not spin).
    e. Distribute 1.5 µL to each well of sample plates.

2. Bind the free DNA to the Magbeads.
    a. Add 20 µL M-Binding Buffer to each well and mix by pipetting.
    b. Incubate at room temperature for 5 minutes.
    c. Move the plates onto the magnets.
    d. Remove supernatant and discard in the waste trough.
3. Wash the beads.
    a. With the plates still on the magnets, add 20 µL M-Wash Buffer to all plates and incubate at room temperature for 30 seconds.
    b. Remove supernatant and discard in the waste trough.
4. Add M-Desulphonation Buffer to the plates.
    a. Add 15 µL M-Desulphonation Buffer to all plates.
    b. Incubate at room temperature for 15 minutes.
    c. Remove supernatant and discard in the waste trough.
5. Wash the beads.
    a. Add 20 µL M-Wash Buffer to all plates and incubate at room temperature for 30 seconds.
    b. Remove supernatant and discard in the waste trough.
    c. Repeat steps 7a-b an additional time.
6. (Optional) Do an extra supernatant removal to further empty out wells if desired.
7. Let the plates dry
    a. Let plates sit for 5 minutes. Keep them on the magnets to avoid beads jumping due to static.
    b. Check that all the wash buffer has evaporated before moving on.
8. Elute the DNA from the beads using the RP elution plate
    a. Add 5.5 µL appropriate Random Primer solution to all plates (note the barcodes).
    b. Remove the plates from the magnets and seal well.
    c. DO NOT VORTEX; instead, throw the sealed plates onto the bench 10-15 times, until beads appear to be mixed into elution buffer. Repeat for all plates.
    d. Quick spin plates for 5s at 200xg
    e. Incubate at room temperature for 5 minutes. Place plates back onto the magnets.
9. Transfer eluted DNA into clean 384-well plates.
    a. Transfer 5 µL sample elution into a clean, labeled 384-well plate
    b. Seal new sample plates and quick spin for 5s at 500xg.
10. If running two sets of plates, repeat steps 1-9 for next set of 8 plates.

**SECTION V: Random Priming**

Steps M-N are automated for up to 16 x 384-well plates using Biomek i7

M. Random Primed DNA Synthesis
    1. Prepare appropriate volume of Random Priming Master Mix (RP MM) in a 50-mL conical tube and invert to mix.

| Reagent | Volume per reaction | Volume for 8 plates | Volume for 16 plates |
|---|---|---|---|
| **Nuclease-free $H_2O$** | 3.45 µL | 12420 µL | 24840 µL |
| **Enzymatics Blue Buffer (10x)** | 1.025 µL | 3690 µL | 7380 µL |
| **dNTPs (10 mM each)** | 0.50 µL | 1800 µL | 3600 µL |
| **Klenow Exo- (50 U/µL)** | 0.025 µL | 90 µL | 180 µL |

2. Denature 8 sample plates at 98 °C for 3 minutes in thermocycler. Remove plates from PCR machines and immediately place on ice to prevent rehybridization. Proceed once bottom of each plate is cool to the touch.
3. Add 5 µL RP MM to each sample well. Seal, vortex, and quick spin (5s at 500xg) sample plates.
4. Incubate plates in thermocycler with the following program:
    a. 4 °C for 5 minutes
    b. 25 °C for 5 minutes
    c. 37 °C for 60 minutes
    d. Hold at 4 °C
5. Repeat steps 2-4 for second set of 8 plates (if applicable).

N. Inactivation of free primers & dNTPs
1. Prepare appropriate volume of Exo/rSAP Master Mix (E/S MM) in a 15-mL conical tube and invert to mix.

| Reagent | Volume per reaction | Volume for 8 plates | Volume for 16 plates |
|---|---|---|---|
| **Nuclease-free $H_2O$** | 1.15 µL | 4140 µL | 8280 µL |
| **Enzymatics Blue Buffer (10x)** | 0.2 µL | 720 µL | 1480 µL |
| **Exonuclease I (20 U/µL)** | 0.1 µL | 360 µL | 720 µL |
| **rSAP (1 U/µL)** | 0.05 µL | 180 µL | 360 µL |

2. Add 1.5 µL Exo/Sap Master Mix to each sample well. Seal, vortex, and quick spin sample plates (5s at 500xg).
3. Incubate plates in thermocycler with the following program:
    a. 37 °C for 30 minutes
    b. Hold at 4 °C
4. Repeat steps 2-3 for second set of plates (if applicable).
5. Either continue to reformat and cleanup or store at -20 °C until needed for next step (no longer than overnight).

*NOTE: If not continuing to the next steps within 2 hours, plates should be moved to -20°C to avoid excess Exonuclease activity damaging sample DNA.*

**SECTION VI: Reformat and Sample Cleanups**

Steps in this section are automated for 16 x 384-well plates using Biomek i7. Note that the bead cleanups for each set of plates will overlap to increase efficiency.

O. Compress 16 x 384-well plates to 8 x 96-well plates and add SeraMag beads
1. Thaw (if applicable) and quick spin (5s at 500xg) sample plates.
2. Compress 384-well plates 1-8 into four new 96-well plates 1-4. 8 wells will be combined into one well. One 384-well plate will be compressed to half a 96-well plate.
    a. Combine samples from each quadrant of the top half of 384-plate 1 (Rows A-H) and dispense into the top half of a clean 96-well plate (Rows A-D).
        i. Eg: A1,A2,B1,B2 of 384-well plate combine into A1 of the 96-well plate. A3,A4,B3,B4 of 384-well plate combine into A2 of the 96-well plate. Etc
    b. Combine samples from each quadrant of the bottom half of 384-plate 1 (Rows I-P) and dispense into top half of the same 96-well plate (Rows A-D).
        i. Eg: I1,I2,J1,J2 of 384-well plate combine into A1 of the 96-well plate along with the A/B row samples.

c. Continue this pattern of pooling with 384-well plate 2 going into the bottom half of the same 96-well plate. Plates 3-4 will go into a second 96-well plate, 5-6 into a third 96-well plate, and Plates 7-8 into a fourth 96-well plate.

3. Bind the free DNA to the Sera-Mag beads in 96-well plates 1-4.
   a. Thoroughly mix Sera-Mag beads by repeatedly inverting bottle and pour Sera-Mag beads to the brim of a low-profile 96-well reservoir.
   b. Add 73.6 µL beads into sample plates and pipette up and down to mix.
   c. Let plates incubate at room temperature for at least 5 minutes.
4. Compress 384-well plates 9-16 into four new 96-well plates 5-8.
   a. Combine 384-well plates 9-16 following the same pattern of pooling in Step 2 above. Plates 9-10 will go into a fifth 96-well plate, 11-12 into a sixth 96-well plate, Plates 13-14 into a seventh 96-well plate, and Plates 7-8 into a eighth 96-well plate.
5. Bind the free DNA to the Sera-Mag beads in 96-well plates 5-8.
   a. Thoroughly mix Sera-Mag beads by repeatedly inverting bottle and pour Sera-Mag beads to the brim of a low-profile 96-well reservoir.
   b. Place plates 1-4 onto 96-well magnet and incubate for at least 5 minutes.
   c. Add 73.6 µL beads to sample plates 5-8 and pipette up and down to mix. Incubate for at least 5 minutes.

P. Perform bead cleanups on 96-well plates 1-8
1. Carefully remove supernatant from plates 1-4 and discard in waste trough.
2. Wash beads in plates 1-4 with 80% EtOH.
   a. Keeping plates on the magnets, add 180 µL of 80% EtOH to plates 1-4 and incubate at room temperature for at least 30 seconds. Subsequently remove and discard EtOH in the waste trough.
   b. Repeat for a second wash.
   c. Remove plates from the magnets and incubate until beads are dry. The beads should appear cracked and there should be no visible droplets of EtOH in the wells.
3. Remove supernatant and add EtOH to 96 plates 5-8.
   a. Carefully remove supernatant from plates 5-8.
   b. Add 180 µL EtOH to plates 5-8 and incubate for at least 30 seconds.
4. Elute DNA from plates 1-4 and perform second EtOH wash on plates 5-8.
   a. Once beads are dry, add 10 µL EB to plates 1-4. Set a 5-minute timer upon completion of transfer.
   b. Seal plates, vortex until beads no longer remain on the sides of the well, and quick spin 5s @ 150xg to avoid pelleting beads. When 5-minute timer is completed, replace plates on the magnets.
   c. Remove EtOH from plates 5-8 and discard in the waste trough.
   d. Repeat EtOH wash of plates 5-8.
   e. Remove plates from the magnets and incubate until beads are dry.
5. Elute DNA from plates 5-8.
   a. Once plates are dry, add 10 µL EB to plates 5-8. Start a 5-minute timer upon completion of transfer.
   b. Seal plates, vortex until beads no longer remain on the sides of the well, and quick spin 5s @ 150xg to avoid pelleting beads. When 5-minute timer is completed, replace plates on the magnets.

Q. Compress 8 x 96-well plates to 1 x 96-well plate

1. Further compress plates 1-4 into a clean 96-well plate (Rows A-D). 4 wells will combine into 1 well of the new plate. Each original 384-well plate will be contained in one row of the 96-well plate.
    a. Carefully unseal plates 1-4, still on the magnets.
    b. Transfer eluted sample from plates 1-4 into a new 96-well plate.
    c. Combine rows A-D of plate 1 into row A of the new plate
    d. Combine rows E-H of plate 1 into row B of the new plate.
    e. Continue this pattern with A-D of plate 2 in row C of the new plate, etc, until rows E-H of plate 4 are in row H of the new plate.
2. Further compress 96-well plates 5-8 into new 96-well plate. 8 wells will combine into 1 well of the new plate. Each original 384-well plate will be contained in one row of the 96-well plate.
    a. Transfer eluted sample from plates 5-8 into a new 96-well plate.
    b. Combine rows A-D of plate 5 into row A of the new plate
    c. Combine rows E-H of plate 5 into row B of the new plate.
    d. Continue this pattern with A-D of plate 6 in row C of the new plate, etc, until rows E-H of plate 8 are in row H of the new plate.
3. Compress the two 96-well plates into single new 96-well plate.
    a. Transfer Columns 1-6 of first 96-well plate into Columns 1-6 of a new 96-well plate.
    b. Transfer Columns 7-12 of first 96-well plate into Columns 1-6 of the new 96-well plate to combine with previous transfer.
    c. Transfer Columns 1-6 of second 96-well plate into Columns 7-12 of the new 96-well plate.
    d. Transfer Columns 7-12 of second 96-well plate into Columns 7-12 of the new 96-well plate to combine with previous transfer.

    *Note: final plate configuration will have six wells per original 384-well plate (i.e., wells A1-A6 contain plate 1, wells B1-B6 contain plate 2, etc. for the rest of the left half of the plate. Wells A7-A12 contain plate 9, wells B7-B12 contain plate 10, etc. for the rest of the right half of the plate.*

R. Perform bead cleanup on compressed 96-well plate.
1. Bind free DNA to the Sera-Mag beads
    a. Thoroughly mix Sera-Mag beads by repeatedly inverting bottle and pour beads to the brim of the bead reservoir.
    b. Add 64 µL beads to each well of the sample plate and pipette up and down to mix. Incubate at room temperature for 5 minutes.
    c. Move plate to plate magnet and wait until beads are settled to continue (approximately 3 minutes).
    d. Carefully remove supernatant and discard in waste trough.
2. Wash beads with 80% EtOH
    a. Keeping plates on the magnets, add 180 µL of 80% EtOH to plates 1-4 and incubate at room temperature for at least 30 seconds. Subsequently remove and discard EtOH in the waste trough.
    b. Repeat for a second wash.
    c. Remove plates from the magnets and incubate until beads are dry.
3. Elute DNA

   a. Add 10 µL EB to each well of the sample plate. Set a 5-minute timer.
   b. Seal plate, vortex until beads no longer cling to sides of wells, then quick spin for 5s at 100xg.
   c. After 5-minute timer is complete, unseal plate and place on magnet.
   d. Once beads have settled on the magnet (about 30 seconds), transfer elution to a clean 96-well plate.
   e. Check plate to ensure volume of wells appear even. Seal and quick spin (500 rcf).

**SECTION VII: Adaptase and PCR**

*Steps in this section are automated for 8 or 16 original 384-well plates using Biomek i7

S. Adaptase addition

1. Prepare appropriate volume of Adaptase Mix in a 1.5 mL microcentrifuge tube and keep on ice.

| Reagent | Volume per reaction | Volume for 8 plates | Volume for 16 plates |
|---|---|---|---|
| **Elution Buffer (EB)** | 4.45 µL | 239.2 µL | 478.2 µL |
| **Buffer G1** | 2.00 µL | 112.5 µL | 225.0 µL |
| **Reagent G2** | 2.00 µL | 112.5 µL | 225.0 µL |
| **Reagent G3** | 1.25 µL | 70.3 µL | 140.6 µL |
| **Enzyme G4** | 0.50 µL | 28.13 µL | 56.26 µL |
| **Enzyme G5** | 0.50 µL | 28.13 µL | 56.26 µL |

2. Denature the 96-well plate in a thermocycler at 98 °C for 3 minutes. Immediately place on ice to prevent rehybridization. Plate is ready once bottom of plate is cool to the touch. Quick spin plate 5s at 500 x g before unsealing.
3. Transfer 10.5 µL Adaptase mix to sample plate. Seal, vortex, and quick spin (5s at 500xg).
4. Incubate plate in a thermocycler with the following program:
   a. 37 °C for 30 minutes
   b. 95 °C for 2 minutes
   c. Hold at 4 °C

T. PCR Reaction

1. Add 5 µL PCR primers to sample plate. Be sure each set of 6 wells (the original 384-well plate) gets a uniquely indexed primer.
2. Transfer 25 µL KAPA to each well. Seal, vortex, and quick spin (5s at 500xg).
3. Incubate plate in a thermocycler using the following method:
   a. 95 °C for 2 minutes
   b. 98 °C for 30 seconds
   c. 98 °C for 15 seconds
   d. 64 °C for 30 seconds
   e. 72 °C for 2 minutes
   f. Repeat steps c through e for a total of 15 cycles
   g. 72 °C for 5 minutes
   h. Hold at 4 °C

*NOTE: Plate can be held at 4°C overnight or stored at -20°C for longer times.*

**SECTION VIII: Final Cleanups and Pooling**

U. Perform bead cleanup on PCR plate.
    1. Thoroughly mix Sera-Mag beads by repeatedly inverting bottle and pour beads to the brim of the bead reservoir.
    2. Add 40 µL beads to the sample plate and mix by pipetting. Upon addition of beads, start a 5-minute timer.
    3. When 5-minute timer is complete, move plate onto magnet. Wait until the beads are settled to continue (approximately 3 minutes).
    4. Carefully remove supernatant and discard in the waste trough.
    5. Perform two 180 µL EtOH washes on plate.
    6. After second EtOH wash is complete, remove plate from the magnet.
    7. Once beads are dry, add 20 µL EB to each well of the plate. Start a 5-minute timer.
    8. Seal plate, vortex until beads no longer cling to sides of wells, and quick spin for 5s at 100xg.
    9. After 5-minute timer is complete, place plate onto magnet and carefully unseal.

V. Perform bead cleanup on compressed samples in tubes
    1. Using a 200 µL 8-channel pipette, compress wells such that samples from each original 384-well plate are contained in a single well.
        a. Sample from A1-A6 should end up in A6, sample from B1-B6 should end up in B6, etc.
    2. Using a P-200 pipette, transfer sample from each well into an individually labeled, 1.5 mL microcentrifuge tube (16 tubes for 16 original 384-well plates).
    3. Add 96 µL Sera-Mag beads to each microcentrifuge tube. Set a 5-minute timer, then vortex and quick spin all tubes.
    4. After the 5-minute timer has completed, place each tube onto the DynaMag tube magnet. Allow beads to settle (approximately 3 minutes).
    5. Remove and discard supernatant, careful to not disturb the beads.
    6. Wash beads with 300 µL EtOH. Remove and discard EtOH.
    7. Repeat step 6.
    8. Using a P-10 pipette, remove any residual EtOH at the bottom of the tube and discard.
    9. Remove tubes from DynaMag tube magnet and allow to dry with lid open until beads are visibly cracked and there are no visible droplets of EtOH.
    10. Add 20 µL EB to each tube. Set a 5-minute timer, then vortex and quick spin all tubes.
    11. After 5-minute timer has completed, place each tube onto the DynaMag tube magnet.
    12. Transfer 20 µL from each sample tube into a new clean, labeled, 1.5 mL microcentrifuge tube. Try not to transfer any beads.
        *TIP: It helps to not push the tubes all the way down on the magnet. Slide them in about halfway so the bead pellet stays near the liquid.*

W. DNA Quantification and sample pooling
    1. To quantify the DNA in each sample tube, prepare an appropriate volume of Qubit reagent in a 15 mL microcentrifuge tube according to manufacturer's instructions.
        a. Label one Qubit assay tube per sample. Label two additional tubes for the standards included in the Qubit kit. Also label tubes for any pools to be made.
        b. Combine Qubit reagent and DNA in Qubit assay tubes according to manufacturer's specifications.
    2. Measure the concentrations using the Qubit.
    3. Create sequencing pool by combining normalized amounts of each sample tube.
    4. Check the concentration of the pool using the Qubit. Pool is now ready for sequencing.

**SECTION IX: Primer Information**

**Part I: Primer structure**

Random primer:
/5SpC3/TTCCCTACACGACGCTCTTCCGATCTXXXXXXXX(H1:33340033)(H1)(H1)(H1)(H1)(H1)(H1)(H1)(H1)
Where the XXXXXXXX is an 8-bp barcode sequence, see the barcode sequence and name below.

PCR i5 primer:
AATGATACGGCGACCACCGAGATCTACACXXXXXXXXXXACACTCTTTCCCTACACGACGCTCT
Where the XXXXXXXXXX is a 10-bp barcode sequence, see the barcode sequence and name below. The i5 and i7 primers are in pairs.

PCR i7 primer:
CAAGCAGAAGACGGCATACGAGATXXXXXXXXXXGTGACTGGAGTTCAGACGTGTGCTCTT
Where the XXXXXXXXXX is a 10-bp barcode sequence, see the barcode sequence and name below. The i5 and i7 primers are in pairs.

**Part II: Random Primer Name and Barcode Sequence**

Barcode sequence of random primers:

| | | | | | |
|---|---|---|---|---|---|
| >A1 | GCTACTCT | >E11 | CGTCTTCA | >I21 | AGGTCAAC |
| ACGATCAG | >C7 | AATGACGC | >G17 | GATCCACT | >M3 |
| >A3 | CTCTGGAT | >E13 | TGCGTAAC | >I23 | TACACACG |
| TCGAGAGT | >C9 | TACCGGAT | >G19 | AGCCTATC | >M5 |
| >A5 | AGATCGTC | >E15 | AACACGCT | >K1 | CAAGTCGT |
| CTAGCTCA | >C11 | TTGCAACG | >G21 | AGCTACCA | >M7 |
| >A7 | GCTCAGTT | >E17 | ACTCGATC | >K3 | AGCTAGTG |
| ATCGTCTC | >C13 | CACTTCAC | >G23 | AGATTGCG | >M9 |
| >A9 | GTCCTAAG | >E19 | TGAGCTGT | >K5 | CTCCTAGT |
| TCGACAAG | >C15 | TAGCCATG | >I1 | CACACATC | >M11 |
| >A11 | TATGGCAC | >E21 | TACTGCTC | >K7 | ACTCCTAC |
| CCTTGGAA | >C17 | ACAGGCAT | >I3 | GAGCAATC | >M13 |
| >A13 | TCGGATTC | >E23 | GACGAACT | >K9 | CAATCAGG |
| ATCATGCG | >C19 | AGGTGTTG | >I5 | ATAGAGCG | >M15 |
| >A15 | AACAGCGA | >G1 | CTTCGCAA | >K11 | TCGTGCAT |
| TGTTCCGT | >C21 | CAGTCACA | >I7 | GACCGATA | >M17 |
| >A17 | CCAACGAA | >G3 | ATGGCGAT | >K13 | TAACGTCG |
| ATTAGCCG | >C23 | TCGATGAC | >I9 | CAGACGTT | >M19 |
| >A19 | CAGTGCTT | >G5 | ACATGCCA | >K15 | AAGGCGTA |
| CGATCGAT | >E1 | GAAGTGCT | >I11 | CTGAACGT | >M21 |
| >A21 | GATCAAGG | >G7 | GTCAACAG | >K17 | TCTTACGG |
| GATCTTGC | >E3 | CTTCCTTC | >I13 | TTGGACTG | >M23 |
| >A23 | TCTTCGAC | >G9 | GTGGTATG | >K19 | CGTGTGAT |
| AGGATAGC | >E5 | CGAACAAC | >I15 | GTCTGCAA | >O1 |
| >C1 | ATCGTGGT | >G11 | CCAACTTC | >K21 | AACAGGTG |
| GTAGCGTA | >E7 | AACAACCG | >I17 | CCACATTG | >O3 |
| >C3 | CGGTAATC | >G13 | GACGTCAT | >K23 | AGTCGAAG |
| AGAGTCCA | >E9 | ACCTCAGT | >I19 | GATGGAGT | >O5 |
| >C5 | AGTTGTGC | >G15 | ACGTCCAA | >M1 | TGGAAGCA |

>O7
CTCGTTCT
>O9
ACGAGAAC
>O11
AAGCCTGA
>O13
CTACAAGG
>O15
CGATGTTC
>O17
ACCGGTTA
>O19
GAACGGTT
>O21
CTGTACCA
>O23
GCGCATAT
>A2
TGATAGGC
>A4
CATCCAAG
>A6
GTGAGACT
>A8
CTGATGAG
>A10
ACGGTACA
>A12
CTCGACTT
>A14
ACAACGTG
>A16
TGCTGTGA
>A18
CCAAGTAG
>A20
AACTGAGG
>A22
AGGTAGGA
>A24
TTCGCCAT
>C2
CAGGTAAG
>C4
GTATCGAG
>C6
TTCACGGA
>C8
GAGCTCTA
>C10
GTCAGTCA
>C12
CACGTCTA
>C14
AATTCCGG
>C16
TCTAGGAG
>C18
ATCCGTTG
>C20
GATAGCCA
>C22
TATGACCG
>C24
CGATTGGA
>E2
ACAAGCTC
>E4
GAACCTTC
>E6
AGCGAGAT
>E8
CCGTAACT
>E10
TCAGACAC
>E12
CGAAGTCA
>E14
GTGATCCA
>E16
ACTGGTGT
>E18
CTAACCTG
>E20
AGCCAACT
>E22
CCAGTTGA
>E24
AAGTGCAG
>G2
AACCGTGT
>G4
CGCGTATT
>G6
AGTTCGCA
>G8
TAGTCAGC
>G10
AACACCAC
>G12
GTAAGCAC
>G14
GTCCTTGA
>G16
CAGGTTCA
>G18
CCAACACT
>G20
GAGAGTAC
>G22
AGATACGG
>G24
GTTCTTCG
>I2
ATTCCGCT
>I4
AAGCTCAC
>I6
TGATCACG
>I8
CAATGCGA
>I10
ATGCGTCA
>I12
TACATCGG
>I14
ACTGCGAA
>I16
TCTGTCGT
>I18
CTCAAGCT
>I20
AACCACTC
>I22
CTTACAGC
>I24
AGTCTTGG
>K2
CACGCAAT
>K4
AGCTTCAG
>K6
CCTCGTTA
>K8
TGAGACGA
>K10
CACAGGAA
>K12
ACTCAACG
>K14
AAGCGACT
>K16
CCTACCTA
>K18
ATCTCCTG
>K20
TCACGATG
>K22
CCACAACA
>K24
AGGTCTGT
>M2
AGAAGGAC
>M4
GCGTATCA
>M6
CAACACAG
>M8
TCCACGTT
>M10
ATCGCAAC
>M12
ACGTCGTT
>M14
CGAATACG
>M16
TGCTTGCT
>M18
CTCGAACA
>M20
ACATGGAG
>M22
ACAAGACG
>M24
CGCCTTAT
>O2
AGCAGACA
>O4
GTTAAGCG
>O6
CATGGATC
>O8
ACAGAGGT
>O10
TAAGTGGC
>O12
AGTCAGGT
>O14
GCCTTAAC
>O16
GTTGGCAT
>O18
CAACCTCT
>O20
TGGATGGT
>O22
CTATCCAC
>O24
GATCTCAG
>B1
GAACGAAG
>B3
ACCTAGAC
>B5
TACGACGT
>B7
TTGAGCTC
>B9
AGTACACG
>B11
TGTCAGTG
>B13
GACTACGA
>B15
TTACGTGC
>B17
ACTGCTTG
>B19
GCCTATGT
>B21
GTACCACA
>B23
TAGTGGTG
>D1
ATACGCAG
>D3
AAGACCGT
>D5
CTCCAATC
>D7
TCTGGACA
>D9
AACACTGG
>D11
TTGGTGCA
>D13
CCTGTCAA
>D15
CTATGCCT
>D17
TTCGGCTA
>D19
ACCGACAA
>D21
CGTAGATG
>D23
CTGTATGC
>F1
GTTGCTGT
>F3
AGAACCAG
>F5
GATGTCGA
>F7
AGGAGGTT
>F9
AATCGCTG
>F11
AGTGACCT
>F13
CGAATTGC
>F15
CAAGAAGC
>F17
CACCAGTT
>F19
GTATTCCG
>F21
TTCGAAGC
>F23
AGACCTTG
>H1
CCAAGGTT
>H3
ACGTATGG
>H5
AAGGACCA
>H7
TATGCGGT
>H9
AAGGAAG
G
>H11
AGCGTGTA
>H13
TCTACGCA
>H15
TGGCTCTT
>H17
CCTTCCAT
>H19
ATACTGGC
>H21
AACCTACG
>H23
CATACTCG
>J1
TGCACTTG
>J3
TCACTCGA
>J5
CACTGTAG
>J7
GTACGATC
>J9
TGGTGAAG
>J11

TAGCTGAG
>J13
AGAGCAGA
>J15
CTTCGGTT
>J17
ACAACAGC
>J19
AGCCGTAA
>J21
CTCTTGTC
>J23
CAGATCCT
>L1
GATGCTAC
>L3
AGGAACAC
>L5
ACCATCCT
>L7
GAACGTGA
>L9
TAGAACGC
>L11
AACCAGAG
>L13
CGACCTAA
>L15
CTCTCAGA
>L17
AGGCTGAA
>L19
ATCGGAGA
>L21
GATACCTG
>L23
TCCTGACT
>N1
TCAGCCTT
>N3
AAGCATCG
>N5
GCCAATAC
>N7
GACACAGT
>N9
AAGAGGCA
>N11
GAAGACTG
>N13
CCGTTATG
>N15
CTAGCAGT
>N17
GCCAGAAT
>N19
CGAGAGAA
>N21
AACTCGGA
>N23
ACAGTTCG
>P1
TGACCGTT
>P3
CATCTGCT
>P5
CGCTGATA
>P7
TCGTCTGA
>P9
CACATGGT
>P11
CGAGTTAG
>P13
AGCTAAGC
>P15
GTTCCATG
>P17
GCATCCTA
>P19
CCATGAAC
>P21
ATCCACGA
>P23
GAGAAGGT
>B2
AGGCAATG
>B4
TCACCTAG
>B6
CATACGGA
>B8
GTCATCGT
>B10
TTACCGAC
>B12
ACCTTCGA
>B14
ACGCTTCT
>B16
GAGTAGAG
>B18
ATGCCTAG
>B20
CAACTCCA
>B22
AAGTCCTC
>B24
GTCGATTG
>D2
GCGTTAGA
>D4
TTGCGAGA
>D6
ACACCGAT
>D8
CGTATCTC
>D10
AAGGAGAC
>D12
TGTCGACT
>D14
TCAATCCG
>D16
GACTTGTG
>D18
CCGATGTA
>D20
TAGGAGCT
>D22
CAACGAGT
>D24
TGTGTCAG
>F2
CTAGGTTG
>F4
GTGTCCTT
>F6
TACCTGCA
>F8
CCTTAGGT
>F10
CACAGACT
>F12
TCGAACCT
>F14
GCATAGTC
>F16
CTCCTGAA
>F18
AACGCACA
>F20
TAGTCTCG
>F22
ACTCTGAG
>F24
GTTATGGC
>H2
CTCGGTAA
>H4
TACAGAGC
>H6
GCATAACG
>H8
GATCAGAC
>H10
CGCAACTA
>H12
TCCGATCA
>H14
CAACTTGG
>H16
TCAGTAGG
>H18
ACAGCAAG
>H20
GAATGGCA
>H22
CGGATCAA
>H24
ACTGCACT
>J2
CCTAAGTC
>J4
TTCGTACG
>J6
TCCTGGTA
>J8
CATTGACG
>J10
ACCTCTTC
>J12
CATTCGTC
>J14
TTCCTCCT
>J16
GCTGTAAG
>J18
GACATCTC
>J20
CAACCGTA
>J22
TGCGATAG
>J24
TGGTTCGA
>L2
AAGCGTTC
>L4
CGATTCTG
>L6
GCAACCAT
>L8
AATCCAGC
>L10
AGTGCATC
>L12
GCATTGGT
>L14
CTTAGGAC
>L16
ATAGTCGG
>L18
GAGACCAA
>L20
AACAAGGC
>L22
CCAGTATC
>L24
CCTCGAAT
>N2
CAACTGAC
>N4
TGCTCTAC
>N6
CATCACGT
>N8
GCCACTTA
>N10
GCTTCACA
>N12
ACCGAATG
>N14
CTCACCAA
>N16
CAGAACTG
>N18
AGAAGCCT
>N20
CACGATTC
>N22
AAGCTGGT
>N24
GCAATGAG
>P2
CTCTATCG
>P4
ACTCTCCA
>P6
CAGCATAC
>P8
TACTCCAG
>P10
GAGGCATT
>P12
ACACCTCA
>P14
CGCAATGT
>P16
CCTAGAGA
>P18
TACTAGCG
>P20
CGTCCATT
>P22
TCGCTATC
>P24
AATGGTCG

**Part III: PCR Primer Name and Barcode Sequence**

Format: primer_name: i7_index_sequence, i5_index_sequence

Most frequently used 32 primer names in this study: I3, E5, O5, C7, K9, M9, A11, I15, K15, M17, C21, O21, C23, I2, C10, G10, M14, K18, K20, E22, I24, J3, L3, J7, B11, J11, J2, B6, D6, F6, H6, D8

A1: CTGTTAGCGG, CGTAGAACAG
A3: CGGAAGATAA, CTGGCATATT
A5: ACACTTCGTT, AGAACCTCGC
A7: CGTCGCTCAA, TGTCGTTAAG
A9: ACTTGGCCGA, CATACCGCTG
A11: GATTGATTGC, CTTCAATAGC
A13: TTACTCCGGT, CTCAGTCCGT
A15: CACGCTTGCA, CGCTTACAAT
A17: TAGACAGAAC, AGAGAGACCG
A19: ATTGCTCTTC, TAGCACACTT
A21: GACACACATA, CGGAGCATCT
A23: CTTCAACGGC, CGCTTGGTTC
C1: CCACAGGCTA, CTTGAAGAGG
C3: TGCCGGCATA, AACGCCATTC
C5: ACTAACGTCT, ACAATTGGTC
C7: TTCCGGATTA, TAGTGTTACG
C9: TTCTCGTGTT, CTTCCTAGGC
C11: CTCGAGTTGT, CTTCTCCAGC
C13: TCATGTGCCG, TAGCGCAATA
C15: TGAGAGATAG, TTAGACCACA
C17: CTGGTAACTT, GACCAAGCAT
C19: GCATAAGGTA, TATGCAACTC
C21: CTGGTTAACA, TAACGCAGTC
C23: GCGAGTACAA, CCATAAGCCA
E1: CTTGATGGTT, CGGAACCAGT
E3: TAGTGAAGGA, CGCCATCAAT
E5: CATGGTGTAC, AGGACCTGTA
E7: CGATGCGAGA, CTAGACACTC
E9: TTCTGCCGTA, ACATGGCCTG
E11: TGCGGTATGG, TCCAGCTCAT
E13: CCATGCGACT, CACCAGCACA
E15: TAACAGTGGT, CCGCGATTAG
E17: TCGGTTACTA, AGTGCCGATG
E19: ATAGCGGTGC, GATCTATTGG
E21: TGGTGAGCAA, CATATTGGAG
E23: TTCCGTTCCA, AAGTCCATAG
G1: CCGCTTAATT, AGTACCATGA
G3: GACCTCGTCT, GCCGGTAAGT
G5: AGCCTATTAG, TACCAGAACG
G7: ATCGGTAGGC, CATTGGCCAA
G9: ACCTGATAAG, GCTGGCCAAT
G11: CCTATATGCA, CTACGTGGTA
G13: GATCAGGTGC, CTTATTCGGA
G15: TCACGAGTAC, CTGGCATCCA
G17: TGCATATGTG, CCTAGCAATT
G19: GTATAGTACG, CCTCCTCTTG
G21: TTGGCTGATA, TCGTTGTAGC
G23: GATGGCTTGG, GCAGACTTCA
I1: TGAGGCGATC, CCTTAAGCAG
I3: CCTTCAGAGC, CCTCATAACT
I5: AGAGGCCAAG, TTCATGGTGC
I7: ATTCCGATGA, CGGACGGAAT
I9: TATACGGTGA, CCGTGAGCAA
I11: TGAGGTAACA, TTGGCATCGC
I13: TTAGGCCTTC, GATCTGGTTA
I15: TCGATTCGTA, CCTATAATGC
I17: ACCTCTAACA, GTAAGGAATC
I19: AATGAGGAAC, AAGTAACGTC
I21: CTAATACCAG, TCCAGTTCTC
I23: TTGGAGGCCT, CTGTAGCGCT
K1: TCATAGCCGA, AGACTGATCA
K3: TATGGCTCGC, CGGTTGAACG
K5: CCTATGAAGG, AGAATGGAGG
K7: CGTCTCAAGC, GCGCGAACAT
K9: CATACTCCTC, ACCGGCGAAT
K11: AAGTGTAAGC, GCCAATTGCA
K13: AATCATCGAC, CCTGCAGCAT
K15: TGACTCGCGT, CCAGAGGAAT
K17: TTCTTCACAG, CCAATCAAGT
K19: CCGTTGCGTA, CACGTTACTC
K21: ACGCAGTGGA, CTACGTTCCA
K23: CCAGGCTCAA, TAAGGTGACA
M1: CGATTCACTA, AAGCAGGCTC
M3: CAGTACAGGC, GCTACACGAG
M5: AGCTTATCAC, TGCTATGCAG
M7: TTCGTGGTAG, ACCGCTAGCA
M9: TCGTATATGG, AGACAGTGGC
M11: ATGGCCGTTC, GTGCAGATTA
M13: TTCACGTACT, CTTACGAGTA
M15: AGTTCCGAGC, CCTCACAGAT
M17: TCTATGTTGG, CCAACAGATT
M19: TGCCACACTG, TAGAGTGAGA
M21: TCCGCTTCGA, TAAGGTTCGT
M23: ACCGTAGTGC, TCGTTCGTAA
O1: TATAGTCGAC, TCAGCAGAGG
O3: TCGTACTTGC, GCTAACGGCA
O5: AGACAACACT, ACGCAAGAAG
O7: TAGGCAACCG, TAGAGGCTTA
O9: TTACCATTCG, AGTAGAGTGT
O11: CCTATTGCGG, TCCTTGTCGG
O13: CTACGCACAG, CACTAATCAG
O15: TGCGCCACAA, AACCGAGATT
O17: AACTTGCCTT, TACACGAAGA
O19: CCATCTAGAA, AGCGACAGGT
O21: ATTGTTCCAC, GAGTAACCTT
O23: CCTCTCCACA, ACGGCTAGAG
A2: GACTGCAGCA, CTACCGCGTT
A4: ACTGTAGCTG, GCAAGAGATG
A6: GCAGATAATC, TCCTACAGCG
A8: TGCGCTCATG, CCAGTGACGA
A10: AGGCCTAAGA, GTGGCGGTTA
A12: TAATCCGCTT, AGACACTCTA
A14: TTCTTAGTCG, CCAAGCACAA
A16: TCAGTTGTTG, TATGTGCGCT
A18: TTCCTTCGAG, CTGTTAGGAT
A20: AGTAAGCTTC, CGCATCTTGA
A22: AAGGCGCCAT, AGAGGAGCAG
A24: CCTAATAGGA, AGGTCGATAA
C2: TCGTGATTGT, AGCGTAGCAA
C4: CTGGTGGTCA, CTCAACGGAG
C6: AGCCTACGTT, CTGTGCATTA
C8: AACCTTCAAG, GTGCGTCGTA
C10: TTCACGTCAA, CCATCCGAGT
C12: TTGGCGTTGT, CGCCACAGAA
C14: TCTAGACCGG, GCATCTTCTT
C16: CCTCATAATC, AAGATACAGC
C18: CCTCACAGAG, TTCCTAGCCG
C20: TCTTCACCTT, ACCTTCCGGT
C22: TATACCTCTC, TTACTAGGCA
C24: GCAGTTAACA, GAGTGTCTCA
E2: TTATCGCGTA, CGCTACACTC
E4: TATAGCCAGG, TCGGATGAGG
E6: CTTATCCGTT, GAGGCCACAT
E8: AGATAGGTAG, CCATAAGAGG
E10: CTTAATACGG, TTGGCTAGGC
E12: GCCGGTGTTA, TTAGAATCGG
E14: CATATGGCCG, GAACGGACTA
E16: CCGGATAAGT, TAATATCCGC
E18: TTGAGACCAG, GCATTCAGCG
E20: TCCGGTGTGT, AGGCCTTATA
E22: GATGTTCATC, AACGGATCGC
E24: TGGCTAGGTT, TACATGCCGT
G2: TACTATTCCG, CTAGATAACG
G4: CACGGTATAA, AGAACCGCTA
G6: ATCTGCAAGA, TTGTCTGTAG
G8: ACTAGCACGC, CGCTCGTGTT
G10: CTCGAATGCG, GATCATTCCT
G12: AGCTGGTAGG, CTTGAATTCG
G14: TAGGATCAAG, CCGTGTTCTC
G16: TCGGTCGATG, AGGCAGCTTA
G18: TTCCTCGGAC, TGACCTGCTG
G20: AAGTGTTACG, CCTATTCAGT
G22: CGAGAGGTGA, GTGTACACTA
G24: AGAGCCTATT, TCGCACTCCA
I2: TAGTTCCATC, CTGTGAATGT
I4: CCACTTCGGA, AGGTTACCTT
I6: GAAGTCATAC, TTGCCATCAG
I8: TCATCTTGCA, TTGTAGGACA
I10: AGTTCTTGAC, GCTACTTGAA
I12: AACGGATTAG, ACGCAAGTTC
I14: AGAGAGACAA, CCTGCTAGGA
I16: TCAACGGTAG, TGCGGACACA
I18: CCAATTCACA, AGGACACATA
I20: TCTGTGCCAC, GCAAGTATTG
I22: CTAGCGGTTA, TGAATCGCGT
I24: CCGAATTGGT, CCAATCCTAA
K2: TTCACATGAC, TTGGCTGCGT
K4: CCATGACGCA, GAGACAAGGA
K6: TCGGCCTATA, CCGGCCATTA
K8: TCCTTGATGG, CCGTAGTTGG
K10: AATCTGCGGC, GTGCAGTAAT
K12: ACTTCAGGCA, TAACTCAGCA
K14: AGAGACTCGG, TTCTGTTCTG
K16: ATGGCGTTAG, GCTTGTGCCA
K18: AAGAGTGTGA, CGATAACCAA
K20: CCTTAATTGC, GATATACCTC
K22: ACGGTTGGAC, CACCAATAGT
K24: AACTTGTTGC, TTCGTCAACT
M2: TGTGATGGCA, CGTGTTCTTA
M4: TCTGCACACT, TCACAGCTGC
M6: AGTCGAATTC, TGAACCACAA

M8: TCAGGACTCG, CCGCTAGCTT
M10: AGGTTGCTAG, GCCTAAGAAG
M12: AGTTATCCTC, TGTTGCACTA
M14: CTCATCTTAC, CCAGCTATAA
M16: TTAGCTGGCA, TGGCATCTGG
M18: TACTTCTGGT, GTCTTATCAG
M20: TAAGTGTGAG, CAAGAGGAGG
M22: ACTGGCGCTA, GCAATCGCAA
M24: TAGCTCTTAG, TCGGTGAATA
O2: AGCTCTGGCA, CGTAACATAG
O4: ACGAATAGTC, AAGCTAATGG
O6: ATGTTACTGC, CAATACCTTC
O8: TTCATGGTGT, CTCATGATGA
O10: AGTGCTAACG, AACGAGCCTC
O12: ATGCAGTTAC, ATGACTGCCG
O14: GAGTTGCCAG, GACATGAGTT
O16: CATACTAAGC, CTTAGTTGCA
O18: GACGACTAAC, CAGGCGTCAA
O20: GCTTATCGTA, GAAGCAACAA
O22: GAAGTAGCCG, TAACTCGTGG
O24: GCACTGCTGT, GACATGTAGA
B1: AACGCGGTAG, TGCTGCAGTC
B3: CATCTCTTGC, CAGCTACAGT
B5: CGCTGTAGGA, GATTATCTGC
B7: TCGTCACTGG, CATGAGCGCA
B9: TAACCGCCAC, TCTTAGGCCT
B11: TAGAGTGTCT, AAGCGGATTA
B13: TTGTGCTTGG, CGACTAAGAA
B15: AGCGCACATA, CAACAACTGA
B17: ATCCTAACAG, CTCGAAGGAA
B19: TCAAGATGCG, GAAGCTTACT
B21: CTGCTCCTCT, ATGGCGCCTT
B23: TTATCGACCT, TCCTATTAGG
D1: TTGCTACGCT, ACAATCACGA
D3: CTCCGTTGAG, TGACCACCAG
D5: TAATGCACAG, AACGTAGGCT
D7: TACGACGCAC, CTTGAAGGTT
D9: ACTCGGATGG, TTGACGTGAA
D11: TTCTGTGGCG, ACAACGCCAA
D13: AAGAAGATGC, CCGGTCTAGA
D15: GCTGTATCTT, GATTATGAGG
D17: CGGCTAGGAA, CTCTGTGAGG
D19: ACCGGAAGGT, AAGGTGAAGA
D21: TGCCTAGTAA, GAGAGGTATA
D23: TGAGACACTC, TGTTAACTGC
F1: GAGTGTAGCG, TACGCGATAA
F3: CCTCATCCGA, CCTGGCTATA
F5: ATGTGGCCTC, AACGGATAAG
F7: CCTCTTATGG, CTACCTATCT
F9: GCCTAGCCAA, CCGTATTAAG
F11: CCGATTCTAA, TAACACCGGC
F13: TAGTCCGTTC, CGGCCATATG
F15: GCGGATATTA, ACTTATCCGG
F17: CGCTGAATGC, CTGGTCTCAA
F19: TATAAGGCCT, ACACACCGGA
F21: GCGATCCGTT, GATGAACATC
F23: ACGGCATGTA, AACCTAGCCA
H1: CGTTATCTAG, CGGAATAGTA
H3: TAGCGGTTCT, TTATACCGTG
H5: CTACAGACAA, TCTTGCAGAT
H7: AACACGAGCG, GCGTGCTAGT
H9: AGGAATGATC, CGCATTCGAG
H11: CGAATTCAAG, CCTACCAGCT
H13: GAGAACACGG, CTTGATCCTA
H15: TAAGCTGCCT, CATCGACCGA
H17: CAGCAGGTCA, GTCCGAGGAA
H19: ACTGAATAGG, CGTAACACTT
H21: TAGTGTACAC, TCACCTCTCG
H23: TGGAGTGCGA, AATAGGCTCT
J1: ACATTCACAG, GATGGAACTA
J3: AAGGTAACCT, TCCGAAGTGG
J5: CTGATGGCAA, GTATGACTTC
J7: TGTCCTACAA, TGCAAGATGA
J9: TTCAAGTAGC, GTCAAGAACT
J11: GAACTTGCGT, CTAATCCGTT
J13: TCCTAGCAGG, TTGTCTCTCT
J15: TGTGTCCGCA, CTACCGTTGA
J17: TATGTGTCCT, TGTGAATTGG
J19: CAATACTTGC, GTGGCACAGA
J21: ACGCGATTCA, TAACCGCTAG
J23: TTAGGATTGG, ACCAATTCGG
L1: ACGCAGCCAA, GTCATGTGAA
L3: TCCTTGTCTC, TGCGTCATGG
L5: TAATGGCCGG, TATAGGCCGA
L7: CCAACTACGG, CCATCAAGGA
L9: ATTACTGCAC, GCCGCAGATT
L11: TGCTGAGTTA, TGCCTGAAGT
L13: CAGAACAGAA, CCGAGTCTCT
L15: TGGCACAAGC, CTAACGCCAT
L17: TTGGTTATCG, TCACACTCTT
L19: GAGGTATATC, GCAATTAAGG
L21: ACTATTGGTG, GTCCAACCGT
L23: AGTTGACGAA, GCAACAAGTT
N1: TAAGAACACG, TGCCATCACA
N3: GCAGCTGTGA, AGTGTGCAGA
N5: TTGTGGTTCA, GAATTCGACT
N7: AAGCTAGCGG, CGAGTCCTGA
N9: CTTCTTAGGC, CTAGCAACCT
N11: TAGTGCAACA, GAGGATAACT
N13: TTATAGCTGG, GTAAGATGAG
N15: CCAGATGCCA, TGCCAGCTAA
N17: CTGATAAGAC, ACCAGAAGTA
N19: CCTCCTCTTG, CTCACACTTA
N21: TTGCGATTGC, TAGCGCCAGT
N23: TATTCACCGA, CTAAGAGCTA
P1: CTATGTTACG, TGCCAGAGCT
P3: CCATTAGCTT, GACTATTCGT
P5: GAAGGTATTG, GCAGTAACAT
P7: TCATCATGAG, ACACCATGAA
P9: GAGGCTCGTT, CGTTAGCACT
P11: CGGCAGTCAT, GTAACTGCAT
P13: AACTCATGTC, CTGGCAACTC
P15: TGCAACTAAG, GCTTAGTATG
P17: TTGACGCCTG, GTTAGTCGTC
P19: TTGTTGCTTC, TACGATAAGC
P21: CCACGAGTTA, CGGCTACTTC
P23: TCTACATGTC, ACAGCAGTGC
B2: CTGTTCTACG, CCGCTAACAG
B4: AATATGCCAG, TTATCTTCCG
B6: GCGAGGTTCT, AACGAAGTGT
B8: CTTAACGACA, TTGAGCGACG
B10: CAGCGGTATG, TCGGCCAAGT
B12: GCTATTGAAG, GCAATCAATC
B14: ACGGACTGAG, ACCGGAGTAA
B16: ATTGTAAGCG, TGCAAGCGTG
B18: CGGTCTCTCT, GTTCTGTCTA
B20: AAGTGTGCTA, GAAGAGCAAT
B22: AGATGCTCCG, TATGTGTGTC
B24: GAACCAAGCG, GCCGTTGAAG
D2: CCTCTTCAAG, TAGCCTGTGG
D4: GAATGGCGTT, TATGCCGGCA
D6: GACCAATTGT, AGACGTTAGT
D8: CGTAACACTA, TAATCCGGAA
D10: GCCTAGGAAG, AACACGAGAA
D12: GCTGGAGAAG, ACAACTCGAG
D14: TATTGCGCTA, CGGCACATGA
D16: TGTGGTCTAA, CTATCTCTCA
D18: ATGCTTGGTC, AAGTTACCAG
D20: GAGTTGCATA, TACCTTATGC
D22: GACTGCGTTA, TGTTAACCAG
D24: TGGCTTATGG, TTGTACTCGC
F2: ACTGGTTCCG, AACCATCAAG
F4: ATTGATGGCG, TCCTTCACTA
F6: TACAGGTCCT, GTACACCATG
F8: GAGTGTCTAG, TCTCGCATCG
F10: CAGGCCATGT, TACGGCAGAA
F12: ATGAGCTGGA, CCATACCGCA
F14: TGTGCTGGTG, AGTCGCATGG
F16: CTAATCGCGG, ACCACTGTTA
F18: CATCGGCACT, TAGTAACCGA
F20: CCAATAGATC, GCACCGCTAT
F22: CTCCAATATG, TTGCTCACCA
F24: CTATGGACTT, TGGAACGGAA
H2: TCATGGTACT, AATTAAGCGG
H4: ACTTACCGGC, AGACGAGGTC
H6: CGTTCTGGTA, CTAATAGGCT
H8: TTGGCCAATG, GCCTACCGAT
H10: ATTGGCCAGC, CTTATCAGGT
H12: TACCACGTAG, TGCATATAGG
H14: TCCGAATAAG, GCACCTGATC
H16: TGGATGCCAG, GTACTCGTGA
H18: AATTGCTAGG, CACATATGCA
H20: CAAGAGGAGG, CGTACTATAC
H22: GCTACAACGA, TATCAGCCAA
H24: TGAAGTCTGC, CCAAGCCATC
J2: CTGCTTAAGG, GATCGCCTCA
J4: AGTTATGAGG, GCTCTGAAGG
J6: GCACCATGAA, CTTGGCCTCT
J8: ATTCCGTCCG, TCATCGGAAT
J10: TTGCTCACGG, TCACCGTATA
J12: GCGATGCCAA, TGTTACCTCA
J14: TAACCAGATC, GAAGGCCTAA
J16: GCATTATAGG, TACGAATCGA
J18: GATTCCTTAC, TGTTAGAGGT

J20: GACGTTACTT, GTTCCTCATT
J22: GAGAACTGGA, CTGGTATTAG
J24: AGCGCTACAG, AGGCCTCCAA
L2: TGATCAGTCT, TCGGCTATGA
L4: CGTTCAACCG, GCGAGCCATA
L6: CCTCCATTCT, CCTTCATAGG
L8: ATGTTCGCGC, GCTTGAGACG
L10: ATTCGCCGGT, GAGGAGTATG
L12: TGCAATTGGC, GCTTACACTT
L14: ATGCTGCAGG, GTCGATGATT
L16: ATTCCTCTGG, ACGCGAGTCA
L18: ACTTGATTGG, CTGTGAAGAA
L20: GAGTAACGTG, TACGCAACGG
L22: TGGAACGTAG, TCCACTGCGT
L24: TGTCACCTTA, TTGAGCCTGG
N2: GAGCCTGCTT, TAGTGAATCG
N4: CTCGTGTAGC, GCCTGTACTG
N6: CTGCATAGCA, GTGATAAGCT
N8: TGCTAGCGGT, CTACCACGAA
N10: GCCACTGTCT, CCATATACGA
N12: TAATCTGCAC, GAACGGCCAT
N14: TACTCGTAAG, AGTACGTGAA
N16: ATCTGTGAGG, GCTCGGAACT
N18: AATCTGTTGG, CCAACATAGA
N20: TCTCACTCTA, CAGTGTGGCA
N22: ACGAACCTTA, TCGAAGCGGA
N24: TTACGAACGA, GCACTACGGT
P2: CCTCTGCTGA, GTCGACTATA
P4: TGCCGTTAGC, GCAAGTACGA
P6: CTTCTTGCGT, AGTGTTGTCT
P8: TAAGCCTCTA, CGGTTGCCTA
P10: ACACTCTACT, CGAATGGTAA
P12: CCGACAAGGA, CCGCAATAGG
P14: CTGATTAGTG, CTGTGCGTAG
P16: AATCTCGGTT, TTGTGGCGCA
P18: TCTTCGTGTA, AAGGCAAGTT
P20: ACCTGTCGCT, TTCTAGATGG
P22: AAGGTTACTC, GTGGAACAAT
P24: CTCTAGCCGT, TGTGGAGAGG
