## Supplementary Information 2 for "Single-cell DNA Methylome and 3D Multi-omic Atlas of the Adult Mouse Brain"

### snm3C-seq FANS gating example

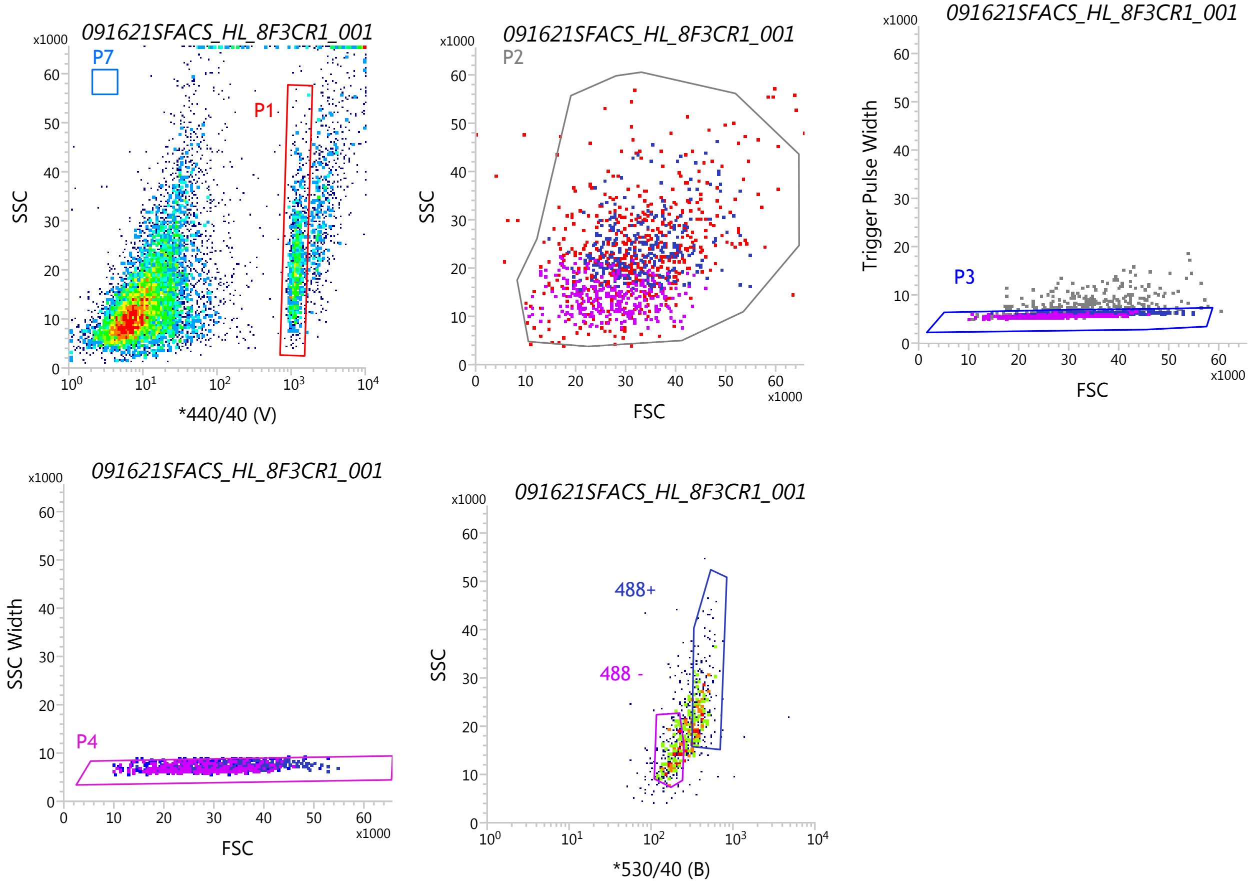


Statistics: *091621SFACS_HL_8F3CR1_001*

| Populations | Events | % Total | % Parent |
|---|---|---|---|
| All Events | 10,140 | 100.00% | #### |
| P1 | 1,049 | 10.35% | 10.35% |
| P2 | 1,035 | 10.21% | 98.67% |
| P3 | 803 | 7.92% | 77.58% |
| P4 | 752 | 7.42% | 93.65% |
| 488+ | 277 | 2.73% | 36.84% |
| 488 - | 267 | 2.63% | 35.51% |

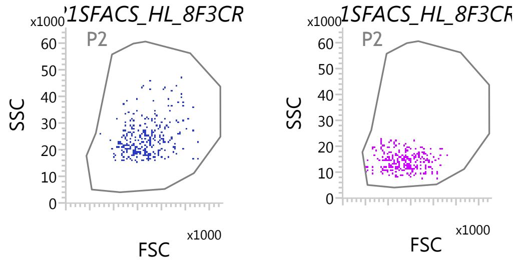

### snmC-seq3 FANS gating example

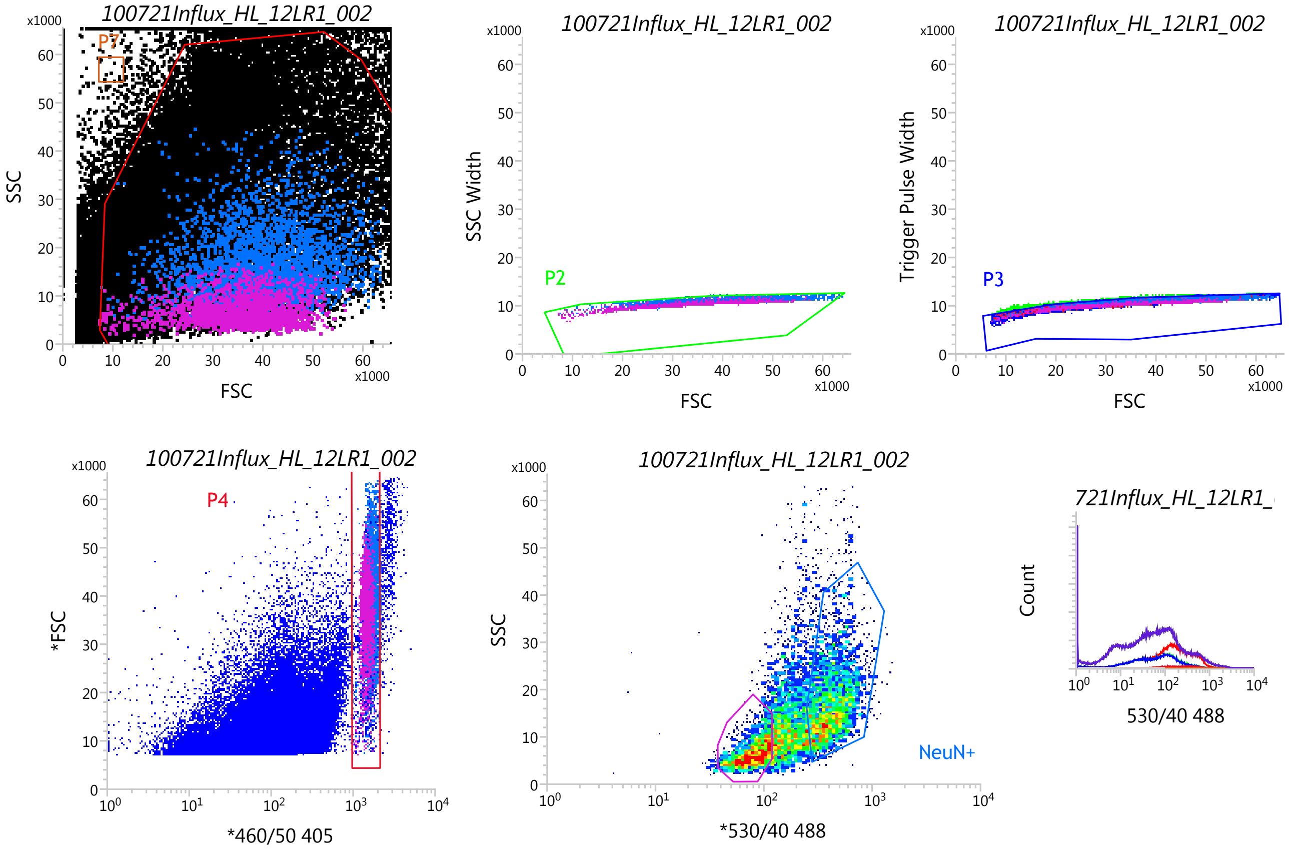


Populations: *100721Influx_HL_12LR1_002*

| Populations | Events | % Total | % Parent |
|---|---|---|---|
| All Events | 284,075 | 100.00% | #### |
| P1 | 116,722 | 41.09% | 41.09% |
| P2 | 69,020 | 24.30% | 59.13% |
| P3 | 66,582 | 23.44% | 96.47% |
| P4 | 7,324 | 2.58% | 11.00% |
| NeuN+ | 2,930 | 1.03% | 40.01% |
| Neg | 1,901 | 0.67% | 25.96% |
| P7 | 17 | 0.01% | 0.01% |
| NOT(P7) | 284,058 | 99.99% | 99.99% |

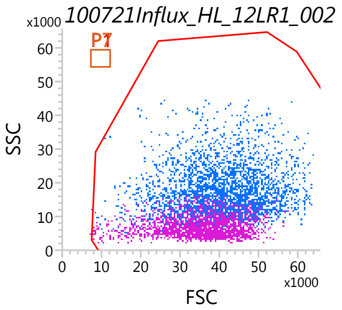
